# Application of modular isoxazoline-β^2,2^-amino acid-based peptidomimetic foldamers as a chemical model system for studying the tau misfolding mechanism

**DOI:** 10.1101/2024.10.04.616679

**Authors:** Davide di Lorenzo, Nicolo Bisi, Raffaella Bucci, Inga Ennen, Leonardo Lo Presti, Veronica Dodero, Roland Brandt, Sandrine Ongeri, Maria-Luisa Gelmi, Nicolo Tonali

## Abstract

Tauopathies are a group of neurodegenerative diseases characterized by the deposition of abnormal aggregates of tau protein into the brain. Tau is a microtubule-associated protein whose physiological role is related to the modulation of microtubule dynamics and the correct axonal transport in neurons. In tauopathies, the mechanism of conversion of the tau molecule from a physiological to a pathological form is still unclear. Nevertheless, two hexapeptide sequences, named PHF6 and PHF6*, located in the R3 and R2 repetition domains of tau, respectively, seem to be crucial in triggering the aggregation. In the native state, the hydrophobic residues of PHF6 and PHF6* are protected by a β-hairpin-like structure. Contrary to that, the misfolded protein exposes hydrophobic residues, which can form hydrophobic interactions and drive the self-assembly process. Chemical model systems, other than being more straightforward and easily accessible than full-length proteins, can provide valuable insight, which is difficult to tease out by studying the natural systems directly, and can especially be employed to examine the structures and modes of folding of amyloid proteins. Here, we exploit the use of chemical systems to modulate and study the mechanism of tau misfolding and aggregation. By employing a new non-natural β^2,2^-amino acid, we have induced the two hot spot sequences of tau (PHF6 and PHF6*) either to have a totally extended conformation or a β-hairpin, according to the *(S)* or *(R)* stereochemistry of the scaffold, respectively. The aim of this work was to provide, through the interchange between β-hairpin-like and extended conformations, a possible explanation for the mechanism of tau misfolding. We demonstrated that an extended conformation, exposing the hydrophobic residues of both sequences and driving away PHF6* from PHF6, can trigger the aggregation of tau and behaves as a seed-competent monomer model system. Conversely, a β-hairpin imitates the favorable folding of tau, allowing the protein to maintain its soluble monomeric form. Furthermore, a β-hairpin mimic, based on the chaperone protein Hsp90, demonstrated the further application of this type of peptidomimetic foldamers as both aggregation inhibitors and chaperone mimics for the corrective folding of tau in a neuronal environment.

## Introduction

Tauopathies are a group of neurodegenerative diseases characterized by the deposition of abnormal aggregates of tau protein in the brain, forming neurofibrillary tangles (NFTs) inside the neuronal cells.^1^ Tau is a microtubule (MT)-associated protein (MAP) whose physiological role is to regulate MT polymerization and ensuring the correct axonal transport in neurons.^2^ Tau interacts with MTs through the microtubule-binding domain (MTBD)(Q244-K370), that differs, among the various tau isoforms, in terms of the repeat domain numbers (three, 3R, to four, 4R).^3^ The 4R isoform was shown to promote microtubule assembly and binding to microtubules better than the 3R isoform.^4^ However, in tauopathies, tau undergoes a conformational transition from a physiological to a pathological folding state, the mechanism of which is still unclear.^5^ It is widely accepted that conformational or post-translational changes in the protein must initiate the autocatalytic aggregation. Furthermore, this self-assembly, associated with abnormal phosphorylation, induces the detachment from MTs and the formation of intracellular aggregates.^6^ This process is structurally unique for each tau strain and tauopathy, resulting in a different panel of fibril structures and polymorphs.^7,8^

Morphologically, NTFs can be composed of two types of fibrils: paired helical filaments (PHFs) and straight filaments (SFs), whose common structural core consists of residues V306-F378, indicating the inclusion of R3 and R4 repeats.^7^ R2 has been demonstrated not to be part of this pronase-resistant core, as immunolabelling of PHFs and SFs from brain patients with an R2-specific epitope antibody (anti-4R, raised against V275-C291 of R2) after pronase treatment, was negative.^7^ It is evident that the MTBD plays a central role in tau protein aggregation. Particularly, tau aggregation seems to be mediated by short motifs, called amyloid motifs, corresponding to highly hydrophobic regions of the full sequence. Two hexapeptide sequences, named PHF6 and PHF6*, which are located in the R3 and R2 repetition domains, respectively, seem to be crucial in triggering the aggregation (Figure 1A-B). The hexapeptide PHF6 (^306^VQIVYK^311^) has been confirmed, by cryo-EM analysis, to be present in the core of Alzheimer’s disease filaments, packing inside the in-register cross β-sheet structure through a heterotypic, non-staggered interface with the opposing residues, ^373^THKLTF^378^.^6^ Recently, cryo-EM structures of fibrils of VQIINK (PHF6*) allowed Seidler *et al*. to hypothesize that the core of the fibrils (PHF6) might be not the primary driver of aggregation, but might serve as a solvent-excluded scaffold that can cluster PHF6* together in the fuzzy coat, which poises the solvent-exposed VQIINK steric zippers for seeding.^9^

**Figure 1.**
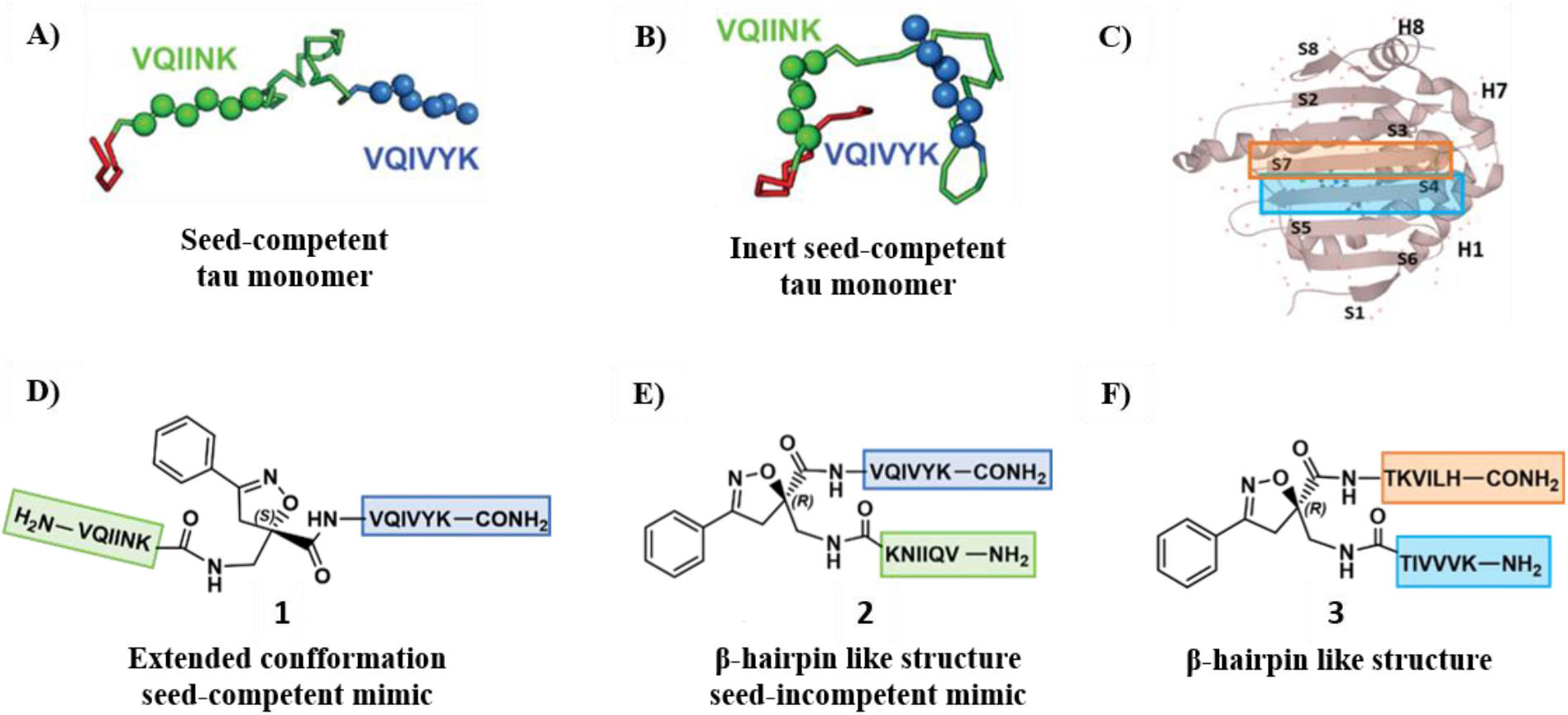
Schematic representation of the designed compounds **1, 2** and **3**: A) conformation of tau monomer in the seed component organization and B) conformation of tau monomer in the inert seed-component organization, as reported by Mirbaha et al.^12^; highlighted in green PHF6*, (VQIINK) and blue PHF6, (VQIVYK). The two Rosetta structural models of discrete tau domains are taken from the original paper. C) Three-dimensional conformation of the crystal structure of the N-terminal portion of Hsp90 (PDB = 3NMQ). The S4 and S7 β-sheet in an antiparallel position are highlighted in blue and orange, respectively; D) Chemical structure of extended peptide **1** bearing the S-configured Isox-β^2,2^-AA which is based on seed-competent tau monomer shown in A; E) Chemical structure of β-hairpin peptide **2** bearing the R-configured Isox-β^2,2^-AA which is based on inert seed-competent tau monomer shown in B and according to the hairpin shape of Kadavath et al.^3^; F) Chemical structure of β-hairpin peptide **3** bearing the R-configured Isox-β^2,2^-AA which is based on the N-terminal portion core of Hsp90 (PDB = 3NMQ) shown in C.

Tau is considered an intrinsically disorder protein (IDP), containing a high proportion of polar and charged amino acids, which results in the absence of a well-defined three-dimensional structure.^10^ Kadavath *et al*. showed, through NMR analysis, that, although tau is highly flexible in solution and can form β-sheet structure only in amyloid fibrils, it can locally fold into a stable structure upon binding to MTs. This conformation is characterized by a hairpin shape, involving the hexapeptides PHF6 and PHF6* located at the beginning of R3 and R2, respectively.^3^ This observation indicates that, when tau plays its physiological role, the two amyloid “hot spots” adopt a favorable conformation, allowing the protein to maintain its soluble monomeric form and thus its MT interaction. Chen *et al*. supported this hypothesis by showing that tau adopts a β-hairpin-like compact structure that shields the ^306^VQIVYK^311^ motif and mitigates aggregation. They also claimed that disease-associated mutations close to this motif may contribute to tau molecular rearrangement from an inert to an early seed-competent form.^11^ By using cross-linking reaction and mass spectrometry, Mirbaha *et al*. demonstrated that tau monomer could exist as inert and seed-competent stable monomer states, and it is only when tau assumes an open configuration, thus making the two ^306^VQIVYK^311^ and ^275^VQIINK^280^ sequences more accessible, that tau monomer produces larger seed-competent assemblies.^12^ By combining all of these observations, we envisaged that PHF6 is the key amyloid motif, which triggers the aggregation propensity of tau by exposing its hydrophobic character in an extended conformation manner. Contrary to the native state, where hydrophobic residues of PHF6 are protected by a β-hairpin-like compact structure, in which PHF6 can directly interact with PHF6*, the misfolded protein exposes hydrophobic residues which can form hydrophobic interactions, driving the self-assembly. The removal of PHF6 from PHF6* might explain why R2 is not involved in the formation of the NFT core. Moreover, 3R-tau has a higher propensity to form dimers and HMW oligomers than 4R-tau isoforms, and that the absence of the second repeat promotes the aggregation.^13^

Our understanding of tau aggregation has come mostly from *in vitro* studies using either tau models, in which exogenous additives are required to initiate tau aggregation, or tau fragments, which can spontaneously aggregate.^6^ T40 (2N4R) full-length tau has been extensively used to study the aggregation process.^14^ The disadvantage of this model is the requirement of heparin, RNA, or fatty acids to induce self-aggregation, leading to the formation of filaments that differ from the core of AD filaments. Because of their main involvement in the NFTs core formation, tau fragments that only comprise the MTBD (K18/K19) represented an easy and common tau model for studying the entire aggregation process. On the other hand, this approach suffers from the same problem of structural dissimilarities of the filaments produced, especially regarding the packing of PHF6 hexapeptide.^5^ To date, the dGAE, fragment (residues 297-391), corresponding to one of the species isolated from the proteolytically stable AD PHF core preparations,^15^ is considered a consistent and reliable model for understanding pathological tau self-assembly and for screening inhibitors, thanks to its ability to self-assemble spontaneously without cofactors. However, although the tau model evolved and arrived to reproduce the correct packing of the NFTs core, the disease-specific conformations and the mechanism of the misfolding transition remain unknown. As a result, other models able to mimic the conformational change of tau are needed to improve the knowledge of the local and global structural changes in tau, which are responsible for its capacity to form a seed-competent conformation.

Chemical model systems that are simpler and easier accessible than full-length amyloid proteins but inspired by their sequences can be designed to behave in controlled fashions to form well-defined conformations. These model systems can afford structures at atomic resolution, providing insights into the structures and modes of folding of amyloid proteins. Such model systems have been, for example, largely employed by Nowick *et al*. to elucidate the structures of amyloid oligomers. These oligomers are formed by macrocyclic β-hairpin mimics derived from Aβ, α-synuclein, and β2-microglobulin.^16,17^ The macrocyclic β-hairpin peptides consist of two peptide β-strands from the amyloidogenic peptide or protein that are constrained by two turn units of ^δ^Orn.

Peptidomimetic foldamers are synthetic molecules that mimic the structure of proteins. They have been developed primarily to mimic the 3D structures and functions of biopolymers, particularly nucleic acids and proteins, through a biomimetic approach. In fact, they are characterized by unique conformations that are inspired by the structural features of their bioactive peptide counterparts.^18–25^ Several examples of peptidomimetic foldamers can be found in the literature in a wide range of applications, such as catalysis, medicinal chemistry, and materials science.^26–28^ To our knowledge, no examples of foldamers application in chemical biology, as chemical model systems for the understanding of amyloid aggregation process, are present in literature. This article presents the application of modular isoxazoline-β^2,2^-amino acid (Isox-β^2,2^-AA) based peptidomimetic foldamers as chemical model system for studying the tau misfolding mechanism. The two “hot spot” sequences of tau (PHF6 and PHF6*) have been forced to adopt either a totally extended conformation or a β-hairpin, according to the *(S)* or *(R)* stereochemistry of the scaffold, respectively. It is here demonstrated that an extended conformation, exposing the hydrophobic residues of both sequences and driving away PHF6* from PHF6, can trigger the aggregation of tau and behaves as a seed-competent model system. Furthermore, the insertion of the *(R)*-(Isox-β^2,2^-AA) into peptide sequences based on the Hsp90 chaperone protein demonstrated to be a valuable way to rationally design stable β-hairpin compounds, acting as potent modulator of tau aggregation.

## Results and Discussion

### Design

Recently, some of us showed that the non-coded Isox-β^2,2^-AA can promote an α-turn or extended conformations when inserted into tripeptide sequences, depending on the absolute *R*- or *S*-configuration at isooxazoline C-5.^29^ Based on these results, we decided to employ this Isox-β^2,2^-AA, to assess its potential as a modulable inducer of secondary structure into peptide sequences, able to stabilize either an extended or a β-hairpin conformation, accordingly to the C-5 stereocenter. Although β-amino acids have been extensively studied and employed to stabilize specific conformation,^22–26^ there are still no examples of a single β^2,2^-AA demonstrating the ability to induce β-turn folding in long peptide sequences.

As previously mentioned, the two hexapeptide sequences of tau, known as PHF6* (^275^VQIINK^280^, 2nd repeat (R2)) and PHF6 (^306^VQIVYK^311^, 3rd repeat (R3)) either are involved in a hairpin-like conformation, when tau physiologically interacts with MTs,^3^ or they are the “hot spot” sequences triggering tau oligomerization and aggregation in pathological conditions.^7,9,12,30,31^ The inert seed competent tau monomer structural model reported by Mirbaha *et al*., ^12^ masking of VQIINK and VQIVYK sequences in compact ‘hairpin’ structures (Figure 1B), is similar to the structure of microtubule-bound tau previously determined by Kadavath *et al*.^3^ Despite different studies supporting the hypothesis that the extended conformation of these sequences promotes the pathological aggregation of tau,^12,32^ a mechanistic explanation is still lacking. Therefore, with the objective to build two original chemical model systems mimicking the seed-incompetent and seed-competent monomer conformation of tau, as reported in Figure 1, Isox-β^2,2^-AA was used as a tool to build extended or β-hairpin conformations when inserted in peptide sequences. We linked to the *R-* and *S-* scaffold the two hexapeptide motifs PHF6* and PHF6 at *N-* and *C*-termini, respectively, to better understand if a conformational change from a β-hairpin (compound **2)** toward an extended structure (compound **1**) might trigger the wild type-(wt-) tau aggregation (Figure 1 A and B, D and E).

Finally, to verify the ability of Isox-β^2,2^-AA to induce more stable β-turn like folding in a long peptide sequence and to behave as synthetic chaperone modulator of tau aggregation in β-hairpin conformation, as previously described by some of us ^33^, the S4 and S7 peptide sequences of the *N*-terminal portion core of Hsp90^34^ were attached to the *R*-configured Isox-β^2,2^-AA (Figure 1, C and F).

### Synthesis

Isoxazoline intermediate **9** was synthesized through 1,3-dipolar cycloaddition between the azido substituted dipolarophile **5** and nitrile oxide **8** (Scheme 1, A). The commercially available bromo-methacrylate **4** was first subjected to a nucleophilic substitution using NaN_3_ [acetone/H_2_O (3:1), 20 min, 25 °C] to obtain the azido methacrylate derivative **5** in nearly quantitative yield. To obtain nitrile oxide derivative **7**, first chlorooxime **7** (90%) [NCS (1.1 eq), DMF, 40 °C, 3 h] was prepared from commercially available benzaldehyde oxime **6**. The treatment of **7** with triethylamine (TEA; 2 eq) leads to the *in-situ* generation of the nitrile oxide derivative **7**. Its reaction with **4** forms the 5-disubstituted isooxazoline compound **9** in good yield (up to 80%), as a racemic mixture.^29^ Regarding the regioselectivity, the oxygen atom of the nitrile oxide preferentially reacts with the most substituted carbon atom of the dipolarophile, generating only the product substituted at the 5-position. The regioisomer at the 4-position was not observed, nor in traces. The methyl ester group of **9** was hydrolyzed to produce the corresponding carboxylic acid **9** (quantitative yield) through a standard saponification reaction [LiOH (2 eq), THF/H_2_O, 25 °C, 30 min]. To resolve the racemic mixture, accordingly with our designed sequences, we performed a first coupling with H-Val-OMe hydrochloride salt (1.5 eq) in the presence of propylphosphonic anhydride (T3P; 3 eq) and DIPEA (4 eq) at pH 8-9. This reaction allowed the formation of a mixture of **10a** and **10b** with an overall yield of 97%. Subsequently, the two diastereoisomers **10a** and **10b** were separated by flash chromatography (DMC/Et_2_O, 98/2) in 42% and 47% yield, respectively. As reported above, the methyl ester of **10a** and **10b** was hydrolyzed, affording carboxylic acid derivatives **11a** and **11b** in quantitative yields without any trace of Val-racemization, as proved by NMR analysis. Crystals were prepared by slow evaporation of a solution containing **11b** dissolved in Et_2_O/*n*-Hexane to define the correct absolute stereochemistry. The X-Ray analysis of the obtained **11b** crystals allowed the determination of the absolute *R*-stereochemistry of C-5 of the isooxazoline ring (Figures 1-4S and Table 1S, SI). Indirectly, the *S*-stereochemistry was assigned to diastereoisomer **11a**.

The azido moiety of **11** was then reduced through a Staudinger reaction to afford the amine intermediate of the Isox-β^2,2^-AA [PMe_3_ (1.1 eq), H_2_O (7 eq), THF (0.1 M), 16 h, rt]. This crude product was then removed from the solvent and directly used for the next synthetic step. By treating compounds **11** with Fmoc-succinimide (Fmoc-OSu) in DCM and DIPEA (2 eq) (4 h, from 0°C to rt), protected compounds **12a** and **12b** were obtained in good yields (65% and 70%, respectively over three synthetic steps), as compatible intermediates for a SPPS synthetic approach.

**Scheme 1.**
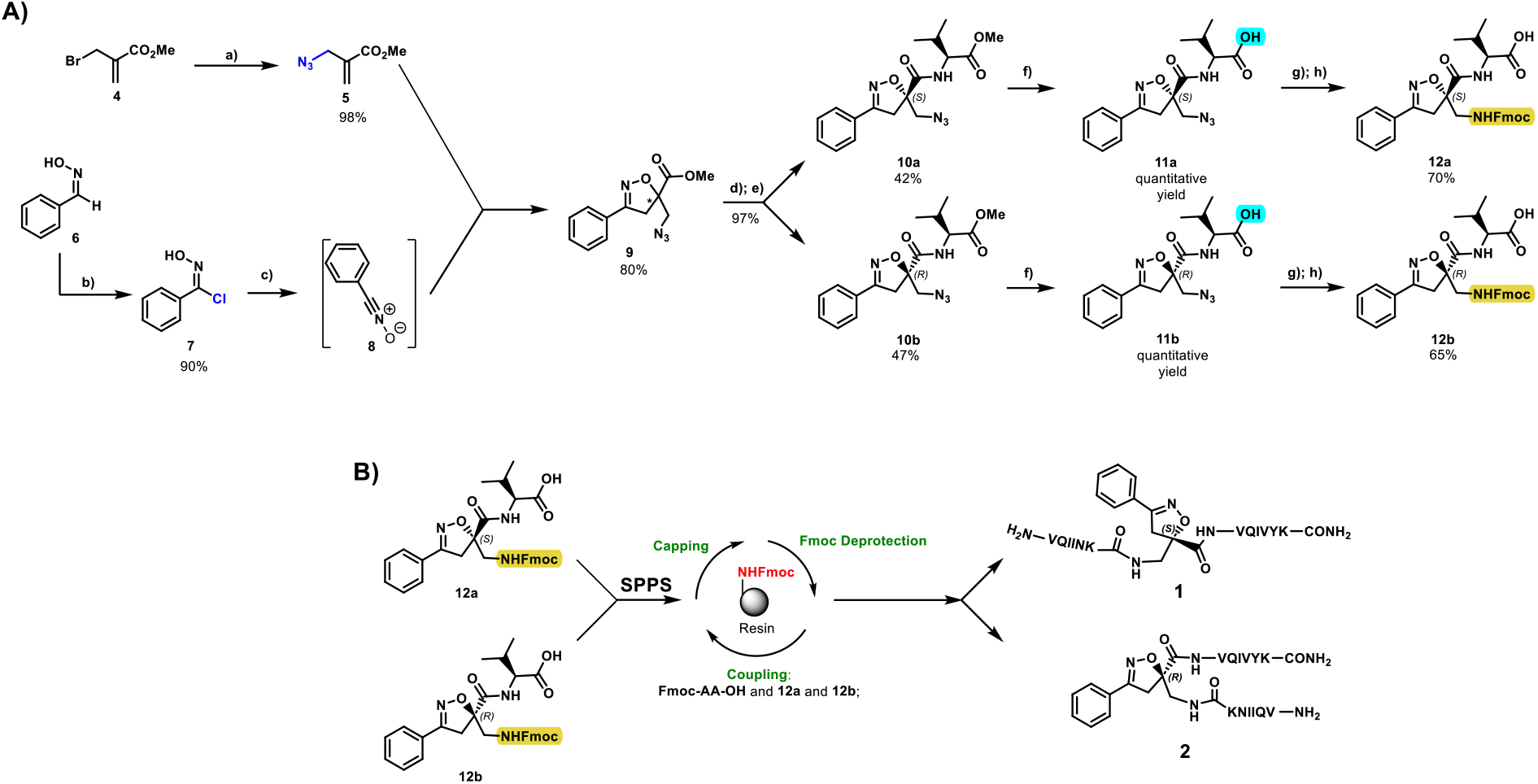
A) Liquid phase synthesis: **a**) NaN_3_ (2 eq), acetone/H_2_O (3:1), 20 min, r.t.; **b**) NCS (1.1 eq), DMF, 40 °C, 3h; **c**) **5** (1 eq), **7** (1.1 eq), TEA (2 eq), r.t., 16h; **d**) LiOH 0.1M in H_2_O (2 eq), THF (1M), r.t., 30 min; **e**) HCl·H-Val-OMe (1.5 eq), T3P (3 eq), DIPEA (3 eq), DCM, r.t., 16h; **f**) LiOH 0.1M in H_2_O (1.2 eq), THF (1M), r.t., 30 min; **g**) PMe_3_ (1.1 eq), H_2_O (7 eq), THF (0.1 M); **h**) Fmoc-OSu (1.1 eq), DIPEA (2 eq), DCM (0.1 M), 0°C to r.t., 4h; **B) Lyberty Blue synthesis: i**) 20% piperidine in DMF (65s, 90°C, mw); **j**) Fmoc-AA-OH (5 eq), DIC (5 eq), OXYMA (5 eq) (110s, 90°C, 40W, mw); **c**) 3 mL 20% piperidine in DMF (2×20’); **k**) Fmoc-Isox-β^2,2^-AA-OH (2 eq), DIC (2 eq), OXYMA (2 eq), DMF/DCM, r.t., 16h; **l**) acidic cleavage (trifluoracetic acid/water/thioanisol/triisopropyl silane; 95/2.5/1.25/1.25) for 2 hours; **m**) H_2_O/ACN + 0.1% TFA; 20 to 70 gradient ACN in 20 minutes.

The synthesis of the final compounds **1** and **2** was performed using the Liberty BlueTM (CEM) Automated Microwave Peptide Synthesizer under classical conditions (Scheme 1, B): the coupling reaction was accomplished at 25 °C for 120 s, followed by 480 s at 50 °C and 35 W. We used Rink-amide resin (0.1 mmol scale, resin loading = 0.55 mmol/g) and commercially available Fmoc-AA-OH (5 eq, 0.2M), DIC (5 eq, 0.2M), OXYMA (5 eq, 0.2M), previously solubilized in DMF. The Fmoc group was cleaved with a standard deprotection protocol (20% piperidine in DMF) at 75 °C, 155 W for 15 s followed by 60 s at 90 °C, 50 W. The coupling of **12a** and **12b** was manually performed using the pre-activation step in the presence of 2 equivalents of **12a** or **12b**, DIC (2 eq), and OXYMA (2 eq) in a DCM/DMF (1/3) mixture during 16 hours at room temperature. Afterwards, the remaining amino acids were coupled under the analog conditions indicated above. Finally, after resin cleavage using an acidic cocktail (TFA/H_2_O/thioanisol/triisopropyl silane; 95/2.5/1.25/1.25). The final peptidomimetic **1** and **2** were purified by semipreparative HPLC using H_2_O/ACN + 0.1% TFA with a 20 to 70 gradient in 20 minutes.

Compound **3** was synthesized using the same synthetic approach, with the exception of the diastereomeric mixture being separated by flash chromatography (Hex/EtOAc, 3/2) after two consecutive couplings (Thr and Lys) on intermediate **9** instead of one (for details, see SI, Schemes S1 and S2).

### Conformational study and self-aggregation assessment with and without heparin of compounds (1) and (2)

Compounds **1** and **2**, both bearing the two nucleation sites of tau, were predicted to be self-aggregative. So, for the assessment of their respective conformation and self-aggregation behavior, according to the to the *(S)* or *(R)* stereochemistry of the scaffold, they were preventively subjected to the HFIP treatment before being employed, to ensure as much as possible their monomerization.

Circular dichroism (CD) analyses were conducted at 125 µM concentration at room temperature in 10 mM phosphate buffer (PB) at pH 7.2 (Figure 2A). Both compounds showed a negative broad band at around 220 nm and a positive band at 198 nm, characteristic of β-sheet structures. At the same concentration, the CD spectrum of **1** is less intense than compound **2**, characteristic of self-assembly peptides.^35^

**Figure 2.**
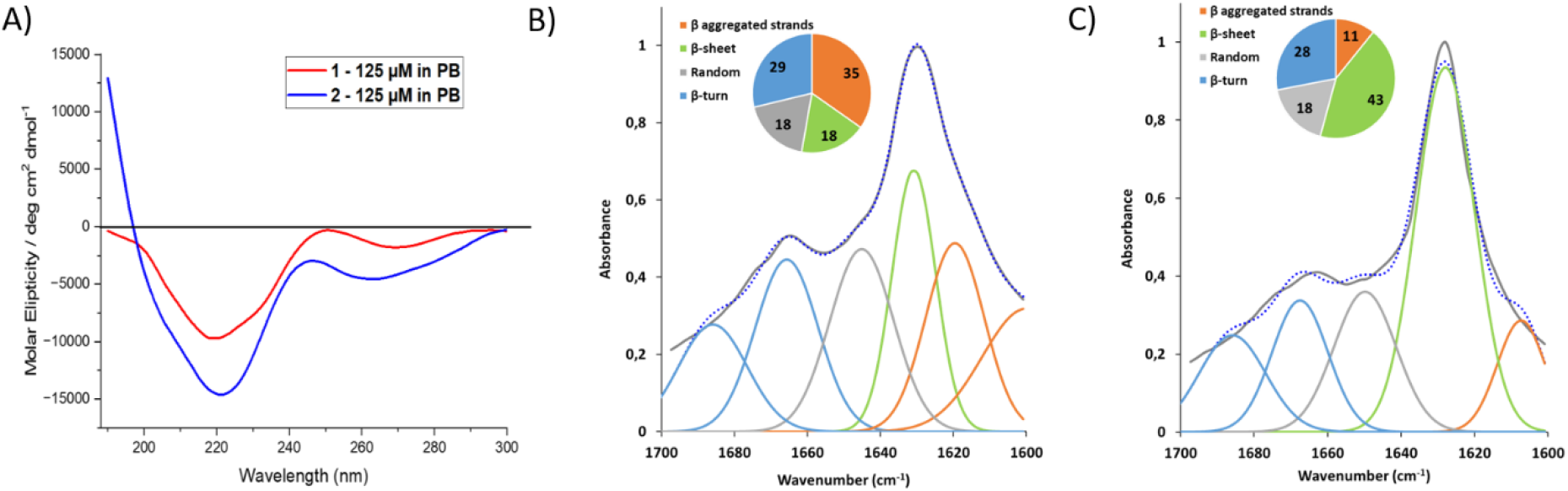
**A**) CD spectra of compounds **1** and **2**; **B-C**) IR-ATR amide I deconvolution of **1 (B)** (error squared = 0.029, standard deviation 8.83) and **2 (C)** (error squared = 0.042, standard deviation 7.16) with the schematic representation of the secondary structure motif. Both compounds were pre-treated with HFIP and immediately employed at the concentration of 125 µM in 10 mM PB, pH 7.2.

Both compounds were also subjected to IR analysis in 10 mM PB (pH 7.2) at the same concentration of 125 µM. As shown in Figure 2 B-C, the amide I deconvolution of **1** showed a predominant content of aggregated strands (35%), β-sheet structures (18%) and β-turn (29%) (Figure 2B). On the other hand, compound **2** displayed a minor content of aggregated strands (11%), a higher content of β-sheet (43%), and similar contents of β-turn (28%) (Figure 2C). Taken together, the results agree with the CD spectra of **1** and **2**. Compound **1** with the *S*-Isox-β^2,2^-AA showed a greater tendency to self-assembly in β-sheet rich insoluble structures than **2**, containing *R*-Isox-β^2,2^-AA, which seems to be more stable as a β-hairpin conformation and less prone to stack into supramolecular structures.

The hexapeptide PHF6* is often used as a model for understanding the aggregation of the full-length tau protein.^9,36,37^ It is well known that PHF6* at 25 µm in the presence of heparin shows an aggregation kinetic profile very similar to PHF6, reaching a plateau within 30 min. So, we first investigated the self-aggregation propensity and β-sheet formation of compounds **1** and **2** by employing the same protocol as the one described in the literature for Ac-PHF6*-NH_2_,^37^ using a Thioflavin-T (ThT) fluorescence spectroscopy assay (Figure 3A). The assay was performed in the presence of heparin (1 µM) in 20 mM MOPS buffer at pH 7.4 (Figure 3A). After 60 min, compound **1** showed an extremely high ThT fluorescent signal, nearly nine times higher than the model Ac-PHF6*-NH_2_, confirming a strong self-aggregative behavior. On the other hand, the ThT signal in the presence of compound **2** did not increase, suggesting no self-aggregation (Fig. 3A). All the results are in agreement with the previous observations by CD and IR. However, to further investigate if this self-aggregation persists even without heparin (as observed in CD and IR), both compounds were assessed by ThT fluorescence spectroscopy in the same conditions of the previous physical-chemical analyses (PB 10 mM pH 7.2). Compound **1** demonstrated to maintain its aggregation propensity in phosphate buffer, while compound **2** didn’t show any aggregation (Figure 3B). Interestingly, lowering the pH from 7.2 to 5.1 led to a considerable decrease in the ThT signal of compound **1** (Figure 3B), demonstrating less aggregation propensity at pH 5.1, also confirmed by the CD analysis (Figure S7, SI). Finally, both compounds were also tested at 1 µM concentration under the conditions for tau aggregation assessment, and none of them showed self-aggregation in the absence of heparin. All together, these results allowed to highlight a different self-aggregation behavior of compounds **1** and **2** according to the *(S)* or *(R)* stereochemistry of the scaffold, even if they are composed by the same hexapeptide sequences, and to find the correct conditions to ensure a stable monomerization.

**Figure 3.**
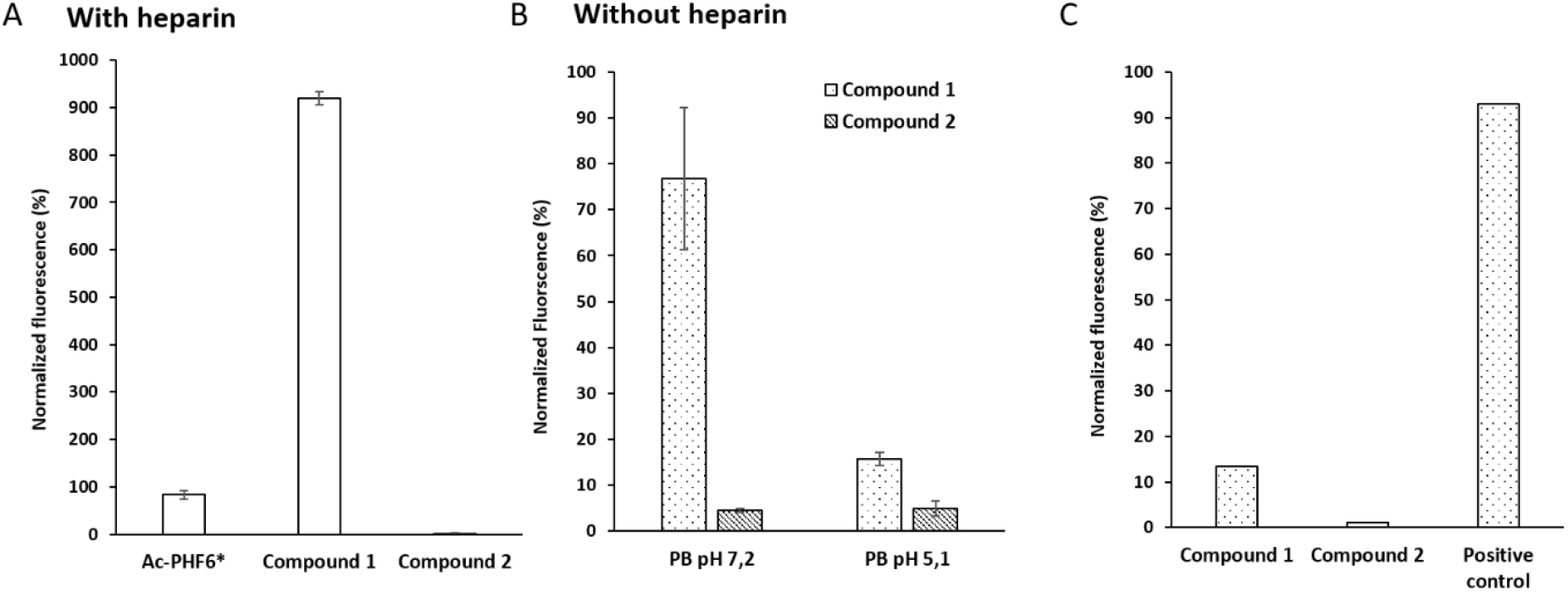
ThT fluorescence spectroscopy assay: A) Self-assembly of compounds **1** and **2** (25 µM) assessed in comparison to the model peptide Ac-PHF6* (25 µM) in MOPS 20 mM buffer at pH 7.4 in the presence of 1 µM of heparin, as reported in the literature.^37^ Fluorescence data have been normalized by putting the max of fluorescence of Ac-PHF6* as 100%. (B) Self-assembly of compounds **1** and **2** (25 µM) in the absence of heparin in phosphate buffer at two different pH, 7.2 and 5.1. All results are represented as the average of three replicates (n = 3) and the error bars are indicated as the ±SEM. C) Self-assembly without heparin of compounds **1** and **2** (1 µM) without heparin under the conditions for tau aggregation assessment (25 mM PB buffer at pH 6.6). Positive control: tau (10 µM) in the presence of heparin (0.1 µM).

To confirm that **1** and **2** were able to adopt a stable extended conformation and a β-hairpin-like structure, respectively, 1D and 2D homo and heteronuclear NMR conformational studies were performed in 10 mM PB, at pH 5.1 (final peptide concentration of 3.6 mM). This pH value was chosen because in this condition both compounds have a comparable high monomer stability (Figure 3B and Figure S7, SI).

^1^H and ^13^C resonances were completely assigned using 1D 1H WATERGATE, 2D ^1^H-^1^H COSY (20 ms mixing time), 2D ^1^H-^1^H DIPSI (20 ms mixing time), 2D ^1^H-^1^H ROESY (20 ms mixing time) and 2D ^1^H-^13^C HSQC, 2D ^1^H-^13^C HMBC. The resonances were assigned using spectrum acquired at 283K, other supplementary spectra were recorded respectively at 298K, 318K, 328K for **1** and 293K, 303K and 313K for **2**, to evaluate the presence of intramolecular hydrogen bonds through temperature coefficient calculation. ^1^H and ^13^C chemical shifts were calibrated on the methyl moiety of aliphatic amino acids. The vicinal coupling constants (^*3*^*J*) were extracted from 1D ^1^H and 2D ^1^H-^1^H COSY spectrum at 283K.

^1^H NMR spectrum of **1** showed a good dispersion of the NH chemical shifts indicating a single conformation. A complete attribution of all the corresponding chemical shifts is provided in Table S2 (see SI).

The vicinal ^*3*^*J*_NH-Hα_ coupling constant showed an average value of 7.4 ± 0.3 Hz, discarding the possibility of a folded structure. However, this value appears to be closer to the coupling constant found in β-sheets (8.9 Hz for antiparallel and 9.7 for parallel) than in α-helices (3.9-4.2 Hz) (Figure 4 A). Both peptide arm sequences present a complete set of strong CH/NH (*i, i+1*) ROEs, suggesting an extended conformation for each peptide arm (Figure 4 B, blue and red curves). Additionally, this hypothesis is supported by the lack of intra-strand NH/NH (*i, i+1*) ROEs and the absence of inter-strand ROEs. Neither cross-strand ROEs interaction between the two peptide arms nor any spatial proximity between the -CH_2_ protons of the Isox-β^2,2^-AA^7^ and the NH/CH_α_ protons of the Val^8^ were observed.

**Figure 4.**
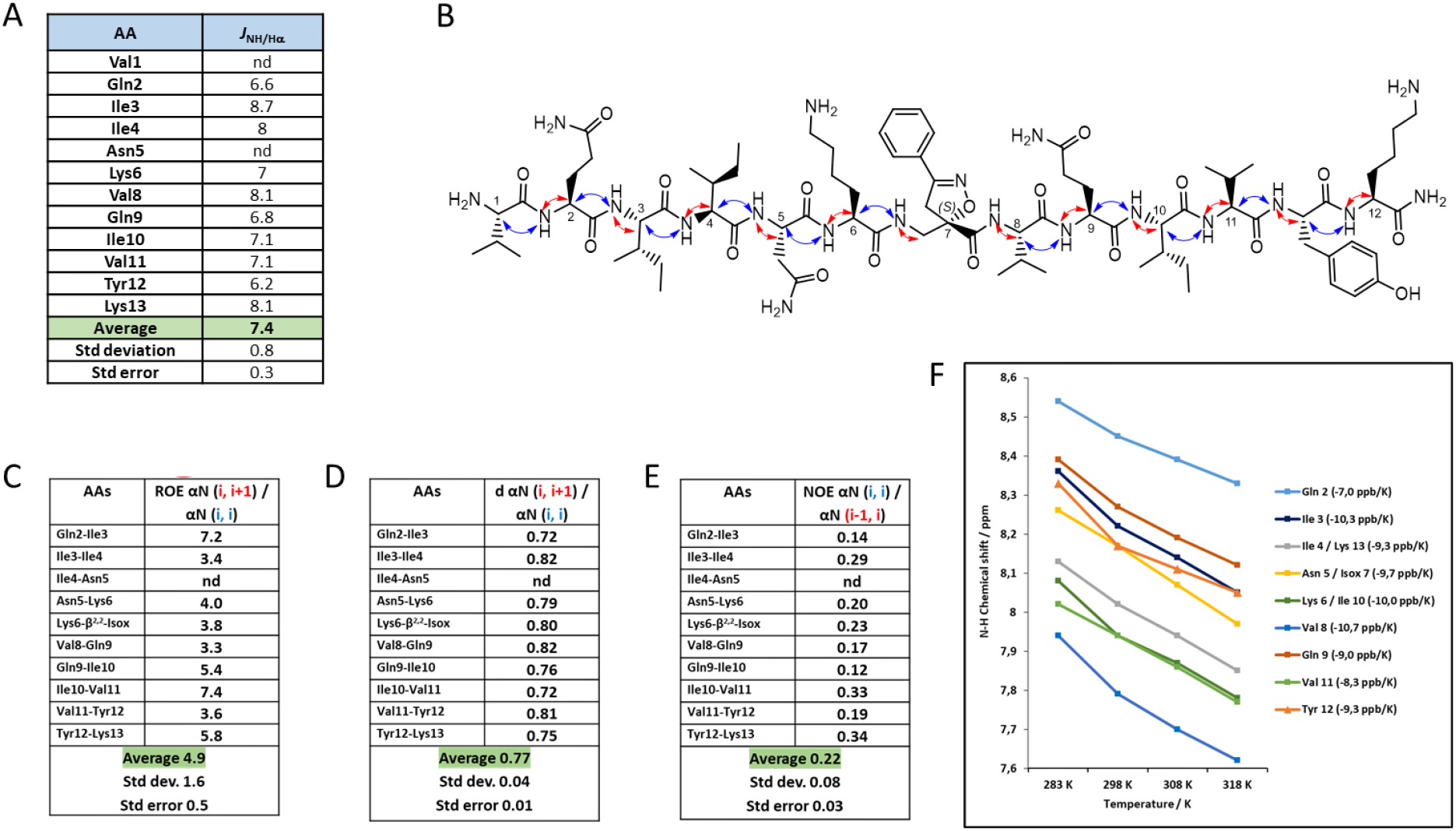
NMR conformational analysis of compound **1** in 20 mM phosphate buffer at pH 5.1; A. Values of ^3^J_NH-Hα_ coupling constants; B. Representation of the CH/NH (i, i) ROEs (red) and the CH/NH (i, i+1) ROEs (blue); C. ROE intensity ratio of αN (i, i+1) / αN (i, i); D. Hα-NH distances; E. ROE intensity ratio of αN (i, i) / αN (i-1, i); F. Temperature coefficients of amide protons between 283 and 318 K.

As demonstrated by Dobson *et al*.,^38^ the intensity ROEs αN (*i, i+1*) / ROEs NN (*i, i+1*) ratio can indicate the propensity of an amino acid to adopt a specific secondary structure. A ratio value close to 1 suggests increased α-propensity, values around 1.4 indicate the tendency to form random coil regions and values of approximately 2.7 reveal a higher-than-average β-propensity. For compound **1**, the lack of ROEs NN (*i, i+1*) leads to an elevated ratio, which is a characteristic of residues with higher-than-average β-propensity.

Additionally, ROEs αN (*i, i+1*) / ROEs αN (*i, i*) intensity ratio about 2.3 is predicted for the population-weighted random coil model, whereas values exceeding 4 are expected for β-strands.^39^ Upon calculation of this ratio for **1**, the resulting average value of 4.9 ± 0.5 suggests the presence of a general extended conformation (Figure 4 C).

Finally, ROEs αN (*i, i*) / ROEs αN (*i-1, i*) intensity ratio is known to be influenced by the ψ angle of residue *i* − *1*, with values ranging from around 6 for α-helices to roughly 0.25 for β-sheets (ratio < 1).^40^ A ratio less than 1 is indicative of β-sheets, whereas a ratio greater than 1 suggests α-helices.^41^ An average value of 0.22 ± 0.03 is in line with an extended conformation for compound **1** (Figure 4 E).

Furthermore, the ratio of sequential to intra-residue H_α_-NH distance varies according to the secondary structure: 1.25 for α-helices and 0.73 for β-strands (where d αN (*i, i+1*) α-helix = 3.5 Å and d αN (*i, i*) α-helix = 2.8 Å; d αN (*i, i+1*) β-sheet = 2.2 Å and d αN (*i, i*) β-sheet = 3.0 Å).^42^ In our case, with an average value of 0.77 ± 0.01, the data fits once more with an extended conformation adopted by **1** (Figure 4 D).

Any of the amide protons were involved in H-bond as demonstrated by all temperature coefficients (Δδ_NH_/ΔT) below -4.5 ppb·K^-1^, and an average value of -9.13 ppb·K^-1^ (Figure 4 F).

Even for compound **2**, the ^1^H NMR spectrum showed a good dispersion of the NH chemical shifts indicating the presence of a single conformation. A complete attribution of all the corresponding chemical shifts is provided in Table S3 (see SI). The average vicinal ^*3*^*J*_NH-Hα_ coupling resulted to be 7.7 ± 0.3 Hz, suggesting an extended conformation of both peptide arms. Remarkably, although the *J* coupling constants of **2** are similar or slightly larger than **1**, an interesting difference is observed in the ^*3*^*J*_NH-Hα_ coupling constant of Lys^6^ in **2**. Lys^6^ is located just before the Isox-β^2,2^-AA, and exhibits a reduction from 7 Hz in **1** to 6.4 Hz in **2**. According to the Karplus formula, this reduction is consistent with a smaller φ dihedral angle, and, therefore, provides evidence for the formation of a local turn (Fig. 5 A).

**Figure 5.**
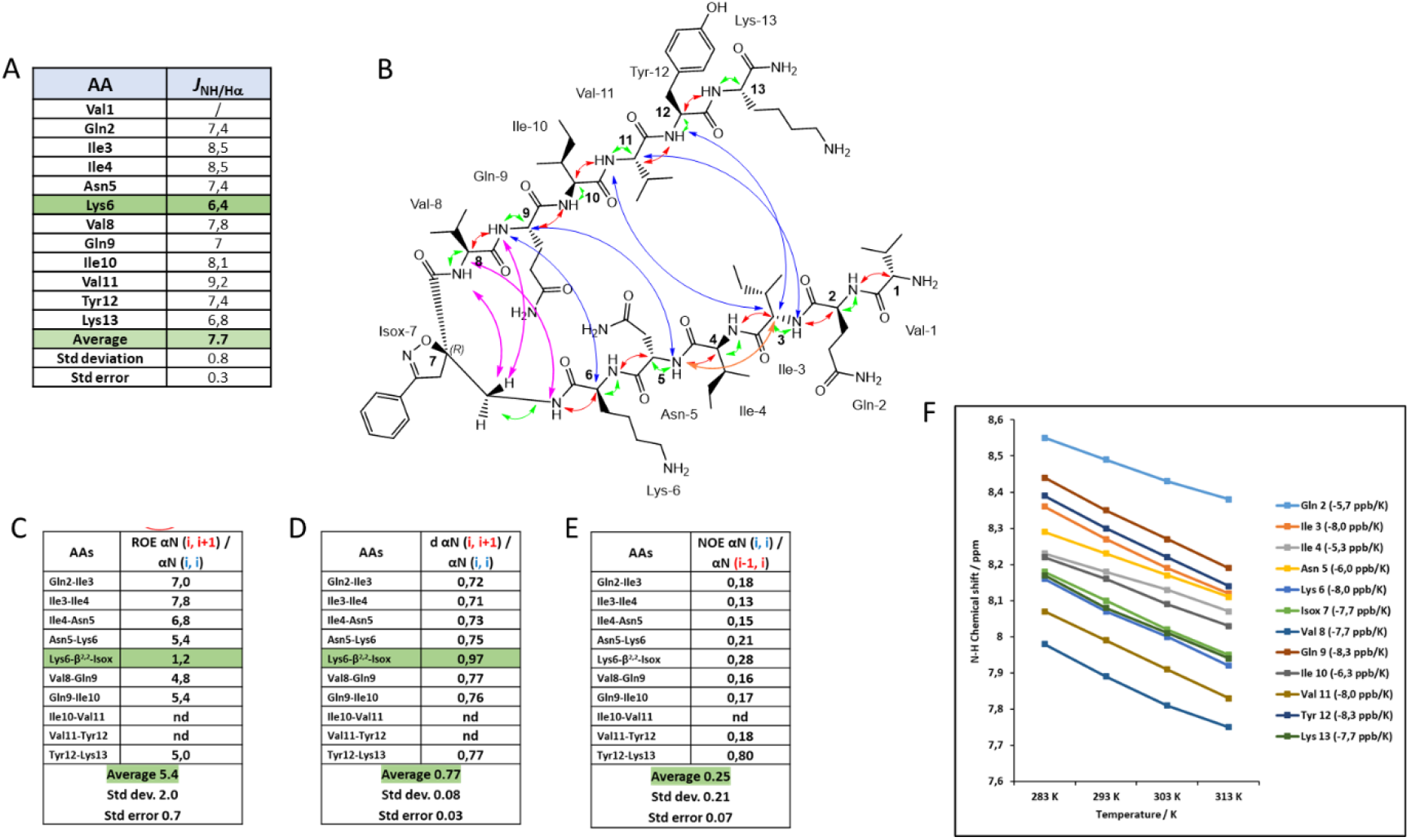
NMR conformational analysis of compound **2** in 20 mM phosphate buffer at pH 5.1; A. Values of ^3^J_NH-Hα_ coupling constants; B. Representation of the CH/NH (i, i) ROEs (green), the CH/NH (i, i+1) ROEs (red), the intra-strand NH/NH (i, i+1) ROEs (orange), the inter-strand CH-NH ROEs in the turn region (purple), and the long-range inter-strands ROEs (blue); C. ROE intensity ratio of αN (i, i+1) / αN (i, i); D. Hα-NH distances; E. ROE intensity ratio of αN (i, i) / αN (i-1, i); F. Temperature coefficients of amide protons between 283 and 318 K.

The ROESY analysis showed a complete set of strong sequential CH/NH (*i, i+1*) ROEs (Figure 5 B, red curves) for both peptide arms, suggesting an extended conformation, also demonstrated by the absence of intra-strand NH/NH (*i, i+1*) ROEs, leading to an elevated ROEs αN (*i, i+1*) / ROEs NN (*i, i+1*) ratio, which is a characteristic of residues with higher-than-average β-propensity. The resulting average value of 5.4 ± 0.7 for the ROEs αN (*i, i+1*) / ROEs αN (*i, i*) intensity ratio also suggests the presence of extended conformations on both peptide arms. Worthy of particular attention, the only ROEs αN (*i, i+1*) / ROEs αN (*i, i*) ratio lower than 2.3 is the one between Lys^6^ (*i*) and Isox-β^2,2^-AA^7^ (*i+1*), suggesting a different orientation of the H_α_ (Lys^6^) with respect to the NH of the Isox-β^2,2^-AA^7^, due to a reduced ϕ dihedral angle value, as seen before for the ^*3*^*J*_NH-Hα_ coupling constant (Figure 5 C).

Furthermore, an average value of 0.25 ± 0.07 for ROEs αN (*i, i*) / ROEs αN (*i-1, i*) intensity ratio is still in line with an extended conformation of the peptide sequences of compound **2** (Figure 5 E).

Finally, an average value of 0.77 ± 0.03 for the ratio of sequential to intra-residue H_α_-NH distances well fits with a β-strand conformation adopted by the two peptide arms of compound **2**. Once again, a shift towards 1 (0.97) for Lys^6^ (*i*) and Isox-β^2,2^-AA^7^ (*i+1*) is observed, compatibly with the presence of a turn in this part of the molecule (Figure 5 D).

Interestingly, in contrast to **1**, multiple cross-strands ROEs were observed for compound **2**. The turn conformation is confirmed by the presence of inter-strand CH-NH ROE signals between H_α_ of Val^8^ and NH of Isox-β^2,2^-AA^7^, H_β_ of Isox-β^2,2^-AA^7^ and NH of Gln^9^, and H_β_ of Isox-β^2,2^-AA^7^ and NH of Val^8^ (purple arrows Figure 5 B). Additionally, further long-range inter-strand interactions between H_α_ of Lys^6^ and NH of Gln^9^, H_α_ of Gln^9^ and NH of Asn^5^, H_α_ of Val^11^ and NH of Ile^3^, H_α_ Ile^3^ and NH of Val^11^, and H_α_ of Ile^3^ and NH of Tyr^12^ confirm the hairpin conformation alongside the full peptide arms (Figure 5 B blue arrows; and Figure S8 in SI). A single long-range intra-strand is found and highlighted with an orange arrow between the NH of Asn^5^ and the H_α_ of Ile^3^, probably indicating a mild twist in this portion (Figure 5 B).

As reported in Figure 5 F, any of the amide protons are involved in stable H-bond, as demonstrated by all temperature coefficients (Δδ_NH_/ΔT) below -4.5 ppb·K^-1^, and an average value of -7.2 ppb·K^-1^. It has to be noticed that the two borderline values for Ile^4^ and Val^8^ (5.3 and 5.7 ppb·K^-1^, respectively) suggest a partial involvement of the two amide protons in weak H-bonds, and this will be in accordance with an alternating 10- and 14-membered rings, typical of antiparallel β-sheet.

### Tau Aggregation in the presence of compound (1) and (2): Evaluation by Thioflavin-T fluorescence spectroscopy, Transmission Electron Microscopy and Fluorescence Decay After Photoactivation

#### Tau aggregation in the absence of heparin is triggered by (1) and not by (2)

To investigate whether the two distinct conformations of compounds **1** and **2** have the potential to induce or prevent Tau aggregation, we exploited ThT fluorescence spectroscopy to obtain real-time information about the supramolecular self-assembly of tau in the presence of **1** and **2**.

We opted to test the compounds at a sub-stoichiometric ratio (1 µM) for two primary reasons. Firstly, we observed that, at higher concentrations, compound **1** exhibits a notable increment in its self-aggregative tendency, thus making the interpretation of the results more challenging (see Figure 3B). Moreover, at this concentration no aggregation of both compounds was observed (Figure 3C, Figure S7 and S9 E and F, SI). Secondly, by working at the sub-stoichiometric ratio, it can be demonstrated that the presence of a minimal amount of a seed-competent mimic, as compound **1**, could drive the full-length wt-tau towards the formation of fibrils.^5,43^

Therefore, the two compounds have been incubated with wt-Tau441 (10 µM) at 0.1/1 ratio without heparin in PB buffer (pH 6.6) at 37°C, and the ThT (25 µM) fluorescence emission has been monitored over time. To have a positive control, the aggregation of wt-Tau441 was induced by the addition of a low quantity of heparin (100/1 ratio, 0.1 µM) to provide evidence of the mandatory use of anionic co-factors like heparin to induce the aggregation of Tau *in vitro* and compared to wt-Tau441 without heparin (negative control).

As previously described, the ThT fluorescence value of wt-Tau441 without heparin (orange line) remains at its basal level during 140 hrs., showing that wt-Tau441 under this condition cannot form amyloid fibrils (Figure 6 A), which was confirmed by TEM (Figure S9, SI). On the other hand, in the case of wt-Tau441 in the presence of heparin (0.1 µM), the ThT fluorescence (green line) shows the classical sigmoidal curve characterized by a lag phase (up to 5h), the elongation phase and the plateau reached after approximately 60 hrs. (Figure 6 A). This curve follows a molecular model (MRE = 0.00249) in which the mechanism dominating the kinetics of aggregation is the nucleation/elongation (AmyloFit),^45^ characterized by rate constant of elongation and primary nucleation (*k+k*_*n*_) of 4.07×10^29^ conc^-n^ time^-2^. The occurrence of amyloid fibrils with a diameter around 20 nm (Figure 6 B, left) with planar and a helical organization of a single protofilament was detected by TEM (Figure 6B, see arrows).

**Figure 6.**
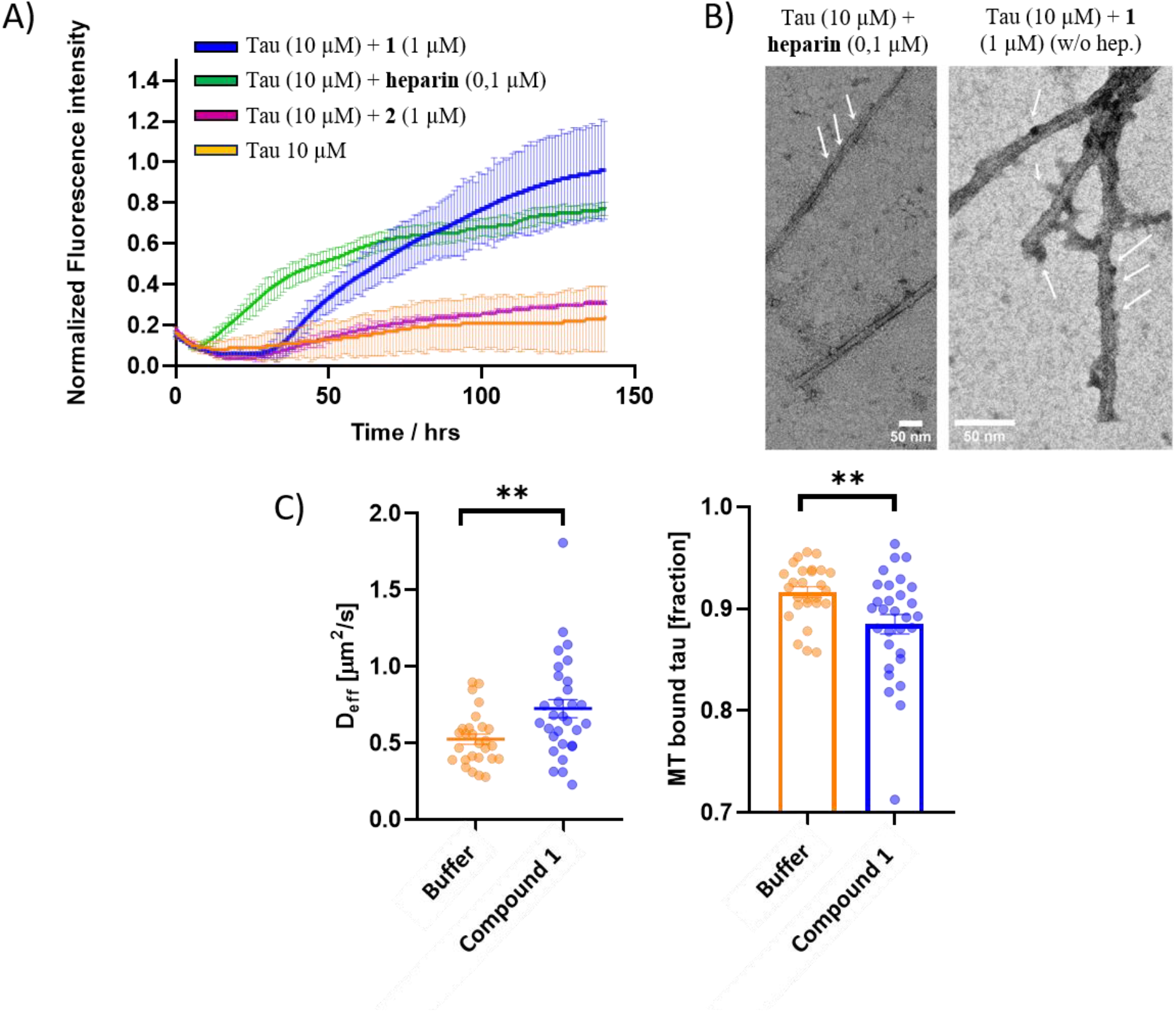
Assessing tau fibrillation in the presence of compounds **1** and **2** A) ThT fluorescence spectroscopy of tau (10 µM) in PB buffer (pH 6.6) at 37°C in the presence of 0.1 µM of heparin (green line), 1 µM of compound **1** (blue line), and 1 µM of compound **2** (purple line), or in the absence of any additive (orange line). Fluorescence data were normalized, and the results are represented as the average of three replicates (n = 3); B) Representative TEM images of tau fibrils formed in the presence of heparin (0.1 µM) on the left, and with compound **1** (1 µM) on the right; arrows are showing the helical organization of tau alone (left) and the presence of oligomers on the surface of the fibrils (right). To see the low magnification images, please refer to Figures S9 B and S9D; C) Fluorescence decay after photoactivation (FDAP) analysis of **1** in differentiated model neurons (PC-12 cells) transfected with an aggregation-prone tau construct (PAGFP-Tau441 ΔK280). The decay is fitted with a mathematical model, which allows the analysis of the effective diffusion constant (D_eff_) and the MT-bound fractions of tau in axon-like processes as an indicator of the effect of the compound on tau aggregation.^44^ Presence of **1** increases Tau D_eff_ while decreasing Tau MT bound fraction, indicating that 1 induces incresed tau aggregation in the neuronal cells. Experiments were performed with PC12 cells differentiated with NGF, 24 hrs. pre-incubation time, DMSO concentration 0.125% (Control).

Interestingly, wt-Tau441 in the presence of compound **1** (1 µM) and without heparin (Figure 6 A, blue line) shows a ThT fluorescence increase with a classical sigmoidal curve with a lag phase of about 30 hours and a growth phase until 140 hours, slightly different from the one of control wt-Tau441:heparin. The AmyloFit fitting of the curve suggests a molecular model (MRE = 0.00291) in which the mechanism dominating the kinetic of aggregation is the secondary nucleation (*k*_*+*_*k*_*n*_ = 9.00×10^5^ conc^-n^ time^-2^ and *k*_*+*_*k*_*2*_ = 1.82×10^16^ conc^-n^ time^-2^). This different kinetic suggests a different aggregation mechanism. Figure 6 B (right), shows a change in the morphology of the wt-Tau441 in the presence of compound **1**. These fibrils are thinner than tau fibrils in the presence of heparin, with a diameter of around 12 nm and surrounded by oligomers on their surface, confirming the secondary nucleation mechanism that the ThT assay suggested. On the contrary, wt-Tau441 (10 µM) in the presence of compound **2** (1 µM) (Figure 6 A, purple line and TEM images in Figure S9, SI) doesn’t exhibit any aggregation similar to the negative control (tau in the absence of heparin). This result demonstrates that the β-hairpin conformation of **2**, mimicking the inert conformation of tau, cannot trigger the aggregation process. In contrast, compound **1**, through its extended conformation, behaves as a seed-competent template, allowing tau to fold in a seed-competent monomer by similarity of conformation.

Fluorescence Decay After Photoactivation (FDAP) is a live-cell microscopy technique applied to study tau-microtubule interaction and investigate tau mobility in neuronally differentiated cells.^46^ Parameters such as the effective diffusion constant (D_eff_), the tau microtubule-bound fraction (MT bound tau), the association (K_on_), and dissociation (K_off_) kinetic constants ^47,48^ can be derived by this analysis to evaluate potential changes in the tau-MT interaction in axon-like processes of model neurons. The constructs are N-terminally tagged with photoactivatable GFP (PAGFP) and exogenously expressed in PC-12 cells differentiated into a neuronal phenotype and locally photoactivated using a laser scanning microscope. When used with an aggregation-prone tau construct (Tau441 ΔK280 construct), the assay can be used to determine the effect of exogenously added compounds on modulating tau aggregation.^44^ If a compound is supposed to induce the aggregation of tau in a cell environment, its effective diffusion constant results to be increased due to a pronounce aggregation. Conversely, when the compound prevents the aggregation, the diffusion constant decreases and tau interacts more with MT.

The toxicity of compound **1** alone was tested through a MTT cell viability assay and in different concentrations up to 50 µM. It did not show any statistically significant modification of cellular metabolism on differentiated PC-12 cells (Figure S10, SI). FDAP analysis was performed at a concentration of 25 µM of the pro-aggregative compound **1**. Remarkably, compound **1** significantly increased the effective diffusion constant (D_eff_), indicating a reduced interaction of Tau with the MTs (Figure 6 C). Accordingly, the MT-Tau bound fraction calculation showed a significant decrement compared to the control, indicating that compound **1** induces a substantial increase in tau aggregation. We did not observe changes in K_on_ and K_off_ (Figure S11, SI), indicating that compound **1** does not affect the tau/MT kinetics.

Compound **1**, employed as a chemical model system of a seed-competent conformation, allowed us to shed light on the early mechanism behind Tau aggregation. It provided evidence that when the two hexapeptide motifs, PHF6* and PHF6, are in an extended conformation, the aggregation of Tau is accelerated, both in vitro and in cells.

### Tau aggregation in the presence of heparin can be controlled by the concentration of the β-hairpin compound (3)

As aforementioned, compound **3** was designed according to previous data showing that a β-hairpin mimic, bearing defined sequences (S4 and S7) from Hsp90 (Figure 1 C), can inhibit both wt-tau and pro-aggregative ΔK280 tau mutant fibrillation.^33^ To verify the ability of the *(R)*-configured Isox-β^2,2^-AA to induce a stable β-turn like folding in a long peptide sequence and, thus, to be employable as β-turn inducer for potential inhibitors of amyloid aggregation, the S4 and S7 peptide sequences of N-terminal portion of Hsp90 were attached to the *(R)*-configured Isox-β^2,2^-AA. Compound **3** forms a stable β-hairpin conformation as demonstrated by CD and IR analyses (Figure S13, SI). The activity of **3** as an inhibitor was assessed according to a classical protocol employing heparin to induce wt-tau aggregation.^49^ The inhibitor activity of **3** resulted dependent on its concentration (Figure 7 A-B). In the absence of **3**, the ThT-fluorescence curve of wt-tau/heparin (10 µM / 0.1 µM) showed a classical sigmoidal shape with a lag phase of 3 hours, followed by an elongation phase and a final plateau reached after 14 hours (Figure 7 A). In the presence of compound **3** at 5/1 and 1/1 ratios, the ThT-fluorescence curve of wt-tau/heparin displayed a complete inhibition of the aggregation, confirmed by the absence of aggregates in TEM (Figure S14, SI). The addition of a sub-stoichiometric quantity of **3** (0.1/1 ratio) shows a ThT kinetic curve similar to the one of tau in the presence of heparin alone, fitting well with a molecular model (MRE = 0.00430) in which the mechanism dominating the kinetic of aggregation is the secondary nucleation, with rate constants (*k*_*+*_*k*_*n*_ = 2.80×10^8^ conc^-n^ time^-2^ and *k*_*+*_*k*_*2*_ = 1.59×10^19^ conc^-n^ time^-2^) slightly higher than the tau-heparin control (see above). Considering the morphologies, fibrils in the presence of heparin alone are longer and more uniform than the straight filaments in AD-tau seeded filaments (Figure 7 B left).^50^ Surprisingly, the morphology of tau in the presence of 0.1 µM of **3** led to the formation of heparin-assembled recombinant tau fibrils that are exclusively short and straight, constituted by one to three filaments associated laterally, more similar to the paired helical filaments (PHF) and straight filaments (SF) in AD tau.^51^

**Figure 7.**
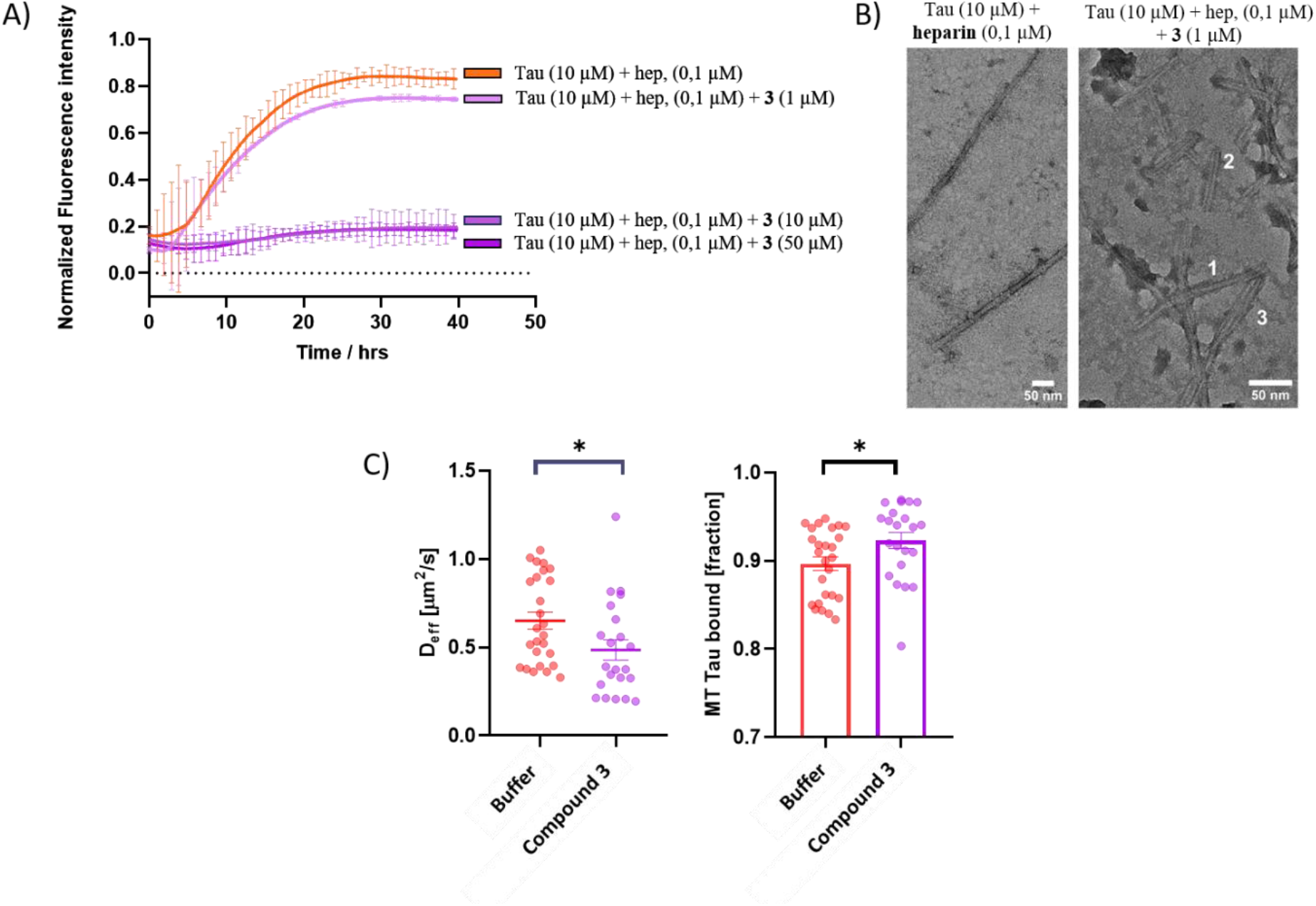
A) ThT fluorescence spectroscopy assessment of compound **3** in the presence of tau (10 µM) in NaPi buffer (pH 6.6) at 37°C and 0.1 µM of heparin at different molar ratios (1:5, 1:1 and 0.1:1, tau/**3**). Fluorescence data were normalized and the results are represented as the average of three replicates (n = 3); B) TEM images of tau fibrils formed in the presence of heparin (0.1 µM) on the left and with compound **3** (1µM) and heparin (0.1 µM) on the right; the number shows the amount of lateral association. 1 for only 1 filament, 2 for two filaments and 3 for three filaments; no fibrils were detected when compound 3 was present at 5:1 and 1:1 ratios; C) FDAP analysis of **3** on PC-12 cells transfected with PAGFP-Tau441 ΔK280. Mathematical modeling shows that compound **3** increases Tau D_eff_ and consequently decreases the tau MT bound fraction at a concentration of the compound of 25 µM, indicating that **3** inhibits tau aggregation in neuronal cells and restores the physiological tau-MT interaction in axon-like processes; Experiments were performed with differentiated model neurons (PC12 cells differentiated with NGF), compound incubation time was 24 hrs., DMSO concentration 0.125% (Control).

FDAP experiments to determine the effect of compound **3** in neuronal cells allowed us to demonstrate that β-hairpin **3** at 25 µM significantly decreases the effective diffusion constant of the aggregative prone Tau441 ΔK280 construct (D_eff_, Figure 7 C), indicating a raised interaction of tau with the microtubules (Figure 7 C) compared to the control. This indicates that compound **3** significantly decreases tau aggregation in neuronal cells thereby restoring a physiological tau-MT interaction. K_on_ and K_off_ parameters were not altered (Figure S15, SI), indicating that **3** reduces tau aggregation but does not affect the association and dissociation of tau to MT.

In conclusion, compound **3**, designed on the Tau-Hsp90 interaction and having a β-hairpin conformation, demonstrated remarkable inhibitory activity against tau aggregation, both *in vitro* and in cells. Furthermore, the data provide evidence that compound **3** is able to restore the physiological tau-MT interaction of aggregation-prone tau in model neurons.

## Conclusions

In this research article, we demonstrated that chemical systems, inspired by the conformations of non-pathological and pathological proteins, can be employed to modulate and study the amyloid misfolding and aggregation mechanism of tau. In particular, according to the stereochemistry of a new non-natural β^2,2^-amino acid, β-hairpin-like and extended conformations could be stabilized in long peptide sequences. Through this folding modulation, we showed that an extended conformation, exposing the hydrophobic residues of both sequences and driving away PHF6* from PHF6, can trigger the aggregation of tau in the absence of additives such as heparin, thus behaving like a seed-competent monomer model system. Conversely, a β-hairpin imitates the favorable folding of tau, allowing the protein to maintain its soluble monomeric form and potentially its MT interaction due to the absence of aggregation. Therefore, this β-hairpin conformation cannot induce the aggregation of tau *in vitro*. The extended conformation of our seed-competent monomer mimic can trigger the tau aggregation process, leading to the formation of fibrils, morphologically different from the ones obtained in the presence of heparin. These fibrils are the result of a different kinetic profile than the one observed with heparin. Indeed, a secondary nucleation-dominated mechanism drives wt-tau to form thinner fibrils, more similar to the ones found in the patients’ brains.^7^ Furthermore, a β-hairpin mimic, characterized by *R-*configuration of the non-natural β^2,2^-amino acid and based on the chaperone protein Hsp90, demonstrated the further application of this type of peptidomimetic foldamers for the development of inhibitors of tau aggregation. Surprisingly, at a sub-stoichiometric ratio and in the presence of low concentrated heparin, the same compound inspired by the chaperone protein Hsp90, can maintain a secondary nucleation-dominated kinetic, as observed with the seed-competent mimic, showing, however, morphology of the fibrils similar to the straight filaments observed in the brain of patients.^50^ Remarkably, our study shows that peptidomimetic foldamers are capable to modulate tau aggregation in living model neurons thereby influencing the tau-MT interaction in axon-like processes. In conclusion, chemical model systems are not only simpler and more accessible than full-length proteins, but can also be used to study the structures and folding modes of amyloid proteins, as well as to modulate the misfolding and aggregation of an amyloid protein in vitro and in cells to obtain a fibril morphology in vitro that approximates the pathologically relevant one.

## Acknowledgements

This research has received funding from the European Union’s Horizon 2020 research and innovation program H2020-MSCAITN-2019-EJD: Marie Skłodowska-Curie Innovative Training Networks (European Joint Doctorate)—Grant Agreement No: 860070—TubInTrain

Karine Leblanc is thanked for her contribution to the LC-MS analysis.

## Author contributions

The manuscript was written through contributions of all authors. D.D.L., M.L.G. and N.T. designed the experiments. D.D.L. performed the synthesis, the conformational analysis of NMR, IR and CD and, the ThT fluorescence assays; N.B. performed and analyzed MTT assays and quantitative live-cell imaging and produce tau protein; I.H. and V.D. performed TEM imaging; L.L.P. performed the X-ray analysis; R.BR. supervised N.B. for the MTT assays and quantitative live-cell imaging; R.BU., N.T. and M.L.G. supervised D.D.L. for the synthesis and the conformational analysis. S.O. revised the manuscript. NT and M.L.G. supervised the entire research.

All authors have given approval to the final version of the manuscript.

