## supplementary information for "Application of modular isoxazoline-β^2,2^-amino acid-based peptidomimetic foldamers as a chemical model system for studying the tau misfolding mechanism"

g New affiliation: CEA Saclay, DRF/JOLIOT/DMTS/SIMoS/LBM, 91191, Gif-sur-Yvette, France.

### General Procedures

Usual solvents were purchased from commercial sources. Thin-layer chromatography (TLC) analyses were performed on silica gel 60 F250 (0.26 mm thickness) plates. The plates were visualized with UV light ($\lambda$ = 254 nm) alternatively stained with a 4 % solution of phosphomolybdic acid or ninhydrin in EtOH.

NMR spectra of intermediates were recorded on an ultra-field Bruker AVANCE 300 ($^1$H, 300 MHz, $^{13}$C, 75 MHz) or on a Bruker AVANCE 400 ($^1$H, 400 MHz, $^{13}$C, 100 MHz); NMR spectra of **1**, **2** and **3** were registered with a Bruker spectrometer operating at 700 MHz equipped with cryoprobe. Chemical shifts $\delta$ are in ppm with the solvent resonance as the internal standard ($^1$H NMR, $CDCl_3$: $\delta$ = 7.26 ppm, $CD_3OD$: $\delta$ = 3.31 ppm, $CD_3CN$: $\delta$ = 1.93 ppm; $^{13}$C NMR, $CDCl_3$: $\delta$ = 77.16 ppm, $CD_3OD$: $\delta$ = 49.00 ppm; $CD_3CN$: $\delta$ = 1.3 ppm), and the following abbreviations are used: singlet (s), doublet (d), doublet of doublet (dd), triplet (t), quintuplet (qt), multiplet (m), broad multiplet (brm), and broad singlet (brs), broad doublet (brd). Mass spectra were obtained using a Bruker Esquire electrospray ionization apparatus. HRMS were obtained using a TOF LCT Premier apparatus (Waters) with an electrospray ionization source. The purity of compounds was determined by HPLC-MS on Agilent 1260 Infinity. Column: ATLANTIS T3 column (C18, 2.1 x 150mm-3µm), mobile phase: ACN/$H_2O$ + 0.1% TFA (gradient 1-30% in 15 or 20 min). Preparative HPLC was performed on Agilent 1260 Infinity II. Column: Pursuit (C18 10 x 250µm-5µm), mobile phase: ACN/$H_2O$ + 0.1% formic acid (FA) and on Waters XBridge BEH300 (C18, 2.1 x 150mm-5µM); POROSHELL 120 column (C18, 2.1 x 50mm-1.9µm); mobile phase: ACN/$H_2O$ + 0.1% formic acid (FA) or 0.1% Trifluoroacetic acid (TFA). UPLC-MS analyses were performed on a Waters Acquity UPLC apparatus equipped with a Luna Omega PSC18 Column (1.5 µM, 2.1 x 50 mm) coupled to a single quadrupole EDI-MS (Mictomass ZQ). ESI mass spectra were recorded on an LCQESI MS on a LCQ Advantage spectrometer from Thermo Finningan and a LCQ Fleet spectrometer from Thermo Scientific. Optical rotations were measured on a Perkin–Elmer 343 polarimeter at 20°C (concentration [c] in g/100 mL).

**Procedure A**

**1**, **2**, and **3** were synthesized by microwave-assisted Fmoc/tBu-based solid phase peptide synthesis (SPPS) using an automated synthesizer (Liberty Blue, CEM). Rink-Amide resin (0.55 mmol/g loading) was used as solid support and the synthesis was carried out on a 0.1 mmol scale. All used amino acids were *N*-terminally Fmoc-protected, while the side chains of trifunctional amino acids were protected with orthogonal, acid labile groups. The coupling was performed using 5 equivalents (eq.) of the protected amino acid, previously dissolved in DMF to obtain a 0.2 M solution. As coupling reagents, 5 eq. of DIC (0.5 M in DMF) and 5 eq. of Oxyma Pure (1 M in DMF) have been used. To deprotect the Fmoc group, a solution of piperidine in DMF (20% v/v) has been applied. The coupling reaction has been accomplished at 25 °C for 120 s, followed by 480 s at 50 °C and 35 W. The Fmoc group was cleaved with a standard deprotection protocol at 75 °C, 155 W for 15 s followed by 60 s at 90 °C, 50 W. The coupling of the synthetic scaffold was performed directly on resin, using as coupling system DIC/Oxyma Pure/DMF (2/2/eq) and running it overnight at room temperature. Finally, the peptidyl-bound resins were cleaved with a mixture of using 4 mL of acidic cocktail (trifluoracetic acid/water/thioanisol/triisopropyl silane; 95/2.5/1.25/1.25%) for 2 h under continuous shaking.

**Procedure B**

The methyl ester derivative compound (1 eq.) was suspended in dry THF (1M) in a round bottom flask equipped with a magnetic stirrer. Afterward, LiOH 0,1 M in H2O (1.5 eq.) was added, and the reaction was left to stir for

30 min at room temperature. At the end of the reaction, checked by TLC (hexane/EtOAc = 7:3), the solvent was removed by reduced pressure and then the aqueous solution was acidified with 10% HCl (checking the pH with litmus paper till pH < 5) and extracted with ethyl acetate (3 x 15 mL). The combined organic phases were then dried over $Na_2SO_4$, and the solvent was removed under reduced pressure, affording a white solid in quantitative yield.

**Procedure C**

In a round bottom flask equipped with a magnetic stirrer, the carboxylic acid (1 eq.) was suspended in dry DCM (0.1M), then the solution was cooled to 0 °C. Afterward, HCl·H-AA-OMe (1.5 eq.) and Propanephosphonic acid anhydride (T3P) solution 50% in EtOAc (3 eq.) were added. Finally, DIPEA was added until pH = 8 (generally 5/6 eq.). The reaction was stirred overnight at room temperature. After 12 hours, the reaction mixture was successively washed with 5% aqueous solution of $KHSO_4$ (20 mL), aqueous solution of $NaHCO_3$ (20 mL), and brine (25 mL). The isolated organic layer was dried over $Na_2SO_4$ and concentrated under reduced pressure, and the crude product was purified by flash chromatography with the appropriate eluants.

**Procedure D**

In a round bottom flask equipped with a magnetic stirrer, azido derivative compound (1 eq.) was dissolved in THF (0.1M). Afterward, $H_2O$ (7 eq.) and $PMe_3$ 1M in Toluene (1.1 eq.) were added to the solution. The reaction mixture turned from colourless to matt white. The reaction was stirred for 16h at room temperature. At the end of the reaction, checked by TLC (hexane/EtOAc = 7:3) and stained with ninhydrin, the mixture was filtered over cotton to remove a part of the white solid of trimethylphosphine-oxide formed during the reaction. After concentration under reduced pressure, the isolated amine obtained was directly in the next step.

**Procedure E**

In a round bottom flask equipped with a magnetic stirrer, the amine derivative compound (1 eq.) was dissolved in DCM (0.1M), and the solution was cooled to 0°C. Afterward, Fmoc-OSU (1.1 eq.) was added, and the pH was checked with litmus paper. The pH must be basic, around 8.5, but not higher to limit the Fmoc-cleavage. If it was found to be acid, it was necessary to add DIPEA (1 eq.) and the pH had to be rechecked. The reaction was stirred for 1 h at 0 °C and then for 4 h at r.t. At the end of the reaction, checked by TLC (DCM/MeOH + AcOH = 95:5 + 1%), silica was directly added to the mixture, the solvent was removed and the crude was purified by flash chromatography.

**Circular Dichroism**

Compounds, previously pre-treated with HFIP, were dissolved in MQ water to a concentration of 500 μM as stock solutions. Before measurement, each compound was diluted to 125 μM concentration with 10 mM PB (pH 7.2 or pH 5.1) buffer or $H_2O$ into a cuvette with a path length of 1 mm. The CD spectra were recorded by J-815 spectropolarimeter (JASCO, Tokyo, Japan) from 190 to 260 nm at 20 and 37°C and a scan rate of 50 nm/min (accumulation n=3). Each CD spectrum was corrected by subtracting the corresponding baseline (PB buffer 10 mM).

**FTIR Spectroscopy**

Infrared spectra were recorded using a Shimadzu IRAffinity-1S spectrometer in the 600–4000 $cm^{-1}$ range with a resolution of 2 $cm^{-1}$. A sample for measurements in the solid state was prepared by dissolving the peptidomimetic in PB 10 mM (previously treated with HFIP), placing the solution on the crystal plate, and evaporating the solvent (64 scans were averaged). The ATR FTIR experiments were measured from a solution at 125 µM compound concentration. Transmittance has been recorded and transformed into absorbance [$A = 2 – \log(T)$]. Data processing was performed using solver in excel software (Microsoft). Deconvolution of the spectra was done in the spectral range of 1500–1800 $cm^{-1}$. The deconvoluted spectra were fitted with Gaussian band profiles. The positions and number of the components used as an input file for the curve-fitting function were obtained from the deconvoluted spectra. The quality of the fitting was estimated by the standard deviation and error squared.

**Self-aggregation assay by ThT fluorescence spectroscopy**

Ac-PHF6*-$NH_2$ and compounds **1** and **2** were dissolved in pure hexafluoro-isopropanol (HFIP) at a concentration of 1 mM and incubated for 10 minutes at room temperature to dissolve any preformed aggregates. Next, HFIP was evaporated under a stream of dry nitrogen gas followed by vacuum desiccation for at least 3 hours. The resulting thin film was then dissolved in 20 mM MOPS buffer (pH 7.4), sonicated for 1 min, and vortexed for 2 min to get a fully dispersed clean solution at 500 µM concentration. Thioflavin-T binding assays were used to measure the formation of fibrils in solution using a plate reader (Fluostar Optima, BmgLabtech) and standard 96-wells flat-bottom black microtiter plates (final volume 200 µL, 440 nm of excitation wavelength and 480 nm for emission). ThT assay was started by adding 2 µL of a 100 µM heparin solution (heparin sodium salt H-3149, average MW 18 kDa, final concentration 1 µM) to a mixture containing 25 µM of Ac-PHF6*-$NH_2$ or compounds **1** and **2**, 20 µM ThT in 20 mM MOPS pH 7.4 buffer. Fluorescence data were normalized by putting the max fluorescence of Ac-PHF6* at 100% (time of kinetics 60 min). The results are represented as the average of three replicates (n = 3) and the error bars are indicated as the ±SEM.

**ThT fluorescence spectroscopy on wt-tau$_{441}$**

Lyophilized full-length wt-Tau$_{441}$ was diluted to 40 µM in PB 25 mM, NaCl 25 mM, and EDTA 2.5 mM pH 6.8. Stock solutions of compounds **1**, **2** and **3** were prepared in water as described in the general protocol « Self-aggregation assay by ThT fluorescence spectroscopy ». ThT fluorescence was measured to evaluate the development of Tau fibrils over time using a fluorescence plate reader (Fluostar Optima, BMG labtech) with 384-wells flat-bottom black plates (final volume in the wells of 40 µL). Experiments were conducted with and without the addition of heparin (final concentration 0.1 µM) and with or without compounds 1 and 2 (final concentration 1 µM) and with different concentrations of compound **3** (50 µM, 10 µM, and 1 µM). The final concentration of tau$_{441}$ was maintained at 10 µM and that of Thioflavin-T at 25 µM. The ThT fluorescence intensity of each sample (performed in triplicate) was recorded every 10 min (440/480 nm excitation/emission) during 140 h or 40 h under continuous agitation (orbital shaking) at 37 °C on plates sealed with a transparent film. The kinetic curves represent the average of measurements made in triplicate from two different experiments and the error bars are indicated as the ±SEM.

**Transmission Electron Microscopy (TEM)**

Wt-Tau$_{441}$ was dissolved in PB buffer ($Na_2HPO_4$ and $NaH_2PO_4$ 25 mM, NaCl 25 mM, EDTA 2.5 mM, pH 6.6) to a final concentration of 40 µM (stock solution). Sample preparation: Experiments were started by adding 10 µL of buffer, 10 µL of Tau solution (final concentration of 10 µM), 10 µL of compound **1**, **2** or **3** (either 200 µM or 40 µM or 4 µM to have 5:1, 1:1 and 0.1:1 ratios, respectively), and, when necessary, 10 µL heparin solution solubilized in NaPi buffer (final concentration 0.1 µM). In blank analysis, 10 µL of active compound was replaced by 10 µL MQ $H_2O$ and 10 µL heparin solution by 10 µL of buffer. These solutions were maintained at 37°C for 96 h at 1400 rpm.

The samples were prepared on carbon-coated copper grids (ECF200- Cu, 200 mesh, Science Services, Munich, Germany). The grids were pre-treated with argon in a plasma cleaner (Diener Electronics, Ebhausen, Germany). The sample (1.2 µL) was applied onto the grids, and after a sedimentation time of 5 minutes, the excess suspension was removed with filter paper. Grids were then stained with 1% acetate uranyl solution (1.2 µL) for 5 minutes, excess stain was removed with filter paper and washed thoroughly with 3 x 10 µL of Milli-Q water. The prepared grids were analyzed with a JEOL JEM-2200FS electron microscope (JEOL, Freising, Germany), a cold field emission electron gun, and an applied acceleration voltage of 200 kV. A bottom-mounted Gatan OneView camera (Gatan, Pleasanton, CA, USA) was used for digital recording. The images were processed using the image-processing system Digital Micrograph GMS3 (Gatan, Pleasanton, CA, USA) and the image editing software ImageJ.

**Metabolic activity and cytotoxicity profiling using a combined LDH and MTT Assays**

PC12 cells were cultured in 96-well plates at $1\times10^4$ cells/well in 50 µl of serum-reduced medium supplemented with 100 ng/ml 7S mouse NGF to induce neuronal differentiation. Cells were incubated for 48, test compounds were added in an additional volume of 50 µl, and incubation was continued for 20 hours. For the LDH assay, 50 µl medium from each well of the assay plate was transferred to a separate 96-well plate. To quantify LDH release, 50µl of LDH reagent (4 mM iodonitrotetrazolium chloride (INT), 6.4 mM beta-nicotinamide adenine dinucleotide sodium salt (NAD), 320 mM lithium lactate, 150 mM of 1-methoxyphenazine methosulfate (MPMS) in 0.2 M Tris-HCl buffer, pH 8.2) was added. Absorbance was measured at 490 nm using a Thermomax Microplate Reader operated with SoftMaxPro Version 1.1 (Molecular Devices Corp., Sunnyvale, CA, U.S.A.). For the MTT assay, MTT reagent (3,(4,5.dimethylthiazol-2-yl)2,5-diphenyltetrazolium bromide) was added to the wells of the remaining assay plate at a final concentration of 1 mg/ml MTT. Cells were incubated for 2 hours before the reaction was stopped by adding 50 µl of lysis buffer (20% (wt/vol) sodium dodecyl sulfate in 1:1 (vol/vol) N,N-dimethylformamide/water, pH 4.7). After overnight incubation at 37 °C, optical densities of the formazan product were determined at 570 nm. MTT conversion measurements were normalized to the optical densities of negative control wells. All experiments were performed in triplicates on two independent plates.

**Live-cell imaging and Fluorescence Decay After Photoactivation (FDAP)**

FDAP experiments were performed essentially as previously described.[1] Briefly, cells expressing TauΔK280 were plated on 35-mm glass-bottom culture dishes (MatTek, USA), transfected, and neuronally differentiated by medium exchange to serum-reduced DMEM containing 100 ng/mL 7S mouse NGF. After 3 days, the medium was exchanged to serum-reduced DMEM without phenol red with NGF, and the respective compound (or DMSO for carrier control) was added at the desired concentration. After 20 hours, live cell imaging was performed using a laser scanning microscope (Nikon Eclipse Ti2-E (Nikon, Japan)) equipped with a LU-N4 laser unit with 488-nm and 405-nm lasers and a Fluor 60× ultraviolet-corrected objective lens (NA 1.4) enclosed in an incubation chamber at 37°C and 5% $CO_2$. Photoactivation was performed with a 405-nm laser using the microscope software (NIS-

Elements version AR 5.02.03 (Nikon, Japan)). A series of consecutive images were acquired at a frequency of 1 frame/s, and 112 images were collected per activated cell at a resolution of 256×256 pixels. Effective diffusion constants were determined by fitting the fluorescence decay data from the photoactivation experiments using a one-dimensional diffusion model function for FDAP. A reaction-diffusion model was used to estimate the association rate $k^*_{on}$ and the dissociation rate $k_{off}$ constant of tau binding.

### Characterization of the synthetic intermediates and the final compounds 1 and 2

***Compound 1:*** *(S)-N[1]-((4S,7S,10S,13S,14S)-7-(2-amino-2-oxoethyl)-1-((S)-5-(((3S,6S,9S,12S,15S,18S)-22-amino-6-(3-amino-3-oxopropyl)-9-((S)-sec-butyl)-18-carbamoyl-15-(4-hydroxybenzyl)-12-isopropyl-2-methyl-4,7,10,13,16-pentaoxo-5,8,11,14,17-pentaazadocosan-3-yl)carbamoyl)-3-phenyl-4,5-dihydroisoxazol-5-yl)-4-(4-aminobutyl)-10-((S)-sec-butyl)-14-methyl-3,6,9,12-tetraoxo-2,5,8,11-tetraazahexadecan-13-yl)-2-((S)-2-amino-3-methylbutanamido)pentanediamide*

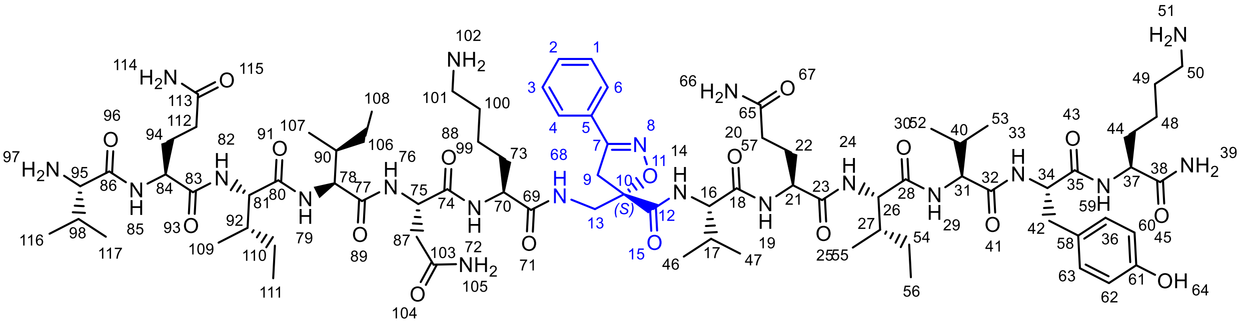

Peptidomimetic **1** was synthesized accordingly to **General procedure A**.

The peptide was purified by semi-preparative HPLC (Linear gradients of 20-70% ACN in $H_2O$ containing 0.1% TFA in 20 min, yield isolation: 50%).

**Molecular weight**: 1644.9715 g/mol

**HRMS**: Calcd. for $[C_{79}H_{128}N_{20}O_{18} + H]^+$: m/z 1645.9788  found 823.4954 $[M+2H]^{2+}$ and 834.4862 $[M+H+Na]^{2+}$

**HPLC purity**: XSELECT column (C18, 2.1 x 75mm-2.5μm); (Linear gradients of 5-100% ACN in $H_2O$ containing 0.1% TFA in 20 min); Rt = 4.792 min, 100 %.

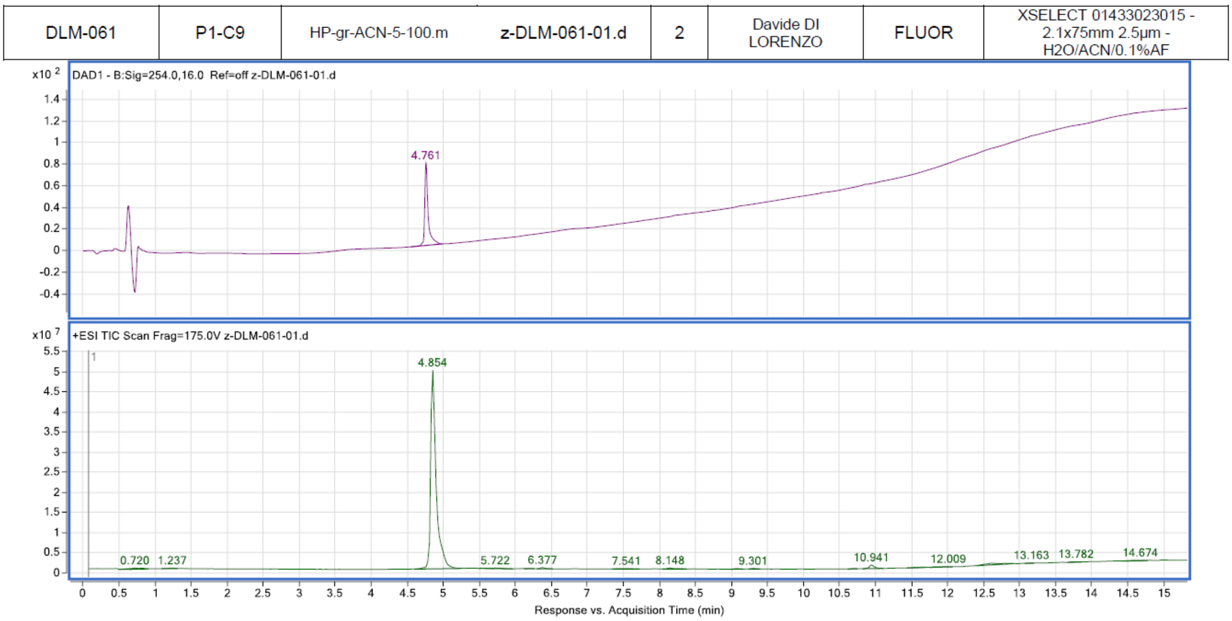

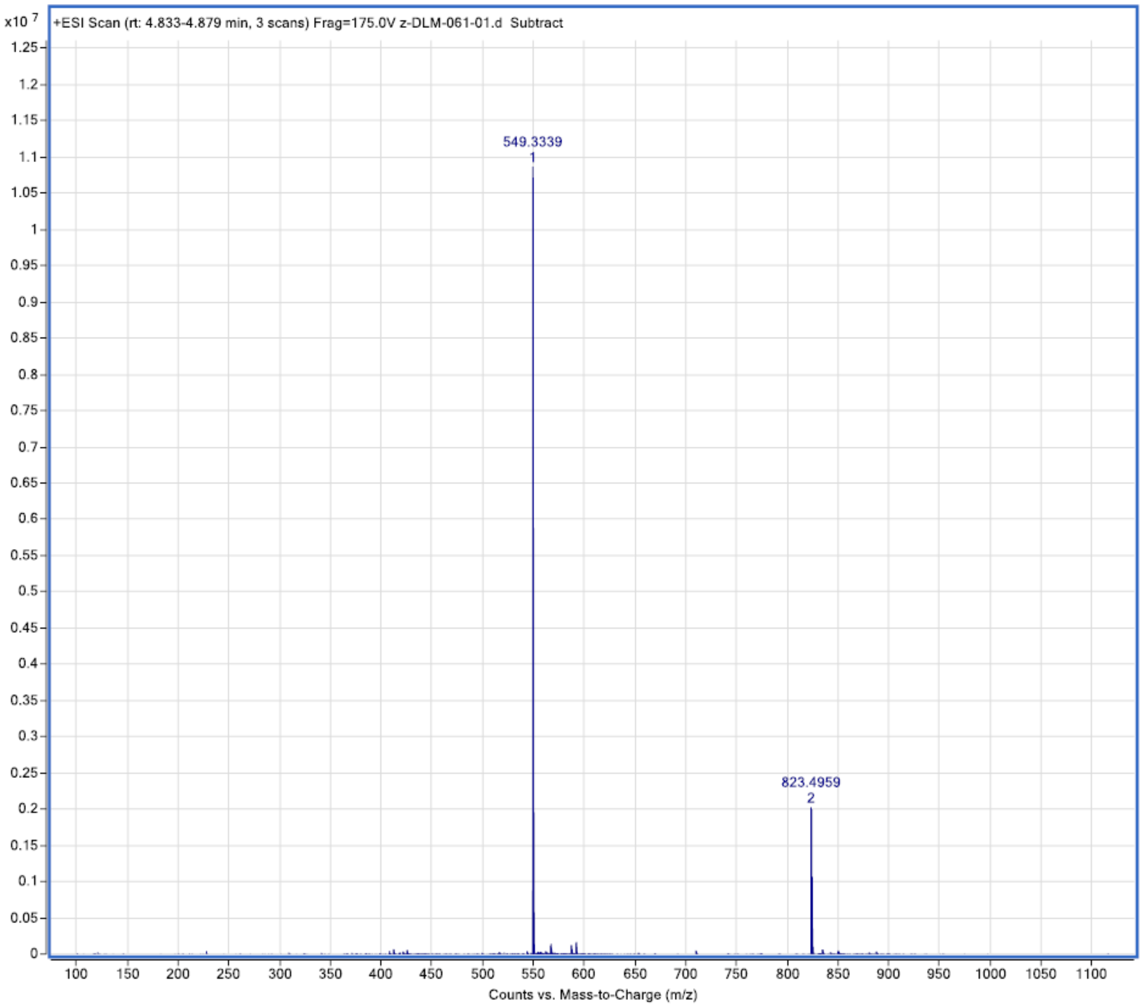

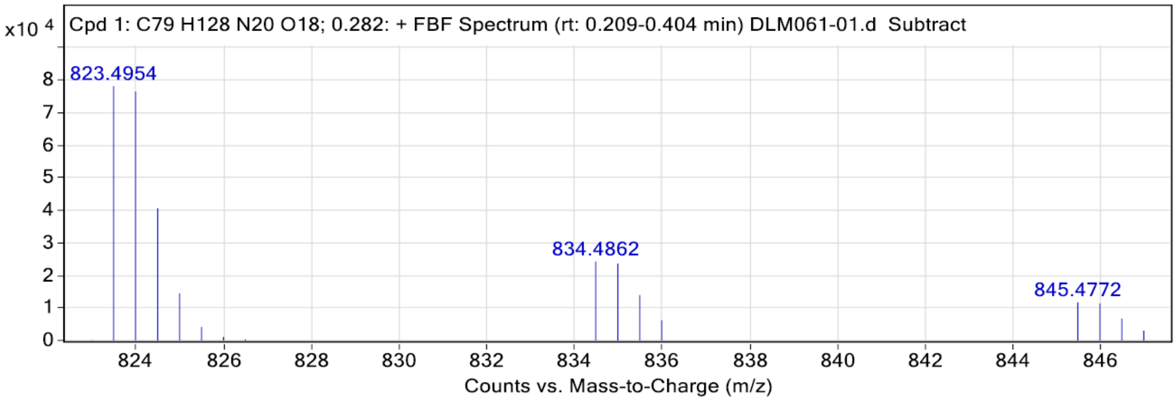

**Peak List**

| *m/z* | z | Abundance | Target Formula | Target Ion Species | Calculated m/z | Diff (mDa) | Diff (ppm) |
|---|---|---|---|---|---|---|---|
| 823,4954 | 2 | 78101,54 | C79H128N20O18 | (M+2H)+2 | 823,4931 | 2,38 | **2,89** |
| 823,9967 | 2 | 76445,04 | C79H128N20O18 | (M+2H)+2 | 823,9945 | 2,19 | **2,66** |
| 824,4980 | 2 | 40560,20 | C79H128N20O18 | (M+2H)+2 | 824,4959 | 2,07 | **2,51** |
| 834,4862 | 2 | 24212,14 | C79H128N20O18 | (M+H+Na)+2 | 834,4840 | 2,20 | **2,64** |
| 834,9871 | 2 | 23555,83 | C79H128N20O18 | (M+H+Na)+2 | 834,9855 | 1,62 | **1,94** |

***Compound 2:*** *(S)-$N^1$-((4S,7S,10S,13S,14S)-7-(2-amino-2-oxoethyl)-1-((R)-5-(((3S,6S,9S,12S,15S,18S)-22-amino-6-(3-amino-3-oxopropyl)-9-((S)-sec-butyl)-18-carbamoyl-15-(4-hydroxybenzyl)-12-isopropyl-2-methyl-4,7,10,13,16-pentaoxo-5,8,11,14,17-pentaazadocosan-3-yl)carbamoyl)-3-phenyl-4,5-dihydroisoxazol-5-yl)-4-(4-aminobutyl)-10-((S)-sec-butyl)-14-methyl-3,6,9,12-tetraoxo-2,5,8,11-tetraazahexadecan-13-yl)-2-((S)-2-amino-3-methylbutanamido)pentanediamide*

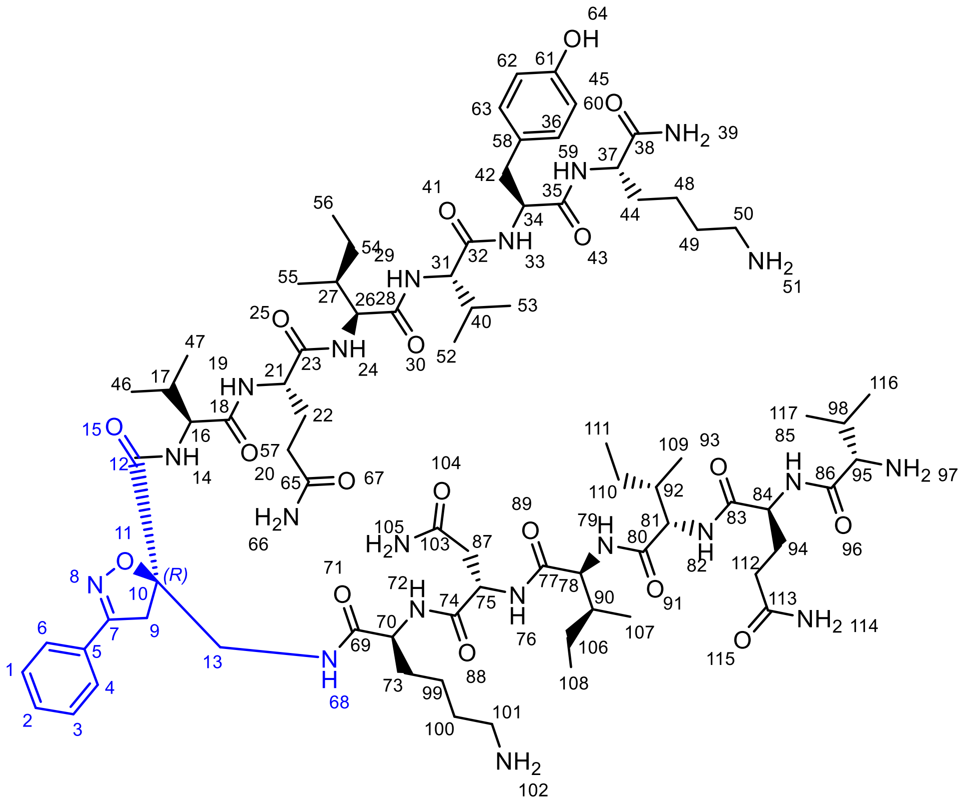

Peptidomimetic **2** was synthesized according to **General Procedure A**.

The peptide was purified by semi-preparative HPLC (Linear gradients of 20-70 % ACN in $H_2O$ containing 0.1% TFA in 20 min, yield isolation: 50%).

**Molecular weight:** 1644.9715 g/mol

**HRMS:** Calcd. for $[C_{79}H_{128}N_{20}O_{18} + H]^+$: m/z 1645.9788 found 1645.9816 $[M + H]^+$; Calcd. for $[C_{79}H_{129}N_{20}O_{18} + Na]^+$: m/z 1667.9608 found 1667.9594 $[M + Na]^+$ and 823.4954 $[M + 2H]^{2+}$

**HPLC purity**: XSELECT column (C18, 2.1 x 75mm-2.5µm); (Linear gradients of 5-100% ACN in $H_2O$ containing 0.1% TFA in 20 min); Rt = 5.043 min, 100 %.

| | | | | | | | |
|---|---|---|---|---|---|---|---|
| DL M 059 A | P1-C4 | HP-gr-ACN-5-100-HighMass.m | z-DL-M-059A-01.d | 2 | Davide DI LORENZO | FLUOR | XSELECT 01433023015 - 2.1x75mm 2.5µm - H2O/ACN/0.1%AF |

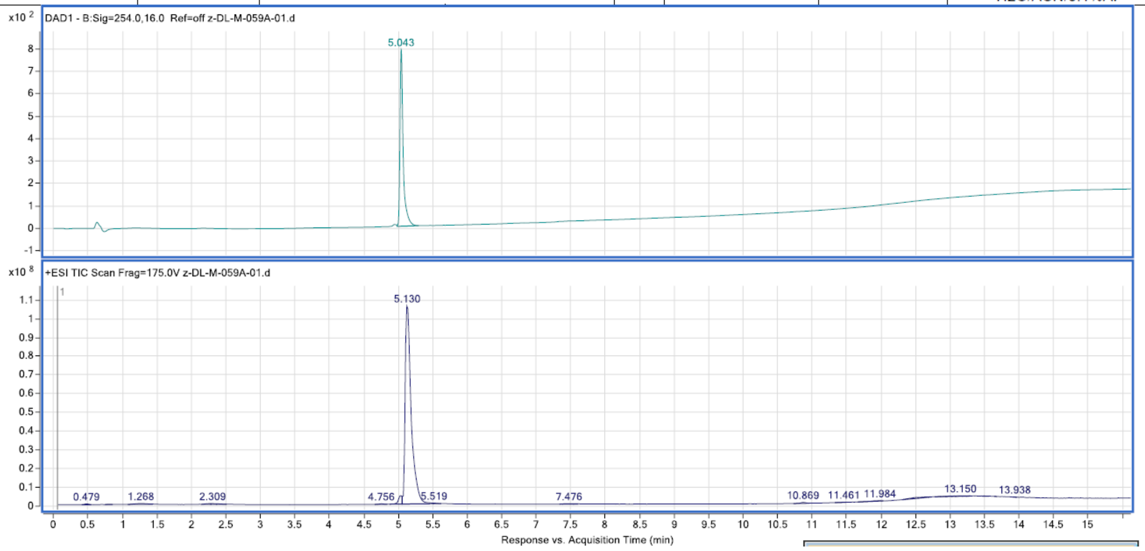

Response vs. Acquisition Time (min)

Peaks: DAD1 - B:Sig=254.0,16.0 Ref=off

| RT | Area | Area Sum % | Height |
|---|---|---|---|
| 5.043 | 2561.98 | 100 | 789.63 |

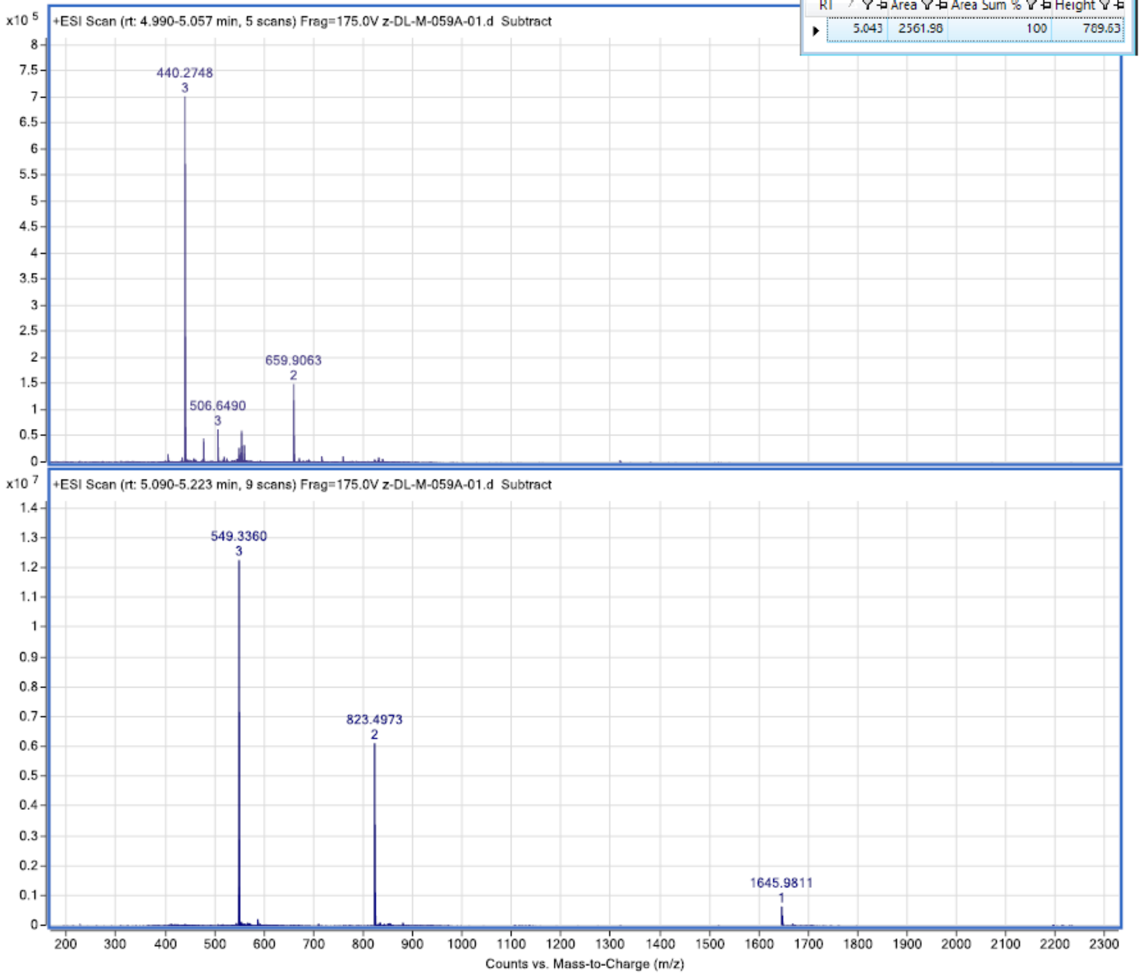

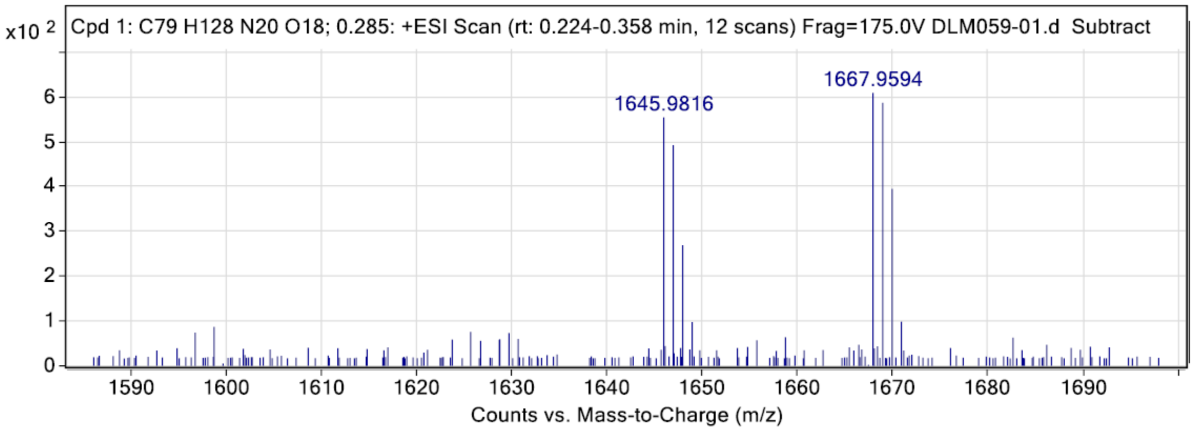

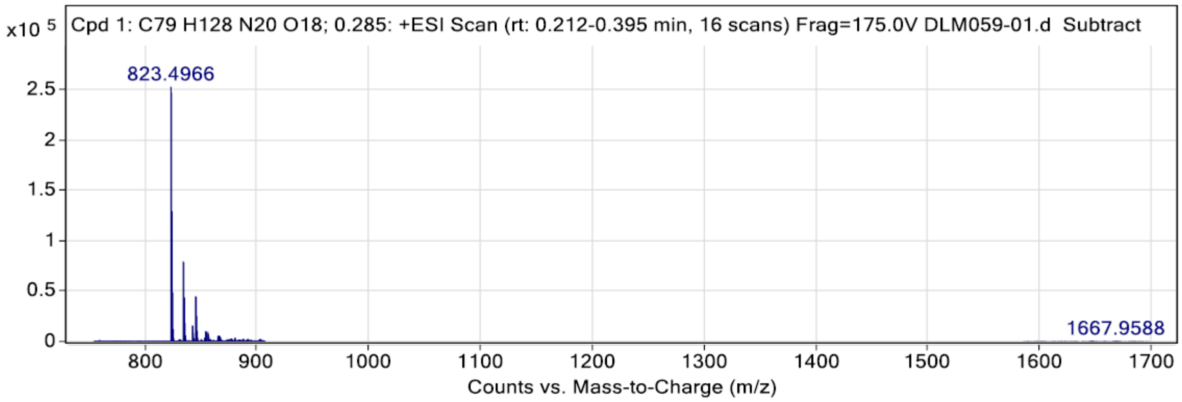

**Peak List**

| *m/z* | z | Abundance | Target Formula | Target Ion Species | Calculated m/z | Diff (mDa) | Diff (ppm) |
|---|---|---|---|---|---|---|---|
| 823,4966 | 2 | 252812,27 | C79H128N20O18 | (M+2H)+2 | 823,4931 | 3,58 | **4,35** |
| 823,9978 | 2 | 247020,44 | C79H128N20O18 | (M+2H)+2 | 823,9945 | 3,28 | **3,99** |

***Compound 5:*** *Methyl 2-(azidomethyl)acrylate*

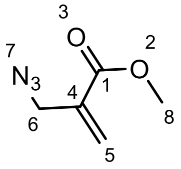

**3.6**

In a round bottom flask equipped with a magnetic stirrer, **4** (74 mg, 4.13 mmol) was suspended in a mixture of $(CH_3)_2CO$ and $H_2O$ 3:1 (16 mL). $NaN_3$ (537 mg, 8.26, 2 eq.) was added to the solution. The reaction was stirred for 4 hours at room temperature. The reaction color turned slowly from colourless to pale orange. At the end of the reaction, the mixture was diluted with $CH_2Cl_2$ (15 mL) and the organic layer was extracted and washed successively with $H_2O$ (5 mL), brine (5 mL) and then dried over $Na_2SO_4$. The solvent was removed under reduced pressure, affording **5** as a pale-yellow oil with 98% yield.

***Rf*** (hexane/AcOEt 9:1) = 0.44

**$^{1}$H NMR (300 MHz, CDCl3)**: δ 6.38-6.35 (1H, m, H5’), 5.85-5.83 (1H, m, H5’’), 4.04 (2H, s, H6), 3.79 (3H, s, H8);

**$^{13}$C NMR: (75 MHz, CD3Cl)**: δ 165.7 (C1), 134.9 (C4); 128.1 (C5); 52.2 (C8); 51.4 (C6).

The NMR analysis is already reported in literature: Sá, M. M., Ramos, M. D. & Fernandes, L. Fast and efficient preparation of Baylis Hillman-derived (E)-allylic azides and related compounds in aqueous medium. *Tetrahedron* 62, 11652–11656, **2006**.

***Compound 7:*** *N-hydroxybenzimidoyl chloride*

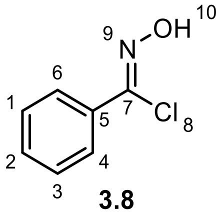

**3.8**

In a two-necked round-bottom flask, equipped with magnetic stirrer and nitrogen inlet, oxime **6** (500 mg, 4.13 mmol) was suspended in dry DMF (6 mL). Afterwards, NCS (551 mg, 4.13 mmol, 1 eq.) was added to the solution. The mixture turned quickly from colourless to bright yellow, and finally pale yellow. The reaction was stirred for 4 hours at room temperature under nitrogen atmosphere.

The reaction mixture was diluted with $CH_2Cl_2$ (15 mL) and the organic layer was extracted and washed with $H_2O$ (10 mL), then dried over $Na_2SO_4$. The solvent was removed under reduced pressure to afford **7** in quantitative yield as a pale-yellow oil, which was used in the next step without further purification.

The NMR analysis are already reported in literature:

**$^{1}$H-NMR (300 MHz, $CDCl_3$)**: δ 7.85-7.35 (5H, m, H1, H2, H3, H4, H6), 8.43 (1H, s, H10).

Tran, N. C., Dhondt, H., Flipo, M., Deprez, B. & Willand, N. Synthesis of functionalized 2-isoxazolines as three-dimensional fragments for fragment-based drug discovery. *Tetrahedron Lett.* 56, 4119–4123, **2015**.

***Compound 9:*** *Methyl 5-(azidomethyl)-3-phenyl-4,5- dihydroisoxazole-5-carboxylate*

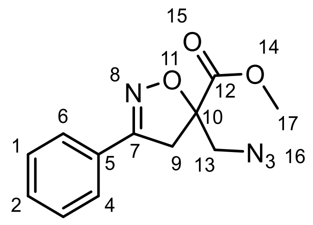

**3.9**

In a two-necked round-bottom flask equipped with a magnetic stirrer and nitrogen inlet, **5** (640 mg, 4.13 mmol, 1 eq.) was suspended in dry THF (5 mL). Afterward, **7** (580 mg, 4.13 mmol, 1 eq.) was diluted in dry THF (7 mL) and added dropwise to the solution. TEA (1,15 mL, 8.26 mmol, 2 eq.) was added dropwise to the mixture, and a white solid precipitate was formed. The reaction was stirred overnight at room temperature under a nitrogen atmosphere. After the concentration of the mixture under reduced pressure, the residue obtained was taken up with AcOEt (10 mL) and washed with $H_2O$ (15 mL). The organic layer was then dried over $Na_2SO_4$, filtered, and the solvent removed under reduced pressure to give a dark yellow oil, which was purified by chromatography on silica gel eluting with Hexane/AcOEt 8:2 to afford **9** (859 mg, 3.304 mmol, 80%) as a white solid.

**Molecular weight**: 260.09 g/mol

***Rf*** (hexane/AcOEt 8:2) = 0.4

**MS**: calcd. for $[C_{12}H_{12}N_4O_3 + H]^+$: m/z 261.0982, found: 261.09 $[M + H]^+$ and $[C_{12}H_{12}N_4O_3 + Na]^+$: m/z 283.0802; found: 283.1 $[M + Na]^+$

**$^1$H NMR (300 MHz, CDCl₃)** : δ 7.70-7.65 (2H, m, H6, H4), 7.54-7.35 (3H, m, H1, H2, H3), 3.87 (3H, s, H17), 3.83-3.76 (2H, m, H13’, H9’), 3.67 (1H, d, *J* = 13 Hz, H13’’), 3.52 (1H, d, *J* = 17.2 Hz, H9’’).

**$^{13}$C NMR (75 MHz, $CDCl_3$)** : δ 170.1 (C12), 156.4 (C7), 130.8 (C2), 128.9 (C3, C1) , 128.4 (C5), 127.0 (C4, C6), 87.6 (C10), 53.4 (C17), 44.2 (C13), 41.5 (C9).

***Compound 9a:*** *5-(Azidomethyl)-3-phenyl-4,5-dihydroisoxazole-5-carboxylic acid*

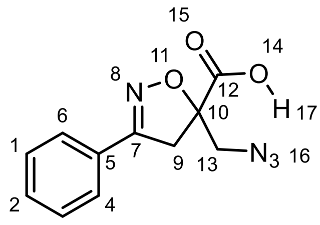

**3.10**

**9a** was obtained as a white solid in quantitative yield following the general procedure **B** starting from **9** (980mg, 3.77 mmol)

**Molecular weight**: 246.08 g/mol

***Rf*** (Hex/EtOAc 7/3) = 0

**MS**: Calcd. for $[C_{11}H_{10}N_4O_3 + H]^+$: m/z 247.0826 found: 247.08 $[M + H]^+$

**$^{1}$H NMR (300 MHz, CDCl$_3$)**: δ 9.77 (s, 1H, H17), 7.73–7.61 (2H, m, H6, H4), 7.54 –7.38 (3H, m, H1, H2, H3), 3.85 (1H, d, *J* = 13.11 Hz, H13'), 3.77–3.67 (2H, m, H13'', H9'), 3.58 (1H, d, *J* = 17,38 Hz, H9'').

**$^{13}$C NMR (75 MHz, CDCl$_3$)**: δ 173.60 (C12), 156.91 (C7), 131.03 (C2), 128.93 (C3, C1), 127.89 (C5), 127.03 (C6, C4), 87.52 (C10), 43.97 (C13), 41.75 (C9).

***Compound 10a :*** *Methyl ((S)-5-(azidomethyl)-3-phenyl-4,5-dihydroisoxazole-5-carbonyl)-L-valinate*

***Compound 10b:*** *Methyl ((R)-5-(azidomethyl)-3-phenyl-4,5-dihydroisoxazole-5-carbonyl)-L-valinate*

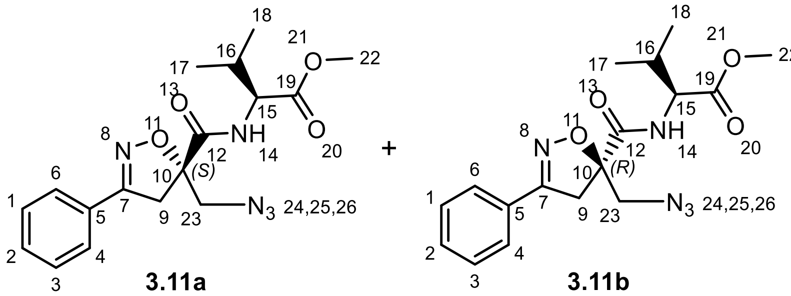

**10a** and **10b** were obtained following the general procedure **C** starting from **9a** (927 mg, 3.77 mmol). The crude product was purified by chromatography on silica gel eluting with DCM/Et$_2$O 98:2 to yield **10a** and **10b** (1.3 g, 3.66 mmol, overall yield 97%) as white solids isolated in 42% and 47% yield, respectively.

**Molecular weight**: 359.16 g/mol

**(10a) R*f*** (DCM/Et$_2$O = 98/2) = 0.3

**(10a) [α]$_{20}^{D}$**: +15° ([c] = 0.001 in CH$_2$Cl$_2$)

**(10b) R*f*** (DCM/Et$_2$O = 98/2) = 0.2

**(10b) [α]$_{20}^{D}$**: -18.8° ([c] = 0.005 in CH$_2$Cl$_2$)

**MS**: Calcd. for [C$_{17}$H$_{21}$N$_5$O$_4$ + H]$^+$: m/z 360.1666; found: 359.98 [M + H]$^+$ and 740.83 [2M + Na]$^+$

**(10a) $^{1}$H NMR (300 MHz, CDCl$_3$)**: δ 7.68 (2H, m, H6, H4), 7.46, 7.44 (3H, m, H1,H2,H3), 7.35 (1H, d, *J* = 8.9 Hz, 1H, NH14); 4.54 (1H, dd, *J* = 9.0 Hz and 5.0 Hz, H15), 3.84 (1H, d, *J* = 13.0 Hz, H23'), 3.71 (3H, s, H22), 3.70 (1H, d, J = 17.53 Hz, H9'), 3.67 (1H, d, *J* = 13.1 Hz, H23''), 3.52 (1H, d, *J* = 17.53 Hz, H9''), 2.26 (1H, m, H16), 1.01 (6H, m, H17, H18).

**(10a) $^{13}$C NMR (101 MHz, CDCl$_3$)**: δ 171.35 (C19), 170.47 (C12), 157.47 (C7), 130.95 (C2), 128.89 (C1, C3), 128.18 (C5), 127.00 (C4, C6), 89.20 (C10), 57.34 (C15), 55.14 (C23), 52.24 (C22), 42.42 (C9), 31.20 (C16), 18.98 (C18), 17.72 (C17).

**(10b) $^{1}$H NMR (400 MHz, CDCl$_3$)**: δ 7.75–7.61 (2H, m, H6, H4), 7.5 –7.41 (3H, m, H1, H2, H3), 7.38 (1H, d, *J* = 9.1 Hz, H14), 4.50 (1H, dd, *J* = 9.1Hz and 5.1 Hz, H15), 3.88 (1H, d, *J* = 13.3 Hz, H23'), 3.65 (3H, s, H22), 3.70 (1H, d, J = 17.49 Hz, H9'); 3.65 (1H, d, *J* = 13.3 Hz, H23''); 3.62 (1H, d, *J* = 17.45 Hz, H9''), 2.29–2.11 (1H, m, H16), 0.90 (6H, d, *J* = 6.9 Hz, H17, H18).

**(10b) $^{13}$C NMR (101 MHz, $CDCl_3$)**: δ 171.34 (C19), 170.76 (C12), 157.77 (C7), 130.99 (C2), 128.94 (C1, C3), 128.21 (C5), 126.97 (C6, C4), 89.39 (C10), 57.30 (C15), 54.29 (C23), 52.33 (C22), 42.01 (C9), 31.05 (C16), 18.97 (C17), 17.69 (C18).

***Compound 11a:*** *((S)-5-(Azidomethyl)-3-phenyl-4,5-dihydroisoxazole-5-carbonyl)-L-valine*

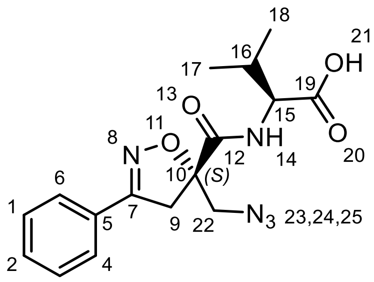

**3.12a**

**11a** (193.2 mg, 0.56 mmol, quant.) was obtained as a white solid following the general procedure **B** from **10a** (201 mg, 0.56 mmol).

**Molecular weight**: 345.14 g/mol

White solid

**R*f*** (Hex/EtOAc 7/3) = 0

**MS**: Calcd. for $[C_{16}H_{18}N_4O_3 - H]^-$: m/z 344.1359; found: 344.44 $[M - H]^-$

**$^1$H NMR (300 MHz, $CDCl_3$)**: δ 7.71 (1H, bs, H21), 7.65–7.59 (2H, m, H6, H4), 7.48–7.34 (4H, m, H1, H2, H3, H14), 4.49 (1H, dd, *J* = 8.9 Hz and 4.7 Hz, H15), 3.81 (1H, d, *J* = 13.1 Hz, H22'), 3.68 (1H, d, *J* = 17.6 Hz, H9'), 3.62 (1H, d, *J* = 13.05 Hz, H22''), 3.46 (1H, d, *J* = 17.6 Hz, H9''), 2.34–2.20 (1H, m, H16), 0.99 (6H, m, H17, H18).

**$^{13}$C NMR (75 MHz, $CDCl_3$)**: δ 175.17 (C19), 170.78 (C12), 157.65 (C7), 131.01 (C2), 128.89 (C1, C3), 127.96 (C5), 127.01 (C6, C4), 89.13 (C10), 57.20 (C15), 55.04 (C22), 42.37 (C9), 30.90 (C16), 18.98 (C17), 17.53 (C18).

***Compound 11b:*** *((R)-5-(Azidomethyl)-3-phenyl-4,5-dihydroisoxazole-5-carbonyl)-L-valine*

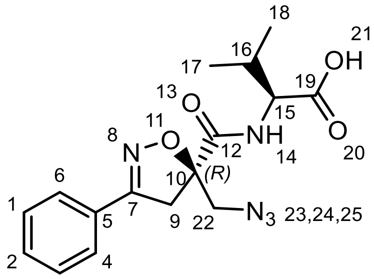

**3.12b**

**11b** (548.8 mg, 1.59 mmol, quant.) was obtained as a white solid following the general procedure **B** from 10b. Compound 11b was crystallized by slow evaporation from a solution containing Et2O/Hexane. 11b was dissolved in the lowest amount of Et2O, and a small amount of hexane was added until the appearance of turbidity. After an extremely low evaporation of the solution, without moving the flask, crystals of **11b** were obtained.

**Molecular weight**: 345.14 g/mol;

***Rf*** (Hex/EtOAc 7/3) = 0

**[α]$_{20}$$^{D}$ (3.12b)**: + 6° ([c] = 0.004 in $CH_2Cl_2$);

**MS**: Calcd. for [$C_{16}H_{19}N_4O_3$ - H]$^-$: m/z 344.1359; found: 344.44 [M - H]$^-$

**$^1$H NMR (400 MHz, $CDCl_3$)**: δ 8.58 (1H, bs, H21), 7.69 (2H, dd, *J* = 7.9 Hz and 1.8 Hz, H6, H4), 7.56–7.19 (4H, m, H1, H2, H3, H14), 4.53 (1H, dd, *J* = 8.9 Hz and 4.9 Hz, H15), 3.90 (1H, d, *J* = 13.3 Hz, H22'), 3.72 (1H, d, *J* = 17.5 Hz, H9'), 3.67 (1H, d, *J* = 13.4 Hz, H22''), 3.64 (1H, d, *J* = 17.5 Hz, H9''), 2.30 (1H, m, *J* = 6.9 Hz and 4.9 Hz, H16), 0.96 (6H, dd, *J* = 6.9, 3.8 Hz, H17, H18) ppm.

**$^{13}$C NMR (101 MHz, $CDCl_3$)**: δ 175.48 (C19), 171.21 (C12), 157.91 (C7), 131.06 (C2), 128.96 (C1, C3), 128.12 (C5), 127.01 (C6, C4), 89.37 (C10), 57.25 (C15), 54.23 (C22), 41.97 (C9), 30.73 (C16), 19.05 (C17), 17.58 (C18) ppm.

***Compound 11a-NH$_2$:*** *Methyl ((S)-5-(aminomethyl)-3-phenyl-4,5-dihydroisoxazole-5-carbonyl)-L-valine*

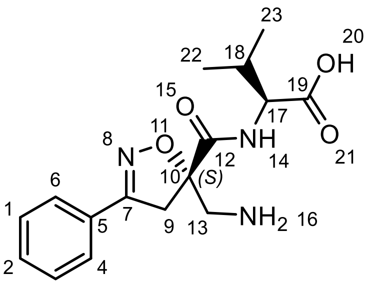

**3.13a**

**11a-NH$_2$** was obtained as a pale yellow oil following the general procedure **D** starting from **11a** (193 mg, 0.56 mmol). The compound was used for the next synthetic step without any further purification.

**Molecular weight**: 319.15 g/mol;

***Rf*** (Hex/EtOAc 7/3) = 0

**MS**: Calcd. for [$C_{16}H_{21}N_4O_3$ + H]$^+$: m/z 320.1605; found: 320.23 [M + H]$^+$

***Compound 11b-NH$_2$:*** *Methyl ((R)-5-(aminomethyl)-3-phenyl-4,5-dihydroisoxazole-5-carbonyl)-L-valine*

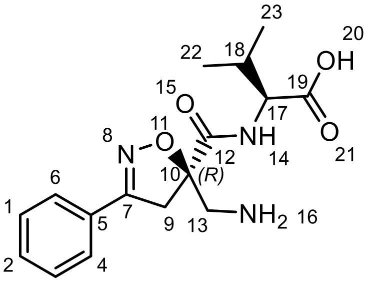

**3.13b**

**11b-NH$_2$** was obtained as a pale yellow oil following the general procedure **D** starting from **11b** (570 mg, 1.59 mmol). The compound was used for the next synthetic step without any further purification.

**Molecular weight**: 319.15 g/mol;

***Rf*** (Hex/EtOAc 7/3) = 0

**MS:** Calcd. for $[C_{16}H_{21}N_4O_3 + H]^+$: m/z 320.1605; found: 320.23 $[M + H]^+$

***Compound 12a:*** *Methyl ((S)-5-(((((9H-fluoren-9-yl)methoxy)carbonyl)amino)methyl)-3-phenyl-4,5-dihydroisoxazole-5-carbonyl)-L-valine*

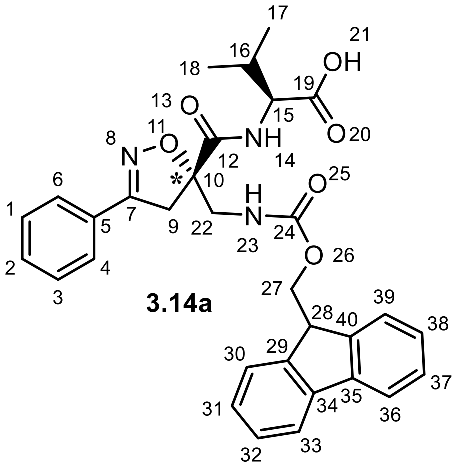

**3.14a**

**12a** was obtained following the general procedure **E** from crude **11a-NH₂** (0.56 mmol). TLC (DCM/MeOH + AcOH = 95:5 + 1%) was used to monitor the reaction. The crude obtained was purified by chromatography on silica gel eluting with DCM/MeOH + AcOH 95/5 + 1% as eluent to afford **12a** (212 mg, 0,392 mmol) as a white spongy solid in 70% yield.

**Molecular weight**: 541.22 g/mol;

**R*f*** (DCM/MeOH + AcOH 95/5 + 1%) = 0.5;

**$[\alpha]_{20}^{D}$ (3.14a)**: -14° ([c] = 0.002 M in $CH_2Cl_2$);

**MS**: Calcd. for $[C_{31}H_{31}N_3O_6 + H]^+$: m/z 542.2286 found: 542.61 $[M + H]^+$; Calcd. for $[C_{31}H_{31}N_3O_6 + Na]^+$: m/z 564.2105; found: 564.79 $[M+Na]^+$ and 1105.90 $[2M + Na]^+$;

**$^1$H NMR (400 MHz, $CD_3OD$)**: δ 7.76 (2H, dd, *J* = 7.6 Hz and 3.6 Hz, H33, H36), 7.72–7.63 (2H, m, H30, H39), 7.59 (2H, d, *J* = 7.5 Hz, H6, H4), 7.51–7.39 (3H, m, H1, H2, H3), 7.34 (2H, t, *J* = 7.5 Hz, H31, H38), 7.22 (2H, dt, *J* = 15.1 Hz and 7.5 Hz, H32, H37), 4.42–4.35 (2H, m, H15, H27), 4.27 (1H, dd, *J* = 10.6 Hz and 6.8 Hz, H27), 4.16 (1H, t, *J* = 6.9 Hz, H28), 3.81–3.64 (3H, m, H9’, H22’, H22’’), 3.55 (1H, d, *J* = 17.8 Hz, H9’’), 2.24 (1H, m, H16), 0.98 (6H, m, H17, H18).

**$^{13}$C NMR (101 MHz, $CD_3OD$)**: δ 172.68 (C19), 172.04 (C12), 157.78 (C24), 157.70 (C7), 143.81 (C40), 143.77 (C29), 141.15 (C34), 141.15 (C35), 130.40 (C2), 128.53 (C5), 128.50 (C1, C3), 127.33 (C6, C4), 126.73 (C38, C31), 126.66 (C32, C37), 124.80 (C39), 124.68 (C30), 119.48 (C33), 119.46 (C36), 89.57 (C10), 66.64 (C27), 57.45 (C15), 46.94 (C28), 45.15 (C22), 41.18 (C9), 30.48 (C16), 18.13 (C18), 16.86 (C17).

***Compound 12b:*** *Methyl ((R)-5-(((((9H-fluoren-9-yl)methoxy)carbonyl)amino)methyl)-3-phenyl-4,5-dihydroisoxazole-5-carbonyl)-L-valine*

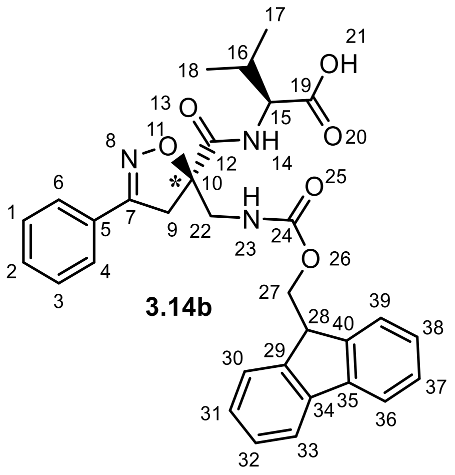

**3.14b**

**12b** was obtained following the general procedure **E** from crude **11b-NH$_2$** (1.59 mmol). TLC (DCM/MeOH + AcOH = 95:5 + 1%) was used to monitor the reaction. The crude obtained was purified by chromatography on silica gel eluting with DCM/MeOH + AcOH 95/5 + 1% as eluent to afford **12b** (500 mg, 0.97 mmol) as a white spongy solid in 60% yield.

**Molecular weight**: 541.22 g/mol;

**R*f*** (DCM/MeOH + AcOH 95/5 + 1%) = 0.5;

**$[\alpha]_{20}^{D}$ (3.14b)**: +20° ([c] = 0.005 in $CH_2Cl_2$);

**MS:** Calcd. for $[C_{31}H_{31}N_3O_6 + H]^+$: M/z 542.2286 found: 542.61 $[M + H]^+$; Calcd. for $[C_{31}H_{31}N_3O_6 + Na]^+$: m/z 564.2105; found: 564.79 $[M + Na]^+$ and 1105.90 $[2M + Na]^+$;

**$^1$H NMR (400 MHz, $CD_3OD$)**: δ 7.75 (2H, dd, *J* = 7.6, 4.4 Hz, H33, H36), 7.72–7.67 (2H, m, H30, H39), 7.60 (2H, dd, *J* = 7.5 Hz and 4.8 Hz, H4, H6), 7.5 –7.39 (3H, m, H1, H2, H3), 7.33 (2H, t, *J* = 7.5 Hz, H31, H38), 7.28–7.14 (2H, m, H32, H37), 4.43– 4.24 (3H, m, H15, H27), 4.18 (1H, t, *J* = 7.1 Hz, H28), 3.80 (1H, d, *J* = 14.6 Hz, H22’), 3.75–3.62 (2H, m, H22’’, H9’’), 3.55 (1H, d, *J* = 17.7 Hz, H9’), 2.23 (1H, m, *J* = 6.9 Hz and 5.2 Hz, H16), 0.92 (6H, m, H17, H18).

**$^{13}$C NMR (101 MHz, $CD_3OD$)**: δ 172.71 (C19), 172.19 (C12), 158.09 (C24), 157.77 (C7), 143.90 (C40), 143.74 (C29), 141.12 (C34, C35), 130.52 (C5), 128.59 (C1, C3), 128.51 (C2), 127.32 (C6, C4), 126.73 (C38, C31), 126.62 (C32, C37), 124.81 (C39), 124.78 (C30), 119.44 (C36, C33), 89.65 (C10), 66.68 (C27), 57.38 (C15), 46.96 (C28), 44.30 (C22), 41.01 (C9), 30.49 (C16), 18.08 (C17), 16.61 (C18).

### Synthetic strategy for compound 3

The two stereoisomers of **9** have been separated after elongating the *C*-terminal peptide sequence. H-Thr(O$^t$Bu)-OMe (1.5 eq) has been coupled with the same protocol based on T3P (3 eq) and DIPEA (3 eq), obtaining **1Sa** and **1Sb** with an excellent 93% yield. However, the diastereomeric mixture could not be separated using silica chromatography. To address this problem, we decided to insert an extra stereogenic center at the *C*-terminal chain. The methyl ester was first removed via basic hydrolysis (LiOH, H2O/THF), followed by the coupling with H-Lys(Boc)-OMe using the T3P protocol. Although the coupling afforded a moderate 28% yield, the resulting diastereoisomers, **2Sa** and **2Sb**, resulted separable by flash chromatography (Hex/EtOAc = 6:4). (**Scheme S1**)

Diastereoisomer **2Sb** was subjected to modification to make it suitable for the SPPS strategy. First, the methyl ester group was hydrolyzed via classical saponification using LiOH (1.5 eq) in THF/$H_2O$, isolating **3Sb** in quantitative yields. Then, the Staudinger protocol was used in the presence of $PMe_3$ (7 eq) and $H_2O$ (49 eq) to reduce the azido group and obtain the amino derivatives **4Sb**, which was then Fmoc-protected (Fmoc-OSu (1.1 eq) in DCM, 4h), affording the final scaffold **5Sb** in satisfactory yields (67%) over two steps. (**Scheme S2**)

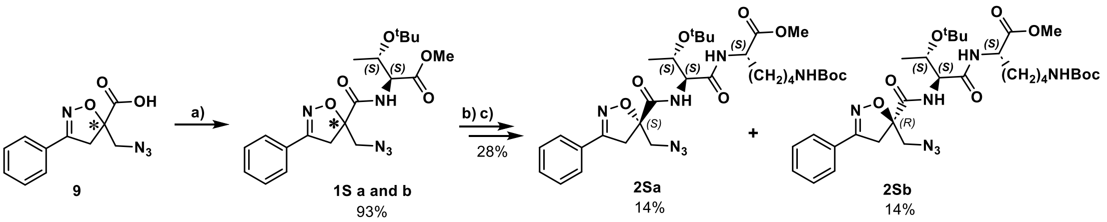

**Scheme S1.** Synthesis and separation of **2Sa** and **2Sb**. **a**) **9** (1 eq), HCl·H-Thr(O$^t$Bu)-OMe (1.5 eq), T3P (3 eq), DIPEA (3 eq), DCM, r.t., 16h; **b**) LiOH 0.1M in $H_2O$ (1.2 eq), THF (1M), r.t., 30 min; **c**) H-Lys(Boc)-OMe (1.5 eq), T3P (3 eq), DIPEA (3 eq), DCM, r.t., 16h.

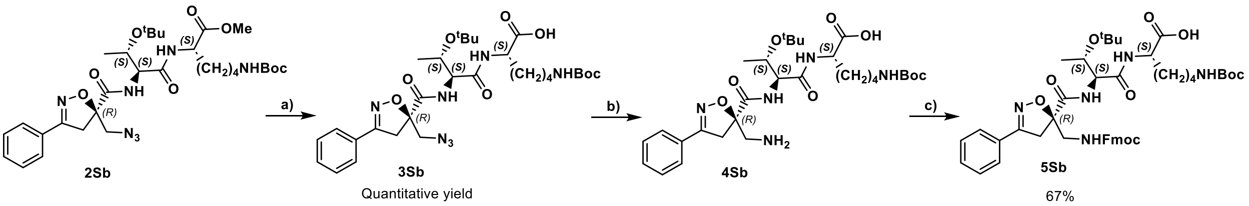

**Scheme S2.** Synthesis of the Fmoc-protected *R*-Isox-$\beta^{2,2}$- Thr(O$^t$Bu)Lys(Boc)-OH (**5Sb**): **a**) LiOH 0.1M in $H_2O$ (1.5 eq), THF (1M), r.t., 30 min; **b**) $PMe_3$ (7 eq), $H_2O$ (49 eq), THF (0.1 M); **c**) Fmoc-OSu (1.1 eq), DCM (0.1 M), 0°C -> r.t., 4h.

***Compound 3:*** *(R)-N-((2S,5S,8S,11S,14S,17S,18S)-2-((1H-imidazol-2-yl)methyl)-1-amino-14-(4-aminobutyl)-8-((S)-sec-butyl)-18-hydroxy-5-isobutyl-11-isopropyl-1,4,7,10,13,16-hexaoxo-3,6,9,12,15-pentaazanonadecan-17-yl)-5-((4S,7S,10S,13S,16S,19S)-19,23-diamino-7-((S)-sec-butyl)-4-((S)-1-hydroxyethyl)-10,13,16-triisopropyl-3,6,9,12,15,18-hexaoxo-2,5,8,11,14,17-hexaazatricosyl)-3-phenyl-4,5-dihydroisoxazole-5-carboxamide*

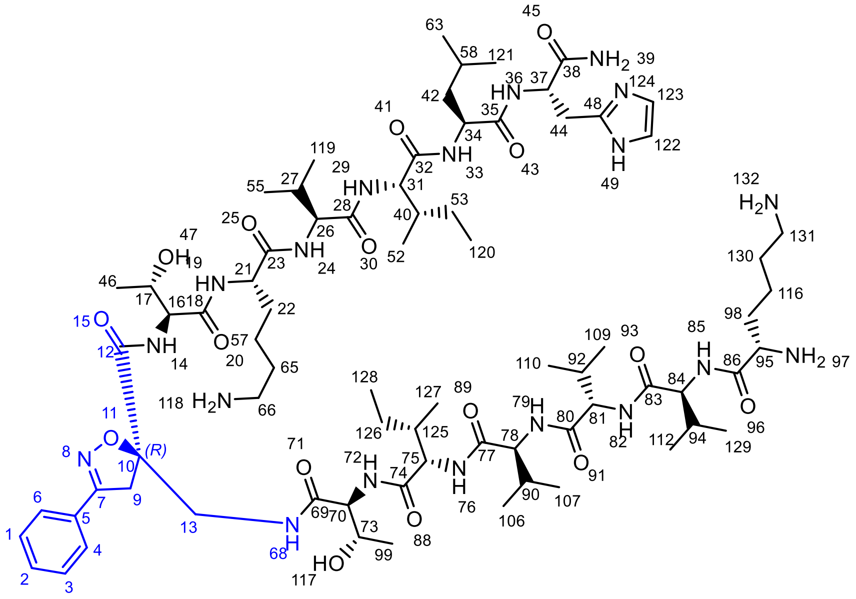

Peptidomimetic **3** was synthesized according to **General Procedure A**.

The peptide was purified by semi-preparative HPLC (Linear gradients of 20-70 % ACN in $H_2O$ containing 0.1% TFA in 20 min, yield isolation: 50%).

**Molecular weight**: 1549.9708 g/mol

**HRMS:** Calcd. for $[C_{75}H_{127}N_{19}O_{16} + H]^+$: m/z 1550.9781 found 1550.9712 $[M + H]^+$; Calcd. for $[C_{79}H_{129}N_{20}O_{18} + Na]^+$: m/z 1572.9600 found 1572.9530 $[M + Na]^+$ and 775,9936 $[M + 2H]^{2+}$ and 517.6668 $[M + 3H]^{3+}$

**HPLC purity**: XSELECT column (C18, 2.1 x 75mm-2.5µm); (Linear gradients of 5-100% ACN in $H_2O$ containing 0.1% TFA in 20 min); Rt = 4.81 min, 100 %.

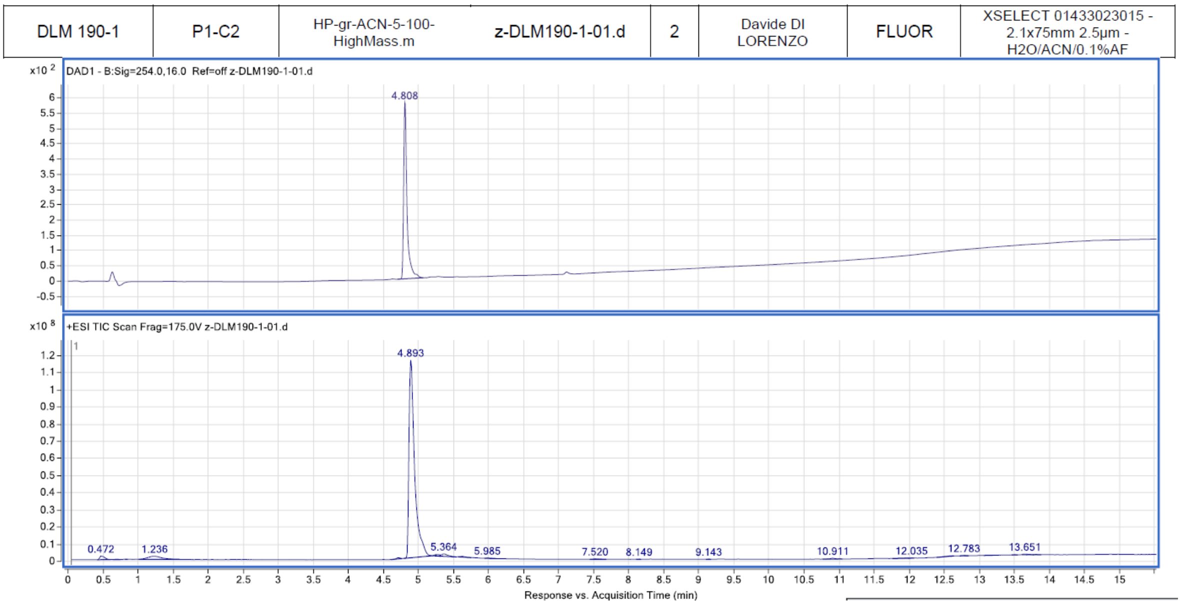

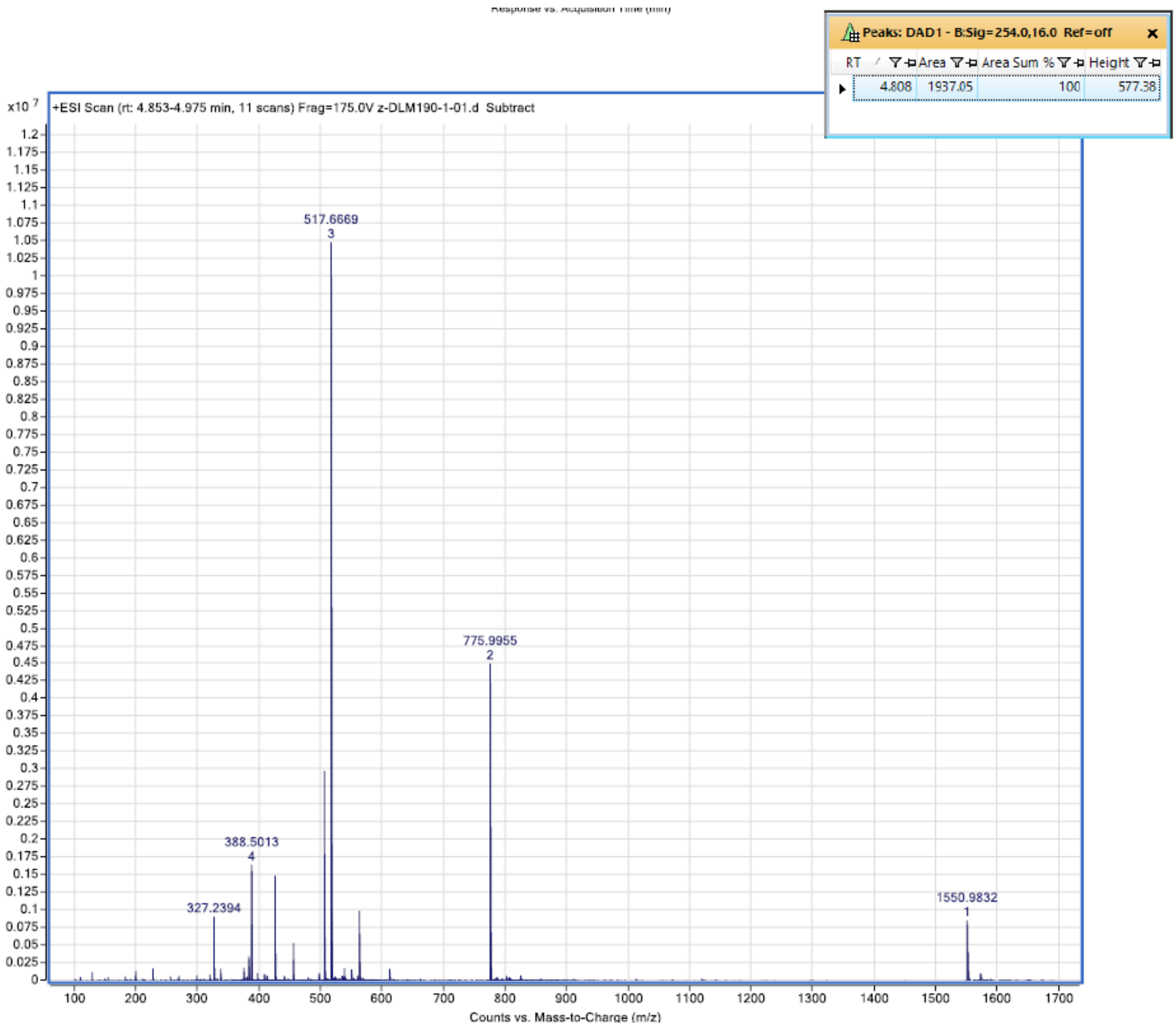

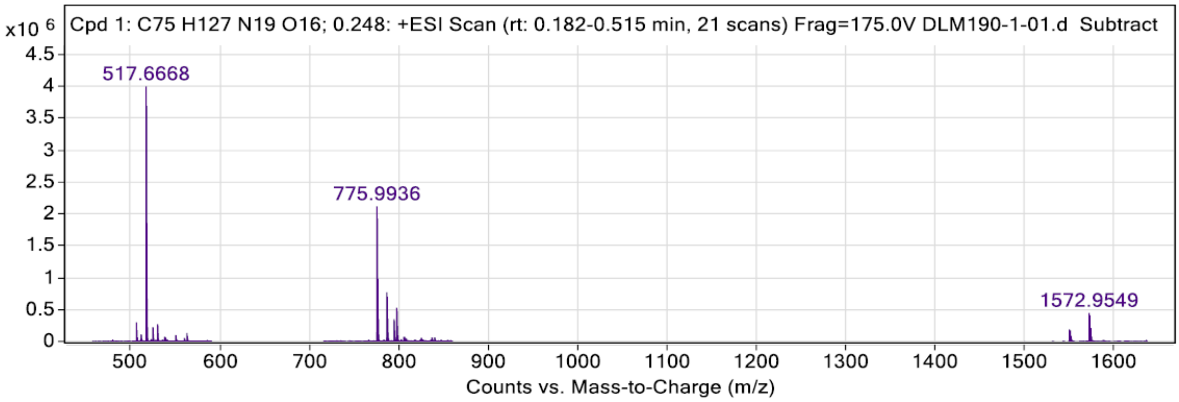

**Peak List**

| *m/z* | z | Abundance | Target Formula | Target Ion Species | Calculated m/z | Diff (mDa) | Diff (ppm) |
|---|---|---|---|---|---|---|---|
| 517,6668 | 3 | 3998935,25 | C75H127N19O16 | (M+3H)+3 | 517,6642 | 2,55 | **4,93** |
| 518,0010 | 3 | 3693351,50 | C75H127N19O16 | (M+3H)+3 | 517,9985 | 2,52 | **4,86** |
| 518,3351 | 3 | 1854781,75 | C75H127N19O16 | (M+3H)+3 | 518,3328 | 2,27 | **4,38** |
| 775,9936 | 2 | 2116123,00 | C75H127N19O16 | (M+2H)+2 | 775,9927 | 0,91 | **1,17** |
| 776,4950 | 2 | 1924265,13 | C75H127N19O16 | (M+2H)+2 | 776,4941 | 0,89 | **1,14** |
| 1550,9712 | 1 | 256764,48 | C75H127N19O16 | (M+H)+ | 1550,9781 | -6,92 | **-4,46** |
| 1551,9743 | 1 | 233447,08 | C75H127N19O16 | (M+H)+ | 1551,9810 | -6,69 | **-4,31** |
| 1572,9530 | 1 | 611022,88 | C75H127N19O16 | (M+Na)+ | 1572,9600 | -7,08 | **-4,50** |
| 1573,9560 | 1 | 561111,56 | C75H127N19O16 | (M+Na)+ | 1573,9630 | -7,00 | **-4,45** |

| Residue | δ NH (ppm) | δ Hα (ppm) $^3J$ (Hz) $C_\alpha$ | δ Hβ (ppm) $C_\beta$ | δ and other protons (ppm) and C $_\delta$ | $^{13}C$ δ |
|---|---|---|---|---|---|
| **Lys-1** | $NH_2$ (29) : ---<br>$NH_2$ (132) : 7.51 | $CH_\alpha$ (95): 4.00<br>54.70 | $CH_{2\beta}$ (98) : 1.82 | $CH_{2\gamma}$ (116) : 1.33<br>$CH_{2\delta}$ (130) : 1.62<br>$CH_{2\varepsilon}$ (131) : 2.92 | CO (86) : -- |
| **Val-2** | NH (85) : 8.56 | $CH_\alpha$ (84) : 4.09 | $CH_\beta$ (94) : 1.92 | $CH_{3\gamma}$ (112) : 0.83<br>$CH_{3\gamma}$ (129) : 0.88 | CO (83) : -- |
| **Val-3** | NH (82) : 8.40 | $CH_\alpha$ (82) : 4.05 | $CH_\beta$ (92) : 1.92 | $CH_{3\gamma}$ (109) : 0.80<br>$CH_{3\gamma}$ (110) : 0.86 | CO (80) : -- |
| **Val-4** | NH (79) : 8.31 | $CH_\alpha$ (78) : 3.98 | $CH_\beta$ (90) : 1.88 | $CH_{3\gamma}$ (106) : 0.75<br>$CH_{3\gamma}$ (107) : 0.82 | CO (77) : -- |
| **Ile-5** | NH (76) : 8.15 | $CH_\alpha$ (75) : 4.10 | $CH_\beta$ (125) : 1.58 | $CH_{3\gamma}$ (127) : 0.68<br>$CH_{2\gamma}$ (126) : 1.30<br>$CH_{3\delta}$ (128) : 1.01 | CO (28) : -- |
| **Thr-6** | NH (72) : 8.17 | $CH_\alpha$ (70): 4.17 | $CH_\beta$ (73) : 4.01 | $CH_{3\gamma}$ (99) : 1.06 | CO (69) : -- |
| **Isox-7** | NH (68) : 8.43 | -- | $CH_{2\beta}$ (13) : 3.68 / 3.88<br>$CH_{2\beta}$ (9) : 3.353/ 3.73 | 5 $CH^{Ar}$ (1-6) : 7.68 – 7.70 (m, 2H) / 7.49 – 7.41 (m, 3H) / 133.4, 130.9, 128.9, | CO (12) : --<br>C (7) : --<br>C (10) : -- |
| **Thr-8** | NH (14) : 8.18 | $CH_\alpha$ (16): 4.25 | $CH_\beta$ (17) : 4.15 | $CH_{3\gamma}$ (46) : 1.06 | CO (18) : -- |
| **Lys-9** | NH (19) : 8.50<br>$NH_2$ (118) : 7.49 | $CH_\alpha$ (21): 4.28 | $CH_{2\beta}$ (22) : 1.73 | $CH_{2\gamma}$ (57) : 1.38<br>$CH_{2\delta}$ (65) : 1.62<br>$CH_{2\varepsilon}$ (66) : 1.92 | CO (23) : -- |
| **Val-10** | NH (24) : 8.18 | $CH_\alpha$ (26) : 3.98 | $CH_\beta$ (27) : 1.92 | $CH_{3\gamma}$ (55) : 0.80<br>$CH_{3\gamma}$ (119) : 0.87 | CO (28) : -- |
| **Ile-11** | NH (29) : 8.25 | $CH_\alpha$ (31) : 4.04 | $CH_\beta$ (40) : 1.74 | $CH_{3\gamma}$ (52) : 1.08<br>$CH_{2\gamma}$ (53) : 1.39<br>$CH_{3\delta}$ (120) : 0.77 | CO (32) : -- |
| **Leu-12** | NH (33) : 8.34 | $CH_\alpha$ (34) : 4.26 | $CH_{2\beta}$ (42) : 1.40 / 1.46 | $CH_{2\gamma}$ (58) : 1.53 / 0.86<br>$CH_{3\delta}$ (121) : 0.79<br>$CH_{3\delta}$ (63) : 0.79 | CO (35) : -- |
| **His-13** | NH (36) : 8.47<br>$CONH_2$ (39) : ND | $CH_\alpha$ (37): 4.59 | $CH_{2\beta}$ (44) : 3.10 / 3.22 | $CH^{Ar}$(123) : 7.24<br>$CH^{Ar}$(122) : 8.55 | CO (38) : -- |

***Compound 1S (a,b):*** *Methyl N-(5-(azidomethyl)-3-phenyl-4,5-dihydroisoxazole-5-carbonyl)-O-(tert-butyl)-L-allothreoninate*

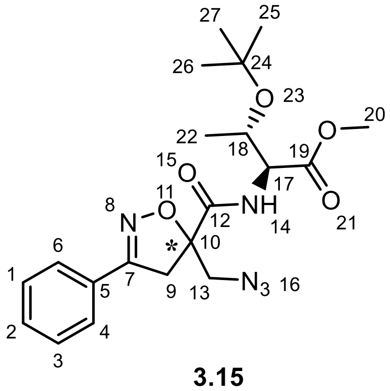

**3.15**

**1S** was obtained as a pale yellow oil following the general procedure **C** starting from **9a** (963 mg, 3.91 mmol, 1 eq.). The crude product was purified by chromatography on silica gel eluting with Hexane/AcOEt to yield **1S** (1.51 g, 3.64 mmol, 93%) as a mixture of inseparable diastereoisomers.

**Molecular weight**: 417.20 g/mol

**R*f*** (Hexane/AcOEt = 85:15): 0.3

**MS**: Calcd. for $[C_{20}H_{27}N_5O_5 + H]^+$: M/z 418.2085 found: 418.31 $[M + H]^+$

**(1S a) $^1$H NMR (300 MHz, $CDCl_3$):** δ 7.75 – 7.65 (2H, m, H6, H4), 7.62 (1H, d, *J* = 9.1 Hz, H14), 7.51 – 7.38 (3H, m, H1, H2, H3), 4.44 (1H, dd, *J* = 9.1, 2.1 Hz, H17), 4.26 (1H, dd, *J* = 6.3, 2.3 Hz, 1H, H18), 3.91 (1H, d, *J* = 13.3 Hz, H13'), 3.73 – 3.64 (2H, m, H13'', H9') 3.63 (3H, s, H20), 3.51 (1H, d, *J* = 17.52 Hz, H9''), 1.22 (3H, d, *J* = 6.24 Hz, H22), 1.12 (9H, s, H25, H26, H27).

**(1S a) $^{13}$C NMR (101 MHz, $CDCl_3$)**: δ 171.09 (C19-A), 170.49 (C12-A), 157.35 (C7-A), 130.82 (C2-A), 128.86 (C1-A, C3-A), 128.37 (C5-A), 126.96 (C6-A, C4-A), 89.18 (C10-A), 74.24 (C24-A), 67.09 (C18-A), 58.14 (C18-A), 54.91 (C13-A), 52.23 (C20-A), 42.51 (C9-A), 29.70 (C25-A, C26-A, C27-A), 20.95 (C22-A).

**(1S b) $^1$H NMR (400 MHz, $CDCl_3$):** δ 7.75 – 7.65 (2H, m, H6, H4), 7.62 (1H, d, *J* = 9.1 Hz, H14), 7.51 – 7.38 (3H, m, H1, H2, H3), 4.41 (1H, dd, *J* = 9.0, 2.3 Hz, H17), 4.26 (1H, dd, *J* = 6.3, 2.3 Hz, H18), 3.91 (1H, d, *J* = 13.3 Hz, H13'), 3.76 (3H, s, H20), 3.72 (1H, d, *J* = 18.0 Hz, H9') , 3.66 – 3.60 (2H, m, H9'', H13''), 1.14 (9H, s, H26, H27, H28), 1.11 (d, *J* = 6.2 Hz, 3H, H22).

**(1S b) $^{13}$C NMR (101 MHz, $CDCl_3$)**: δ 171.43 (C19-B), 170.46 (C12-B), 157.55 (C7-B), 130.84 (C2-B), 128.88 (C1-B, C3-B), 128.42 (C5-B), 126.99 (C6-B, C4-B), 89.46 (C10-B), 74.29 (C24-B), 66.94 (C18-B), 58.30 (C17-B), 54.39 (C13-B), 52.35 (C20-B), 41.63 (C9-B), 28.35 (C25-B, C26-B, C27-B), 20.98 (C22-B).

***Compound 2S a** : Methyl N²-(N-((S)-5-(azidomethyl)-3-phenyl-4,5-dihydroisoxazole-5-carbonyl)-O-(tert-butyl)-L-allothreonyl)-N⁶-(tert-butoxycarbonyl)-L-lysinate*

***Compound 2S b** : Methyl N²-(N-((R)-5-(azidomethyl)-3-phenyl-4,5-dihydroisoxazole-5-carbonyl)-O-(tert-butyl)-L-allothreonyl)-N⁶-(tert-butoxycarbonyl)-L-lysinate*

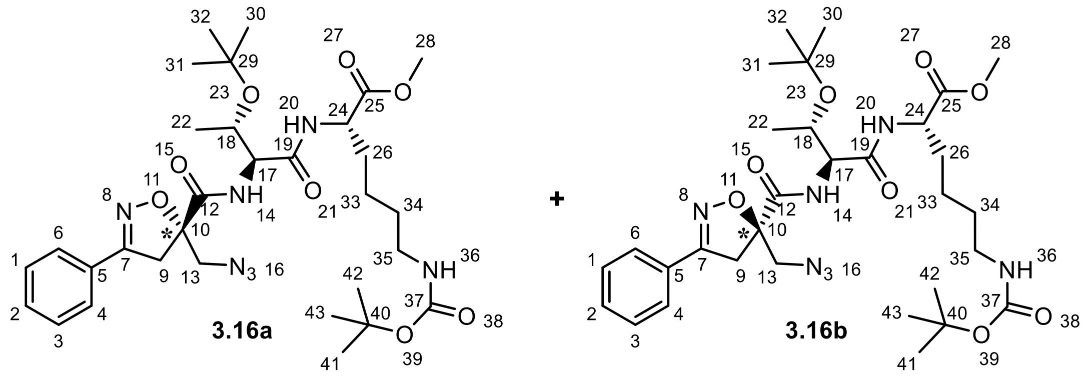

**3.16a** + **3.16b**

After saponification of **1S** according to procedure **B**, in a round bottom flask equipped with a magnetic stirrer, deprotected **1S** (927 mg, 3.77 mmol, 1 eq.) was suspended in dry DCM (20 mL), then the solution was cooled to 0 °C. At this moment, H-Thr(OtBu)-OMe (947 mg, 5.65 mmol, 1.5 eq) and $T_3P$ solution 50% in EtOAc (4.10 mL, 8.88 mmol, 3 eq.) were added and secondly DIPEA was added until pH = 8 (22.6 mmol, 5 eq.). After stirring overnight at room temperature, the reaction mixture was successively washed with a 5% aqueous solution of KHSO4 (20 mL), an aqueous solution of NaHCO3 (20 mL), and brine (25 mL). Then, the organic layer was isolated, dried over $Na_2SO_4$, filtered, and concentrated under reduced pressure. The crude pale-yellow oil obtained was purified by chromatography on silica gel eluting with DCM/$Et_2O$ 98/2 to afford compounds 2Sa (316 mg, 0.5 mmol, 13%) and 2Sb (341 mg, 0.54 mmol, 14%) separately as white solid crystals.

**Molecular weight**: 645.35 g/mol

**(2S a) R*f*** (DCM/$Et_2O$ 98/2) = 0.3

**(2S b) R*f*** (DCM/$Et_2O$ 98/2) = 0.18

**MS**: Calcd. for $[C_{31}H_{47}N_7NaO_8 + Na]^+$: m/z 668.3378 found: 668.37

**(2S a) $^1$H NMR (400 MHz, $CDCl_3$)**: δ 7.97 (1H, d, *J* = 5.9 Hz, NH14), 7.66 (3H, m, H6, H4, NH20), 7.49–7.35 (3H, m, H1, H2, H3), 4.60–4.43 (2H, m, H24, NH36), 4.35 (1H, dd, *J* = 5.9 Hz and 4.0 Hz, H17), 4.26 (1H, dd, *J* = 6.4 Hz and 4.0 Hz, H18), 3.84 (1H, d, *J* = 13.0 Hz, H13’), 3.76–3.68 (4H, m, H9’, H28), 3.64 (1H, d, *J* = 13.0 Hz, H13’’), 3.45 (1H, d, *J* = 17.5 Hz, H9’’), 3.08 (2H, m, *J* = 6.4 Hz, H35), 1.90–1.77 (1H, m, H26’), 1.7–1.64 (1H, m, H26’’), 1.43 (11H, s, H34, H41, H42, H43), 1.33 (11H, bs, H33, H30, H31, H32), 1.16 (3H, d, *J* = 6.4 Hz, H22).

**(2S a) $^{13}$C NMR (101 MHz, $CDCl_3$)**: δ 172.22 (C25), 170.48 (C12), 168.73 (C19), 157.04 (C7), 155.89 (C37), 130.78 (C2), 128.78 (C1, C3), 128.30 (C5), 126.99 (C4, C6), 89.04 (C10), 79.10 (C40), 75.69 (C29), 65.86 (C18), 57.44 (C17), 55.39 (C13), 52.27 (C24, C28) 42.26 (C9), 40.24 (C35), 31.79 (C26), 29.65 (C34), 28.39 (C41, C42, C43), 28.16 (C30, C31, C32), 22.45 (C33), 16.74 (C22).

**(2S b) $^{1}$H NMR (400 MHz, CDCl$_3$)**: δ 7.91 (1H, d, *J* = 6.2 Hz, NH14), 7.74–7.59 (3H, m, H6, H4, NH20), 7.52–7.38 (3H, m, H1, H2, H3), 4.56 (2H, m, H24, NH36), 4.35 (1H, dd, *J* = 6.3 Hz and 3.8 Hz, H17), 4.16 (1H, m, H18), 3.89 (1H, d, *J* = 13.1 Hz, H13'), 3.76 (3H, s, H28), 3.73–3.62 (2H, m, H13'', H9'), 3.56 (d, 1H, *J* = 17.5 Hz, H9''), 3.12 (2H, q, *J* = 6.4 H, H35), 1.96–1.82 (1H, m, H26'), 1.81–1.68 (1H, m, H26''), 1.58–1.48 (2H, m, H34), 1.45 (9H, s, H41, H42, H43), 1.29 (s, 11H, H33, H30, H31, H32), 1.02 (3H, d, *J* = 6.4 Hz, H22).

**(2S b) $^{13}$C NMR (101 MHz, CDCl$_3$)**: δ 172.27 (C25), 170.76 (C12), 168.81 (C19), 157.20 (C7), 155.93 (C37), 130.86 (C2), 128.89 (C1, C3), 128.34 (C5), 126.94 (C4, C6), 89.16 (C9), 79.14 (C40), 75.51 (C29), 65.98 (C18), 57.59 (C17), 54.65 (C13), 52.31 (C24), 52.29 (C28), 41.95 (C9), 40.27 (C35), 31.87 (C26), 29.69 (C34), 28.42 (C41, C42, C43), 28.21 (C30, C31, C32), 22.54 (C33), 17.26 (C22).

***Compound 3S b:*** *N$^2$-(N-((R)5-(azidomethyl)-3-phenyl-4,5-dihydroisoxazole-5-carbonyl)-O-(tert-butyl)-L-allothreonyl)-N$^6$-(tert-butoxycarbonyl)-L-lysine*

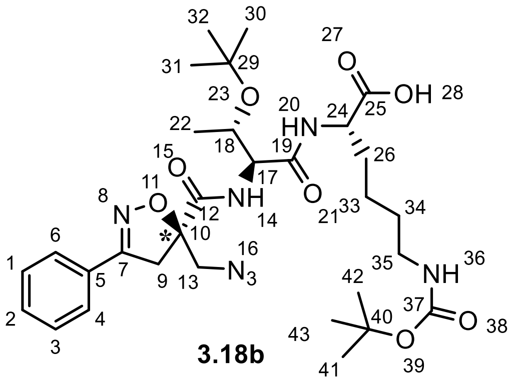

**3.18b**

**3S b** (341 mg, 0.54 mmol, quant.) was obtained following the general procedure **B** starting from **2S b** (341 mg, 0.54 mmol, 1 eq.) as a white solid in quantitative yield.

**Molecular weight**: 631.34 g/mol

***Rf*** (Hex/Et  7/3) = 0

**MS**: Calcd. for [$C_{30}H_{46}N_7O_8$ + H]$^+$: m/z 632.3402, found: 632.01; Calcd. for [$C_{30}H_{46}N_7O_8$ + Na]$^+$: m/z 654.3222 found: 654.38

**$^{1}$H NMR (400 MHz, CDCl$_3$)**: δ 7.92 (1H, d, *J* = 6.5 Hz, NH14), 7.74–7.60 (3H, m, H6, H4, NH20), 7.45 (3H, m, H1, H2, H3), 6.00 (1H, bs, NH36), 4.74–4.46 (1H, m, H24), 4.37 (1H, *J* = 6.5Hz and 3.7 Hz, 1H, H17), 4.18 (1H, m, H18), 3.92 (1H, d, *J* = 13.1 Hz, H13'), 3.77–3.64 (2H, m, H13'', H9'), 3.56 (1H, d, *J* = 17.5 Hz, H9''), 3.14 (2H, m, H35), 2.11–1.71 (2H, m, H26), 1.46 (13H, bs, H41, H42, H43, H34, H33), 1.28 (9H, s, H30, H31, H32), 1.03 (3H, d, *J* = 6.3 Hz, H22).

**$^{13}$C NMR (101 MHz, CDCl$_3$)**: δ 174.51 (C25), 170.93 (C12), 169.26 (C17), 157.30 (C7), 130.91 (C2), 128.91 (C1, C3), 128.29 (C5), 126.97 (C4, C6), 89.15 (C10), 79.55 (C40), 75.47 (C29), 66.05 (C18), 57.81 (C17), 54.72 (C13), 52.44 (C24), 41.95 (C9), 40.29 (C35), 31.50 (C26), 29.70 (C34), 28.43 (C41, C42, C43), 28.24 (C30, C31, C32), 22.57 (C33), 17.66 (C22).

***Compound 4S b:*** *N*[2]*-(N-((R)5-(aminomethyl)-3-phenyl-4,5-dihydroisoxazole-5-carbonyl)-O-(tert-butyl)-L-allothreonyl)-N*[6]*-(tert-butoxycarbonyl)-L-lysine*

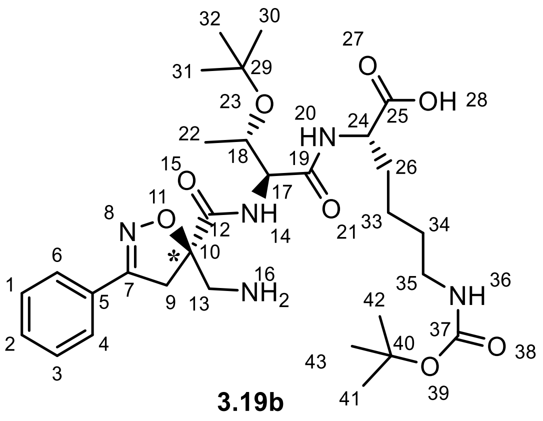

**4S b** was obtained as a pale yellow oil following the general procedure **D** starting from **3S b** (193 mg, 0.56 mmol, 1 eq.), $H_2O$ (49 eq.) and $PMe_3$ (7 eq.). The crude obtained was directly used in the next step. The compound was used in the next step without any further purification. A small amount of crude was purified to fulfill the NMR characterization.

**Molecular weight**: 605.35 g/mol

**R*f*** (DCM/MeOH 9/1) = 0.1

**MS**: Calcd. for $[C_{30}H_{47}N_5O_8 + Na]^+$: m/z 606.3497; found: 606.52 $[M + Na]^+$

**$^{1}$H NMR (400 MHz, $CD_3OD$)**: δ 7.85–7.63 (2H, m, H6, H4), 7.58–7.34 (3H, m, H1, H2, H3), 4.35–4.21 (3H, m, H17, H18, H24), 3.81 (1H, d, *J* = 17.7 Hz, H9'), 3.67 (1H, d, *J* = 17.8 Hz, H9''), 3.55 (1H, d, *J* = 13.5 Hz, H13'), 3.33 (1H, m, H13''), 3.04 (2H, m, H35), 1.92–1.67 (2H, m, H26), 1.54–1.35 (13H, m, H41, H42, H43, H33, H34), 1.21 (9H, s, H30, H31, H32), 1.13 (3H, d, *J* = 6.3 Hz, H22).

**$^{13}$C NMR (101 MHz, $CD_3OD$)**: δ 177.19 (C25), 171.98 (C12), 169.78 (C19), 158.15 (C7), 130.70 (C2), 128.61 (C1, C3), 128.21 (C5), 126.70 (C4, C6), 88.09 (C10), 80.78 (C40), 74.41 (C29), 66.69 (C18), 59.37 (C17), 54.75 (C24), 44.83 (C13), 42.54 (C9), 39.97 (C35), 32.22 (C26), 29.36 (C34), 27.45 (C41, C42, C43), 27.42 (C30, C31, C32), 22.59 (C33), 19.53 (C22).

***Compound 5S b:*** *$N^2$-(N-((R)-5-(((((9H-fluoren-9-yl)methoxy)carbonyl)amino)methyl)-3-phenyl-4,5-dihydroisoxazole-5-carbonyl)-O-(tert-butyl)-L-allothreonyl)-$N^6$-(tert-butoxycarbonyl)-L-lysine*

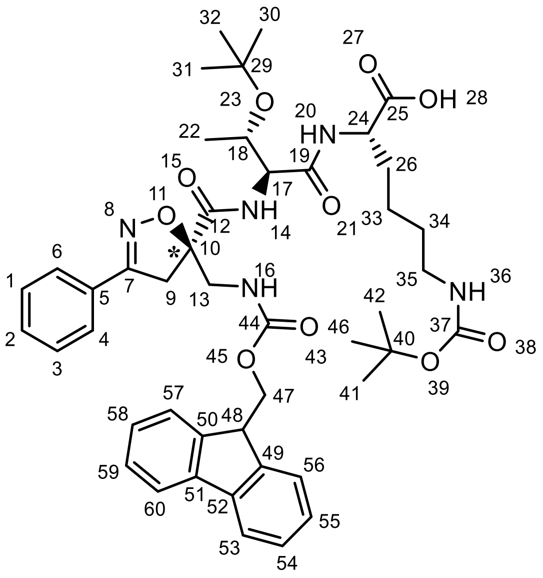

**3.20b**

**5S b** was obtained following the general procedure **E** from **4S b** (0.55 mmol). TLC (DCM/MeOH + AcOH 95/5 + 1%) was used to monitor the reaction. The crude obtained was purified by chromatography on silica gel using DCM/MeOH + AcOH 97/3 + 1% as eluent to yield **5S b** (305 mg, 0.3685 mmol) as a white spongy solid in 67% yield..

**Molecular weight**: 827.42 g/mol

**R*f*** (DCM/MeOH + AcOH 97/3 + 1%) = 0.5

**$[\alpha]_{20}^{D}$ (3.20b)**: +25.1° ([c] = 0.01 in $CH_2Cl_2$)

**MS**: Calcd. for $[C_{45}H_{57}N_5O_{10} + Na]^+$: m/z 850.3998 found: 850.66 $[M + Na]^+$

**$^1$H NMR (400 MHz, $CD_3OD$)**: δ 7.76 (2H, dd, *J* = 7.7 Hz and 3.6 Hz, H53, H60), 7.73–7.66 (2H, m, H4, H6), 7.62 (2H, d, *J* = 7.5 Hz, H54, H59), 7.45 (3H, m, H1, H2, H3), 7.34 (2H, t, *J* = 7.5 Hz, H56, H57), 7.29–7.14 (m, 2H, H55, H58), 4.44 (1H, m, 1H, H24), 4.37 – 4.25 (3H, m, H47, H17), 4.23 – 4.10 (2H, m, H18, H48), 3.84 (1H, d, *J* = 15.3 Hz, H13'), 3.77–3.63 (2H, m, H13'', H9'), 3.57 (1H, d, *J* = 17.7 Hz, H9''), 3.02 (2H, dd, *J* = 6.7 Hz, H35), 1.97–1.81 (1H, m, H26'), 1.81–1.69 (1H, m, H26''), 1.43 (bs, 12H, H41, H42, H43, H34, H33), 1.31 (1H, m, H34), 1.21 (9H, s, H30, H31, H32), 1.11 (3H, d, *J* = 6.2 Hz, H22).

**$^{13}$C NMR (101 MHz, $CD_3OD$)** δ 172.27 (C25), 170.11 (C19), 157.89 (C7), 157.80 (C37), 157.09 (C44), 143.88 (C49, C50), 141.12 (C51, C52), 130.48 (C2), 128.56 (C1, C3), 128.55 (C5), 127.35 (C56, C57), 126.77 (C55, C58), 126.63 (C4, C6), 124.90 (C54, C59), 119.46 (C53, C60), 89.47 (C10), 79.60 (C40), 74.57 (C29), 66.91 (C18), 66.74 (C47), 58.35 (C17), 51.55 (C24), 44.70 (C13), 41.09 (C9), 39.77 (C35), 31.34 (C26), 29.17 (C34), 27.41 (C41, C42, C43), 27.38 (C30, C31, C32), 22.56 (C33), 18.53 (C22).

### Single crystal X-ray diffraction analysis of intermediate 11b

CCDC 2363965 contains the supplementary crystallographic data for this paper. The data is available from the Cambridge Crystallographic Data Centre via www.ccdc.cam.ac.uk/structures. An X-ray quality colourless needle (Figure S1) was obtained from the crystallization of the freshly synthesized compound from diethyl ether:hexane 1:1, overnight, with the slow evaporation method. The specimen is weakly pleochroic when seen through polarized light.

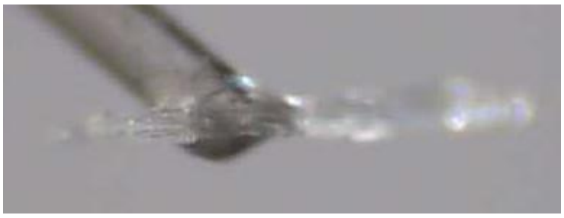

**Figure S1**. A single crystal of **11b** is mounted on the top of a capillary fibre with a drop of bicomponent epoxy glue. The dimensions of the crystal are ~ 0.925 x 0.075 x 0.075 mm.

The data was collected at room temperature with a Bruker AXS Smart APEX 3-circle diffractometer equipped with a normal focus sealed X-ray Mo tube ($\lambda$=0.71073 Å) operating at a nominal power of 50 kV · 30 mA. A graphite monochromator was used in conjunction with a CCD detector. A 99.4 % complete sphere of diffraction data was recorded up to $\sin\theta/\lambda$ = 0.56 $Å^{-1}$, resulting in 4871 independent reflections, of which 3151 were significantly above the background ($I > 2\sigma(I)$). Integration of the diffraction effects was carried out with Bruker SAINT+. The raw data were corrected for absorption and possible anisotropies of the primary beam with SADABS [Bruker (2001). SADABS. Bruker AXS Inc., Madison, Wisconsin, USA]. The structure was solved by direct methods and subsequently refined with the SHELX suite of programs.[2] The least squares algorithm consisted of 451 parameters and 3 geometrical restraints. It converged to R1(F) = 0.1561 for 3151 $F_o > 4\sigma(F_o)$, with maximum and minimum Fourier residuals as large as $\Delta\rho$ = +1.45 / –0.33 e/$Å^{-3}$ (see below).

The compound crystallizes in the triclinic chiral P1 space group with 2 formula units in the asymmetric unit, corresponding to the whole cell (Z = Z' = 2). The unit cell parameters (Å, deg) are *a* = 5.9736(12), *b* = 10.901(2), *c* = 13.209(3), $\alpha$ = 93.05(3), $\beta$ = 93.22(3), $\gamma$ = 90.24(3), as estimated from 979 intense reflections among 6.2 and 58.8 deg in 2$\theta$ (final integration result). The cell volume reads V = 857.6(3) $Å^3$, corresponding to estimated density $\rho$ = 1.334 g·$cm^{-3}$.

Figure S2 shows the asymmetric unit of the title compound, while Figure S3 summarizes the main features of crystal packing. The compound is chiral; both molecules in the asymmetric unit share the same absolute configuration, that is, R, S at the stereogenic centres C9 (quaternary carbon), C12 (tertiary carbon). See Figure S2 for the atom numbering. The correctness of these attribution is secured by the value of the Flack parameter [3] computed on the final model, -0.8(9).

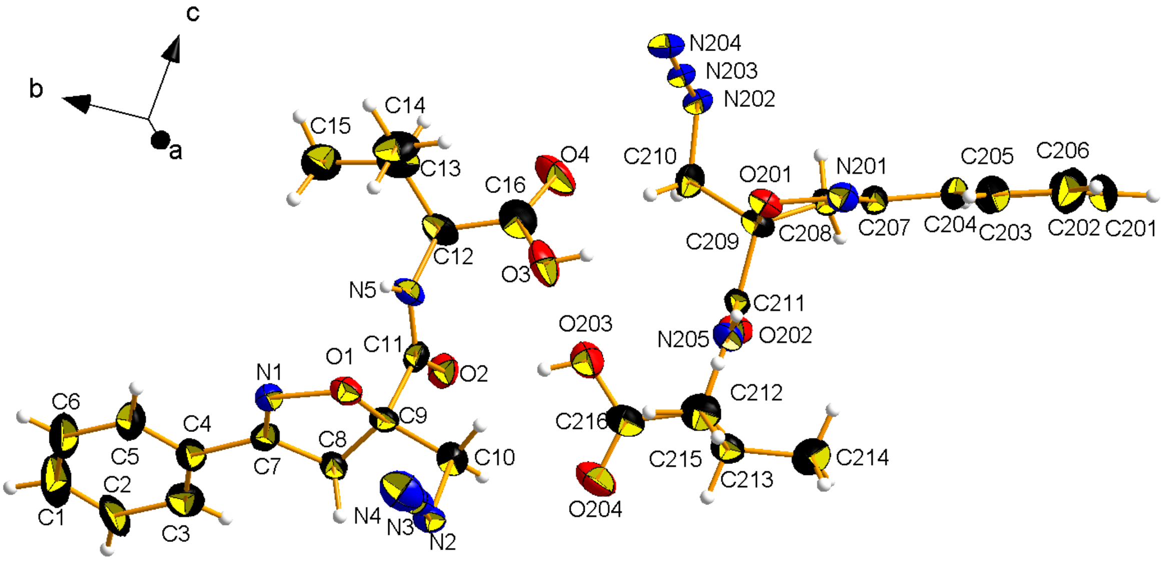

**Figure S2**. Asymmetric unit of **10b**, with the atom-numbering scheme highlighted for the symmetry–independent part of the molecule. Thermal ellipsoids at RT were drawn at the 30 % probability level. Atoms are represented with the usual colour code (C: black; N: blue; O: red; H: white).

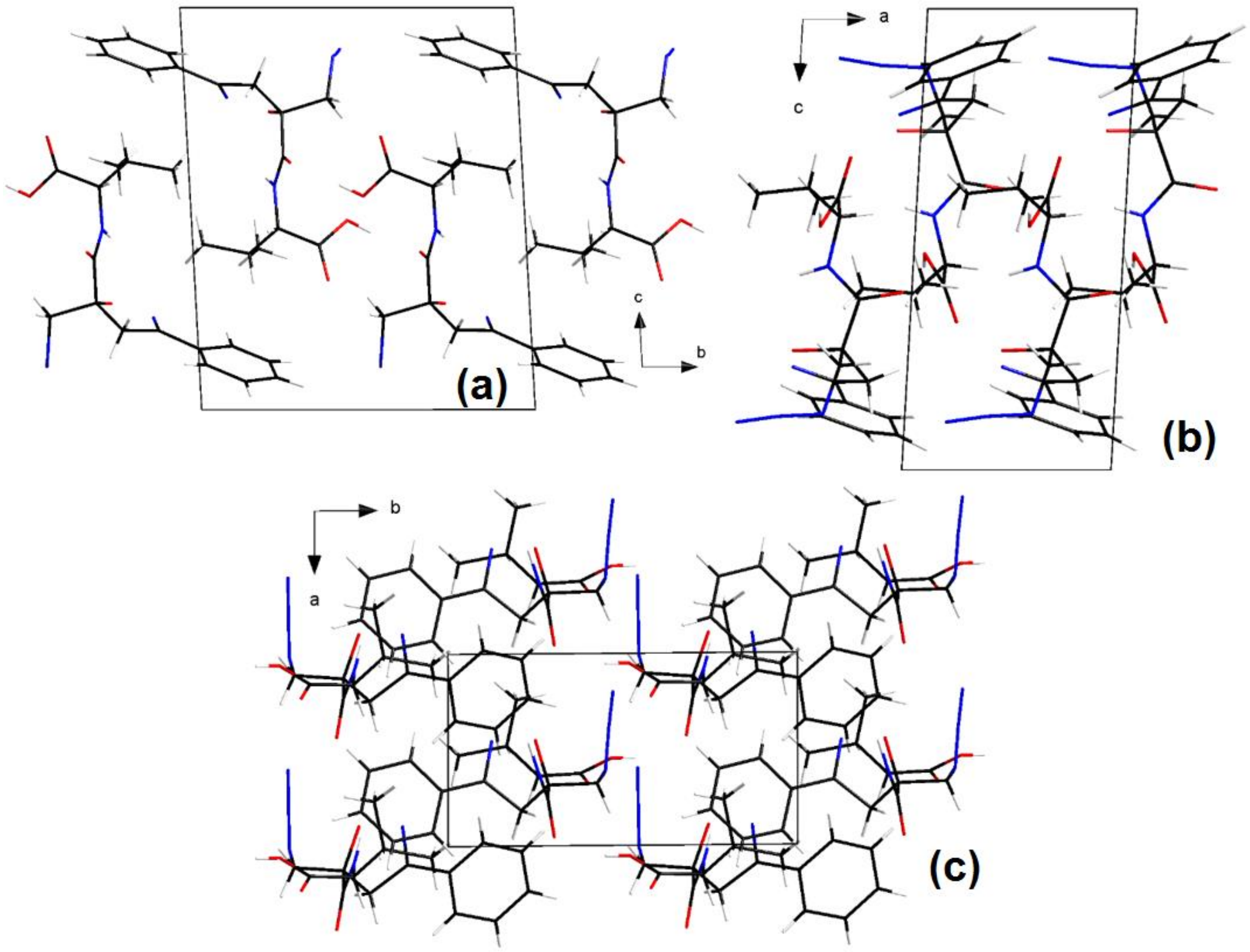

**Figure S3.** Wires–stick representation of the crystal packing of **10b** at RT, as seen along the *a* (a), *b* (b) and *c* (c) cell axes. Colour code as in Figure S2.

The 5-membred isoxazole–like rings share the same distorted envelope conformation, with puckering amplitudes and phases [4] 0.2116 Å, 135.19 (molecule 1, from C1 forward), or 0.2204 Å, 134.85 deg (molecule 2, from C201 forward). The C9 and C209 atoms lie 0.32(1)-0.33(1) Å apart from the mean least–squares plane computed from the atomic coordinates of the remaining atoms in the cycle (O1, N1, C7 and C8 or O201, N201, C207 and C208; see Figure S2 for the atom numbering).

Figure S3 shows the main packing motifs in the (b,c), (a,c) and (a,b) planes. The title compound has at least one strong hydrogen bond donor (the acidic function), which in fact is employed to form zig-zag chains that run parallel to the a cell axis (Figure S4, Table S1). In turn, these involve alternately both the molecules in the asymmetric unit. Neither relevant intramolecular hydrogen bonds nor intermolecular stacking interactions were detected. The azide groups are aligned roughly along a, while the aromatic rings form apolar layers sandwiched between continuous $-N_3$ and –COOH groups (Figure S4).

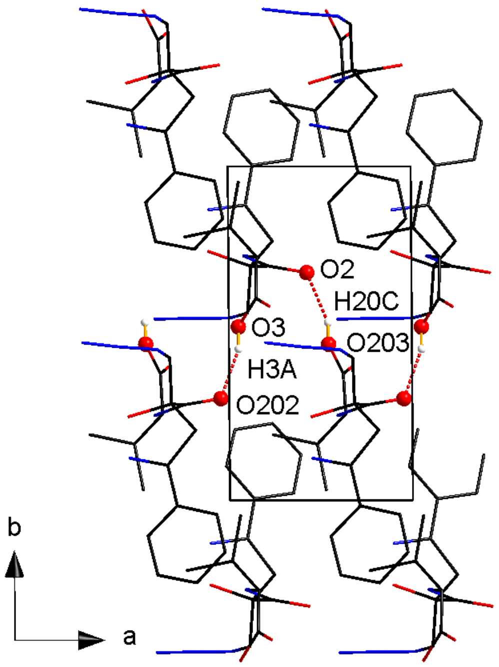

**Figure S4.** O–H···O hydrogen bond motif along *a*. Same colour code as in Figures S2-S3. Nonessential hydrogen atoms were omitted for clarity.

**Table S1**. Symmetry-independent intermolecular hydrogen bonds in DLM068B. Estimated standard deviations are reported in parentheses.

| O–H···O | $d_{O–H}$ / Å | $d_{H\cdots O}$ / Å | $d_{O\cdots O}$ / Å | $\alpha_{OHO}$ / deg | Symmetry |
|---|---|---|---|---|---|
| O3–H3A···O202 | 0.82 | 1.92 | 2.715(1) | 162 | -1+x, y, z |
| O203–H20C···O2 | 0.82 | 1.92 | 2.700(1) | 160 | x, y, z |

As a general remark, the specimen employed for the present analysis had a low scattering power and was slightly twinned. The minor twin component (<1 % *w/w*) affected only low-angle reflections and was ignored in the structure refinement procedure. The title compound crystallizes as very thin and long needles, which tend to exfoliate into smaller pieces even under moderate mechanical stress. Despite extensive crystallization attempts, no better specimens were found. However, the resolution of the present dataset is high enough to unequivocally solve the structure, which was the main aim of the X-ray experiment. The current structural model does not account for a couple of weak positive residual Fourier peaks, roughly 1.3-1.4 e·Å$^{-3}$ large, that lie close to the azide groups of both the molecules in the asymmetric unit. More in detail, these residues are placed at roughly 1.6-1.8 Å from nitrogens N2 and N202 (Figure S2), along the main azide axis. They could be related to co-crystallized water molecules, but we prefer to not explicitly attribute them as higher-quality specimens should be required to gain insights on their nature.

### Circular dichroism of $N_3$-Isox-$\beta^{2,2}$-OH and of compounds 1 and 2 at 24 hrs

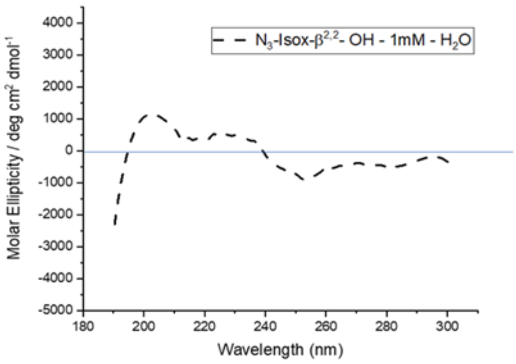

**Figure S5.** CD spectrum of $N_3$-Isox-$\beta^{2,2}$-OH (1 mM), showing no interference for the CD interpretation of secondary structures of peptides.

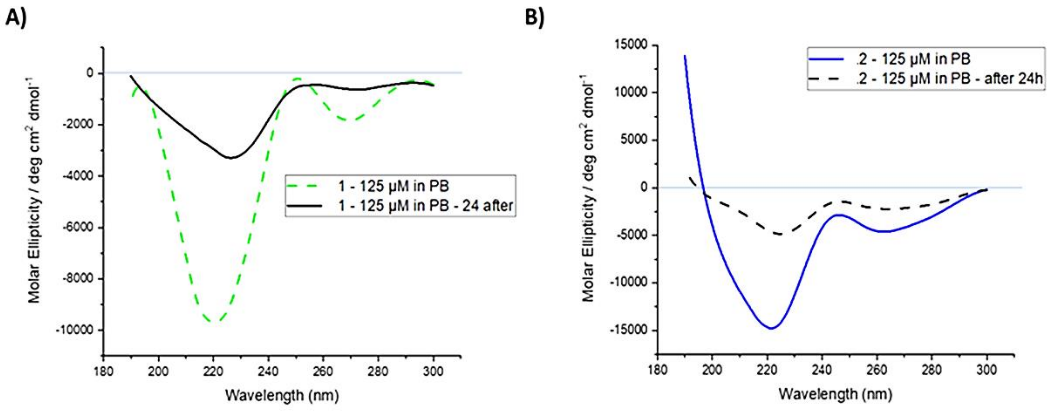

**Figure S6.** Comparison of CD spectra of **1** and **2** at time 0 and at 24h in 20 mM PB pH 7.2 **A**) CD spectrum of **1** at 125 µM; **B**) CD spectrum of **2** at 125 µM.

### Circular dichroism of compounds 1 and 2 in PB at pH 5.1

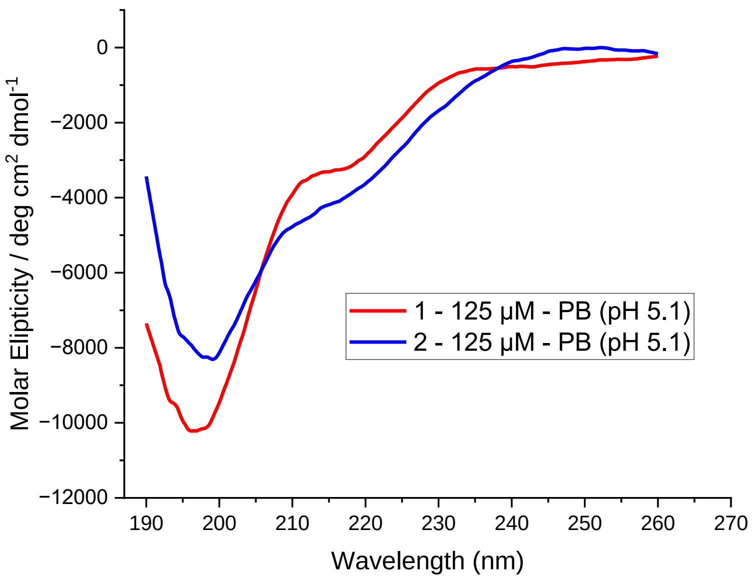

**Figure S7.** CD spectra of compounds **1** and **2** in 20 mM PB (125 µM) at pH 5.1; the calculated ratio $[\Phi]_{218\ nm}$ / $[\Phi]_{195\ nm}$, indicating the propensity to fold in a hairpin like structure (value < 1 random coil majority, value = 1 random coil and hairpin in 1:1 ratio, value > 1 hairpin preference), was 0.32 for **1** and 0.52 for **2**.

### NMR conformational analysis of compounds 1 and 2

| Residue | δ NH (ppm) | δ Hα (ppm)<br>$^3J$ (Hz)<br>$C_{\alpha}$ | δ Hβ (ppm)<br>$C_{\beta}$ | δ and other protons (ppm)<br>and $C_{\delta}$ | $^{13}C$ δ |
|---|---|---|---|---|---|
| **Val-1** | $NH_2$ (97) : --- | $CH_{\alpha}$ (95) : 3.54<br>d, $J$ = 6.46 Hz<br>58.26 | $CH_{\beta}$ (96) : 1.93 | $CH_{3\gamma}$ (116) : 0.73<br>$CH_{3\gamma}$ (116) : 0.73 | CO (86) : -- |
| **Gln-2** | NH (85) : 8.54<br>$NH_2$ (114) : ND | $CH_{\alpha}$ (84):<br>4.16<br>52.76 | $CH_{2\beta}$ (94) :<br>1.72<br>36.0 | $CH_{2\gamma}$ (112) :<br>2.05 | CO (83) : --<br>*CO (113)* : -- |
| **Ile-3** | NH (82) : 8.09 | $CH_{\alpha}$ (81) : 3.71<br>58.03 | $CH_{2\beta}$ (92) : 1.43 | $CH_{3\gamma}$ (111) : 0.89<br>$CH_{2\gamma}$ (110) : 1.18<br>$CH_{3\delta}$ (109) : 0.55 | CO (80) : -- |
| **Ile-4** | NH (79) : 8.31 | $CH_{\alpha}$ (78) : 3.71 | $CH_{2\beta}$ (90) : 1.45 | $CH_{3\gamma}$ (107) : 0.55<br>$CH_{2\gamma}$ (106) : 0.89<br>$CH_{3d}$ (108) : 0.55 | CO (77) : -- |
| **Asn-5** | $NH_2$ (76) : 8.27<br>NH (105) : ND | $CH_{\alpha}$ (75):<br>4.35<br>49.94 | $CH_{2\beta}$ (87) :<br>2.31 / 2.35<br>36.0 | --- | CO (74) : --<br>*CO (103)* : -- |
| **Lys-6** | NH (72) : 8.09<br>$NH_2$ (102) : 7.28 | $CH_{\alpha}$ (70) : 3.92 | $CH_{2\beta}$ (73) : 1.35 | $CH_{2\gamma}$ (99) : 1.02 /<br>$CH_{2\delta}$ (100) : 1.22<br>$CH_{2\varepsilon}$ (101) : 2.48 /<br>38.87 | CO (69) : -- |
| **Isox-7** | NH (68) : 8.28 | -- | $CH_{2\beta}$ (13) : 3.40<br>/ 3.73 / 42<br><br>$CH_{2\beta}$ (9) : 3.31 /<br>3.45 / 41 | 5 $CH^{Ar}$ (1-6) : 7.45 –<br>7.38 (m, 2H) / 7.28 –<br>7.24 (m, 3H) /<br>131.0, 127.64,<br>126.89 | CO (12) : --<br>C (7) : --<br>C (10) : -- |
| **Val-8** | NH (14) : 8.02 | $CH_{\alpha}$ (16) : 3.82<br>/ 59.63 | $CH_{\beta}$ (17) : 1.89 | $CH_{3\gamma}$ (46) : 0.65 /<br>$CH_{3\gamma}$ (47) : 0.7 / | CO (18) : -- |
| **Gln-9** | NH (19) : 8.40<br>$NH_2$ (66) : ND | $CH_{\alpha}$ (21):<br>4.02 / 52.62 | $CH_{2\beta}$ (22) :<br>1.61 | $CH_{2\gamma}$ (57) :<br>1.91 | CO (23) : --<br>*CO (65)* : -- |
| **Ile-10** | NH (24) : 8.07 | $CH_{\alpha}$ (26) : 3.70<br>58.01 | $CH_{2\beta}$ (27) : 1.42 | $CH_{3\gamma}$ (55) : 0.52 /<br>$CH_{2\gamma}$ (54) : 0.80 /<br>$CH_{3\delta}$ (56) : 0.37 / | CO (28) : --- |
| **Val-11** | NH (29) : 7.93 | $CH_{\alpha}$ (31) : 3.77<br>59.1 | $CH_{\beta}$ (40) : 1.65 | $CH_{3\gamma}$ (52) : 0.58 /<br>$CH_{3\gamma}$ (53) : 0.58 / | CO (32) : -- |
| **Tyr-12** | NH (33) : 8.33 | $CH_{\alpha}$ (34):<br>4.27 / 55.06 | $CH_{2\beta}$ (42) :<br>2.65 | $CH^{Ar}$(63) : 6.85 /<br>130.4<br>$CH^{Ar}$(62) : 6.53 /<br>115.5 | CO (35) : -- |
| **Lys-13** | NH (36) : 8.16<br>$NH_2$ (51) : 7.33<br>$CONH_2$ (31) : ND | $CH_{\alpha}$ (37) : 3.93 | $CH_{2\beta}$ (44) : 1.48 | $CH_{2\gamma}$ (48) : 1.07 /<br>$CH_{2\delta}$ (49) : 1.37 /<br>$CH_{2\varepsilon}$ (50) : 2.68 /<br>39.16 | CO (38) : -- |

**Table S2.** $^1H$ and $^{13}C$ chemical shifts of 3.6 mM compound **1** in 20 mM PB/10% $D_2O$ at pH 5.1 and at 283K.

| Residue | δ NH (ppm) | δ Hα (ppm) $^3J$ (Hz) $C_\alpha$ | δ Hβ (ppm) $C_\beta$ | δ and other protons (ppm) and C $_\delta$ | $^{13}$C δ |
|---|---|---|---|---|---|
| **Val-1** | $NH_2$ (97) : --- | $CH_\alpha$ (95) : 3.56<br>d, *J* = 6.46 Hz | $CH_\beta$ (96) : 1.93 | $CH_{3\gamma}$ (116) : 0.77 /<br>$CH_{3\gamma}$ (116) : 0.77 / | CO (86) : -- |
| **Gln-2** | NH (85) : 8.55<br>$NH_2$ (114) : ND | $CH_\alpha$ (84): 4.17 | $CH_{2\beta}$ (94) : 1.72 | $CH_{2\gamma}$ (112) : 2.06 | CO (83) : --<br>*CO (113)* : -- |
| **Ile-3** | NH (82) : 8.36 | $CH_\alpha$ (81) : 3.91 | $CH_{2\beta}$ (92) : 1.53 | $CH_{3\gamma}$ (111) : 0.90 /<br>$CH_{2\gamma}$ (110) : 1.22 /<br>$CH_{3\delta}$ (109) : 0.57 / | CO (80) : -- |
| **Ile-4** | NH (79) : 8.22 | $CH_\alpha$ (78) : 3.85 | $CH_{2\beta}$ (90) : 1.51 | $CH_{3\gamma}$ (107) : 0.87 /<br>$CH_{2\gamma}$ (106) : 0.1.16 /<br>$CH_{3d}$ (108) : 0.55 / | CO (77) : -- |
| **Asn-5** | $NH_2$ (76) : 8.28<br>NH (105) : ND | $CH_\alpha$ (75): 4.33 | $CH_{2\beta}$ (87) : 2.26 | --- | CO (74) : --<br>*CO (103)* : -- |
| **Lys-6** | NH (72) : 8.15<br>$NH_2$ (102) : 7.28 | $CH_\alpha$ (70) : 3.93 | $CH_{2\beta}$ (73) : 1.48 | $CH_{2\gamma}$ (99) : 1.05 /<br>$CH_{2\delta}$ (100) : 1.27 /<br>$CH_{2\varepsilon}$ (101) : 2.55 / | CO (69) : -- |
| **Isox-7** | NH (68) : 8.28 | -- | $CH_{2\beta}$ (13) : 3.45 / 3.63<br>$CH_{2\beta}$ (9) : 3.37 / 3.53 / | 5 $CH^{Ar}$ (1-6) : 7.45 – 7.38 (m, 2H) / 7.28 – 7.24 (m, 3H) / | CO (12) : --<br>C (7) : --<br>C (10) : -- |
| **Val-8** | NH (14) : 8.02 | $CH_\alpha$ (16) : 3.86 | $CH_\beta$ (17) : 1.82 | $CH_{3\gamma}$ (46) : 0.60 /<br>$CH_{3\gamma}$ (47) : 0.60 / | CO (18) : -- |
| **Gln-9** | NH (19) : 8.43<br>$NH_2$ (66) : ND | $CH_\alpha$ (21): 4.15 | $CH_{2\beta}$ (22) : 1.72 | $CH_{2\gamma}$ (57) : 2.068 | CO (23) : --<br>*CO (65)* : -- |
| **Ile-10** | NH (24) : 8.23 | $CH_\alpha$ (26) : 3.85 | $CH_{2\beta}$ (27) : 1.52 | $CH_{3\gamma}$ (55) : 0.55 /<br>$CH_{2\gamma}$ (54) : 1.16 / 0.86 /<br>$CH_{3\delta}$ (56) : 0.53 / | CO (28) : -- |
| **Val-11** | NH (29) : 8.06 | $CH_\alpha$ (31) : 3.82 | $CH_\beta$ (40) : 1.67 | $CH_{3\gamma}$ (52) : 0.6 /<br>$CH_{3\gamma}$ (53) : 0.6 / | CO (32) : -- |
| **Tyr-12** | NH (33) : 8.39 | $CH_\alpha$ (34): 4.30 | $CH_{2\beta}$ (42) : 2.67 / 2.72 | $CH^{Ar}$(63) : 6.87 /<br>$CH^{Ar}$(62) : 6.54 / | CO (35) : -- |
| **Lys-13** | NH (36) : 8.17<br>$NH_2$ (51) : 7.31<br>$CONH_2$ (31) : ND | $CH_\alpha$ (37) : 3.93 | $CH_{2\beta}$ (44) : 1.52 | $CH_{2\gamma}$ (48) : 1.09 //<br>$CH_{2\delta}$ (49) : 1.39 /<br>$CH_{2\varepsilon}$ (50) : 2.70 /<br>39.16 | CO (38) : -- |

**Table S3.** $^1$H and $^{13}$C chemical shifts of 3.6 mM compound **2** in 20 mM PB/10% $D_2O$ at pH 5.1 and at 283K.

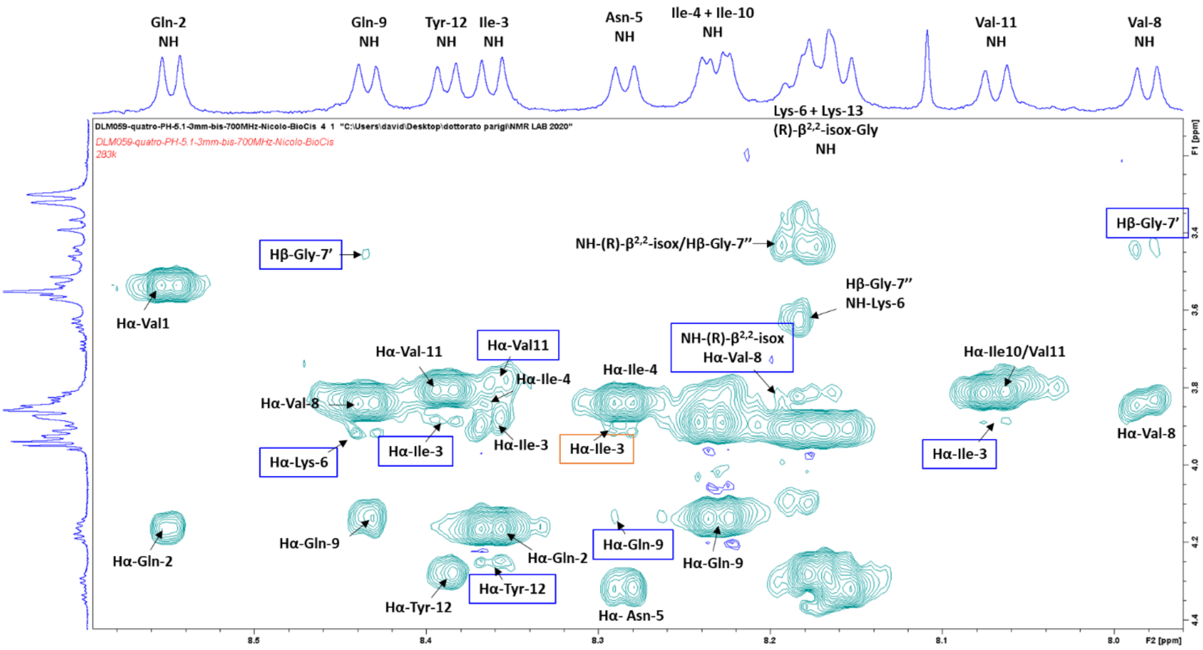

**Figure S8.** The $^{1}H$-$^{1}H$ ROESY spectrum showing sequential ROEs $CH_{\alpha}$/NH (*i*, *i+1*) of compound **2** (highlighted in black) and inter-strand $CH_{\alpha}$/NH interactions highlighted in blue.

### Transmission Electron Microscopy (TEM) images of control for compounds 1 and 2

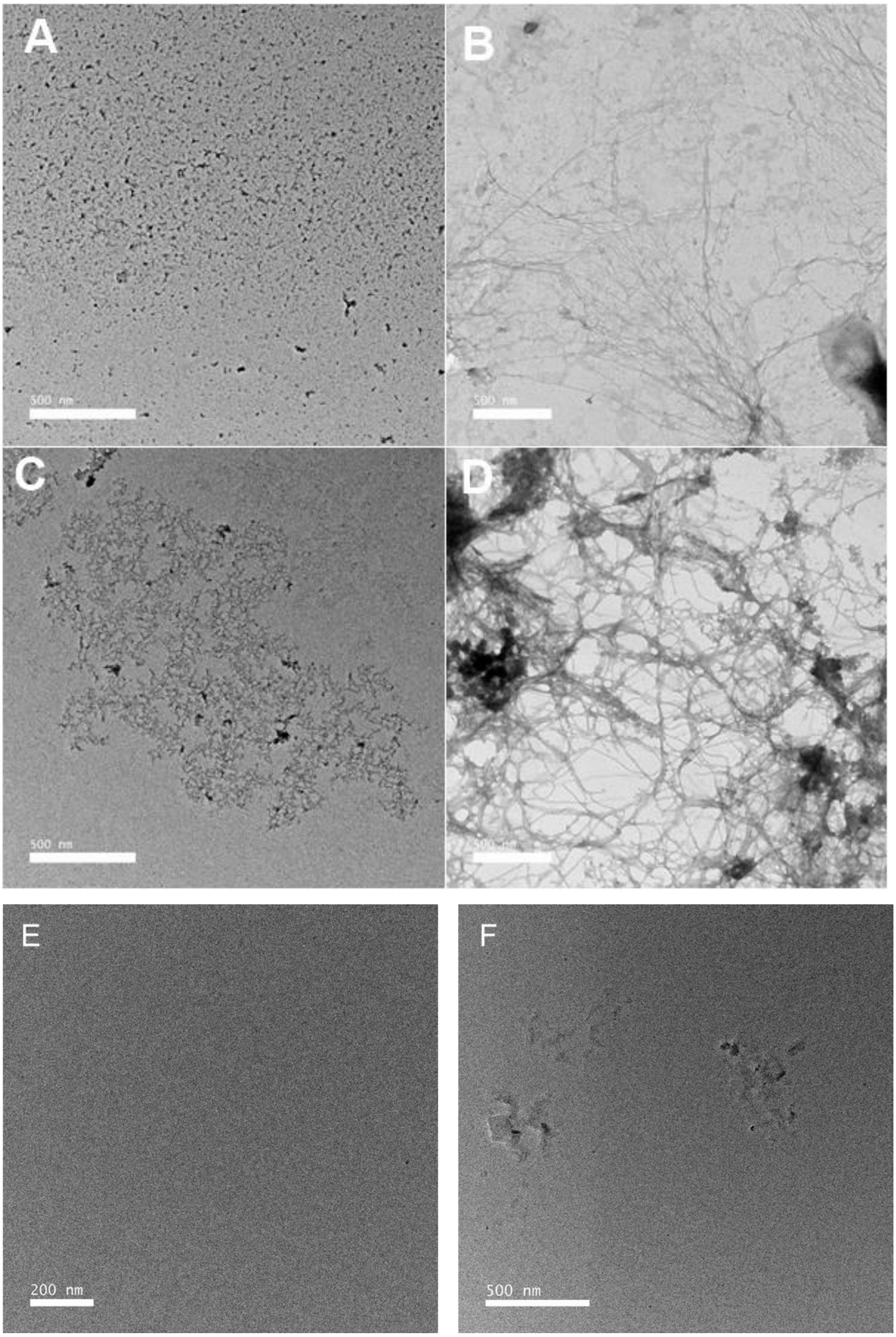

**Figure S9**. A) Tau (10 µM) without heparin; B) Tau (10 µM) with heparin (0.1 µM); C) Tau (10 µM) and **2** (1 µM); D) Tau (10 µM) and **1 (**1 µM); E) Compound **1** at 1 µM; F) Compound **2** at 1 µM.

### Cell viability assay of compound 1

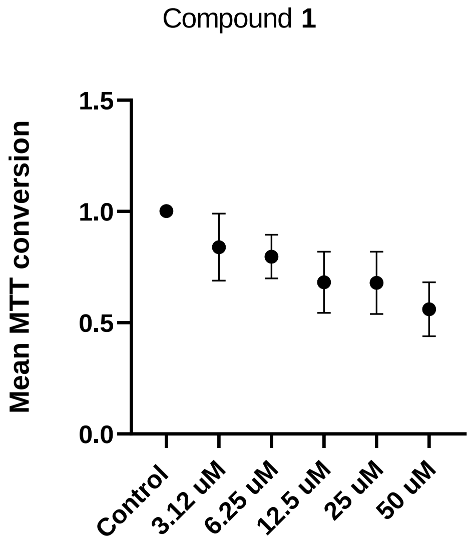

**Figure S10**: MTT assay on compound **1** at different concentrations (from 3.12 to 50 µM). The graph represents two independent plates with three replicates per compound concentration. Error bars indicate mean values ± SEM. Each concentration: n = 6; Positive control: n = 8; Negative control: n = 10. As control, DMSO 0.6% is used.

### FDAP analysis of 1 on PC-12 cells transfected with PAGFP-Tau441 ΔK280

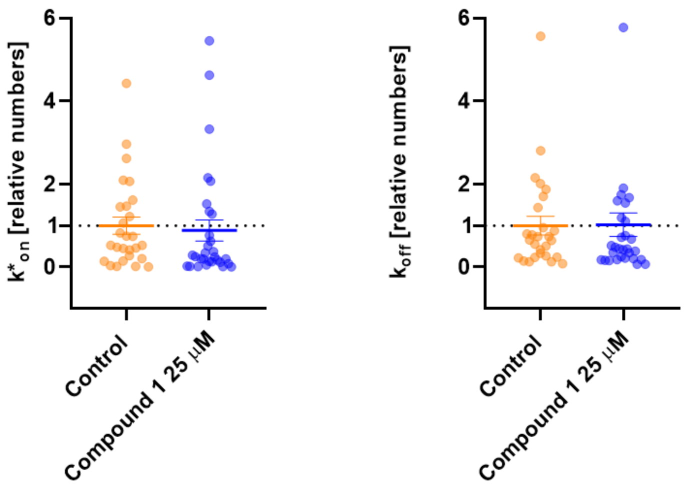

**Figure S11**. FDAP data were fitted with a mathematical model, whose outcome allowed the analysis of Tau $K_{on}$ and $K_{off}$. Experiments were performed with differentiated model neurons (PC12) cells differentiated with NGF), pre-incubation time was 24 h, DMSO concentration 0.125% (Control).

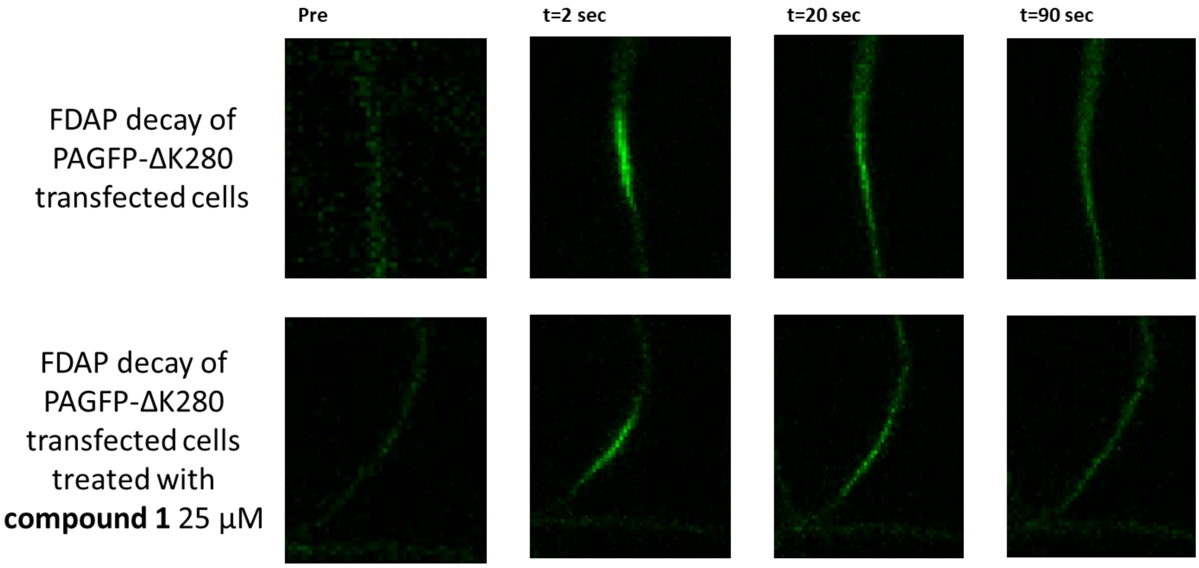

**Figure S12**. FDAP decay images of PAGFP-ΔK280 transfected cells at 0, 2, 20 and 90 sec in the control (up) and in the presence of compound **1** (down).

### Circular dichroism and infra-red spectroscopy of compound 3

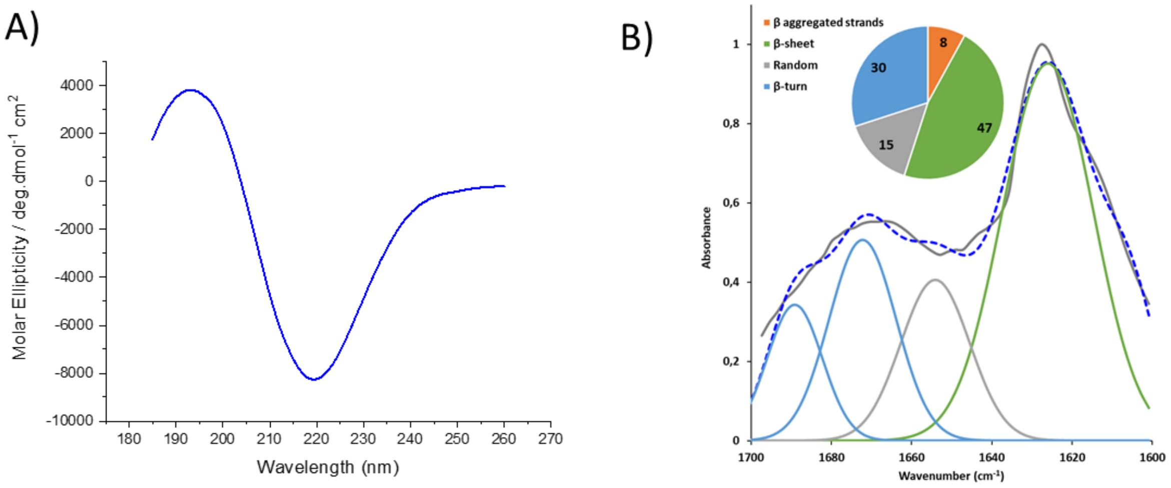

**Figure S13:** A) CD spectrum of compound **3** in 20 mM PB pH 7.2 at 125 µM concentration; B) IR-ATR amide I deconvolution of **3** (error squared = 0.068, standard deviation 7.74) in PB (125 µM, pH 7.2) with schematic representation of their secondary structure contents.

### Transmission Electron Microscopy (TEM) images of control for compound 3

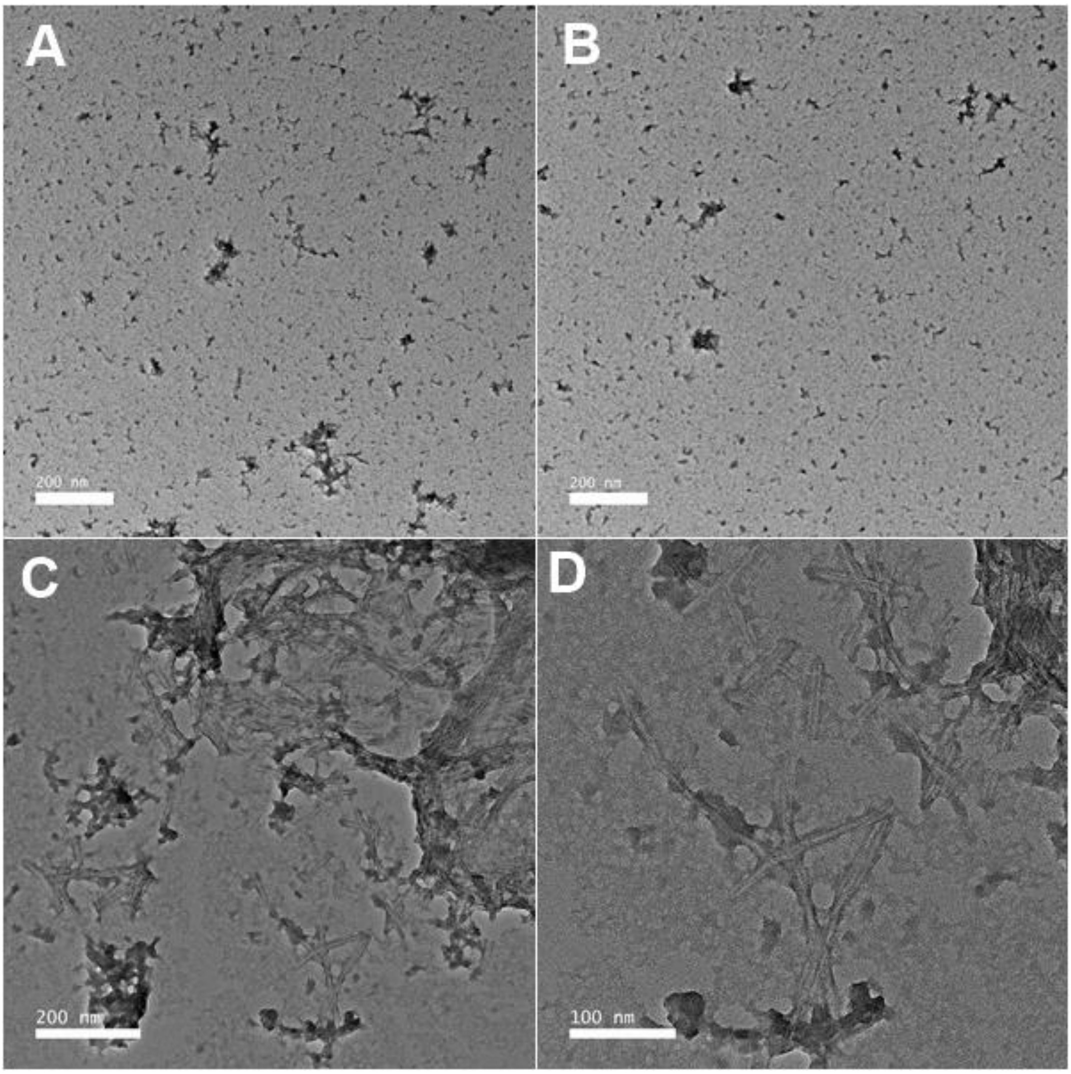

**Figure S14:** A) Tau (10 µM) and **3** (50 µM) B) Tau (10 µM) and **3** (10 µM) C-D) Tau (10 µM) and **3** (1 µM)

### FDAP analysis of 3 on PC-12 cells transfected with PAGFP-Tau441 ΔK280

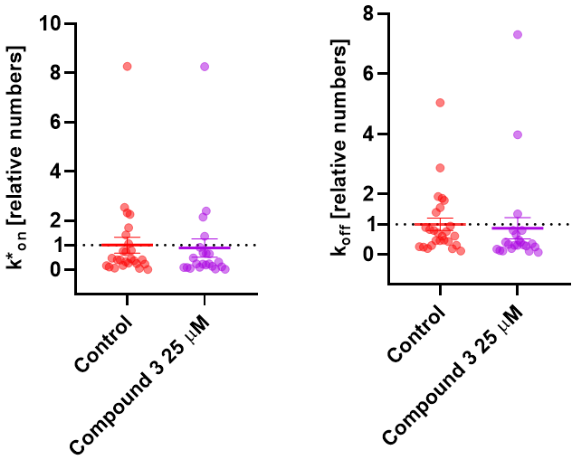

**Figure S15:** The fluorescence decay is fitted with a mathematical model, whose outcome allow to analyze Tau $K_{on}$ and $K_{off}$. Experiments were performed with differentiated model neurons (PC12) cells differentiated with NGF), compound incubation time was 24 h, DMSO concentration 0.125% (Control).

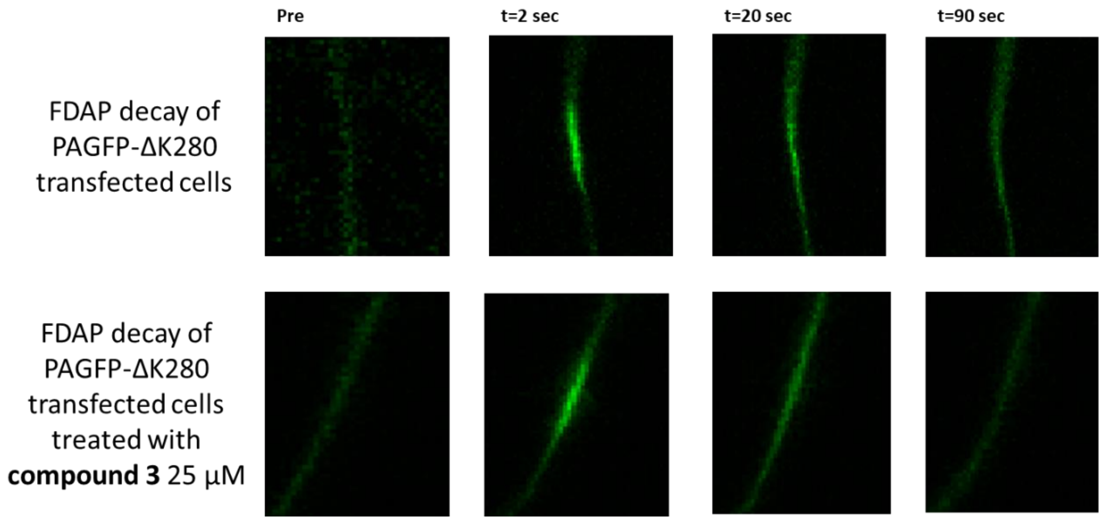

**Figure S16**. FDAP decay images of PAGFP-ΔK280 transfected cells at 0, 2, 20 and 90 sec in the control (up) and in the presence of compound **3** (down).
